# Drag-and-drop genome insertion without DNA cleavage with CRISPR-directed integrases

**DOI:** 10.1101/2021.11.01.466786

**Authors:** Eleonora I. Ioannidi, Matthew T. N. Yarnall, Cian Schmitt-Ulms, Rohan N. Krajeski, Justin Lim, Lukas Villiger, Wenyuan Zhou, Kaiyi Jiang, Nathaniel Roberts, Liyang Zhang, Christopher A. Vakulskas, John A. Walker, Anastasia P. Kadina, Adrianna E. Zepeda, Kevin Holden, Jonathan S. Gootenberg, Omar O. Abudayyeh

## Abstract

Programmable and multiplexed genome integration of large, diverse DNA cargo independent of DNA repair remains an unsolved challenge of genome editing. Current gene integration approaches require double-strand breaks that evoke DNA damage responses and rely on repair pathways that are inactive in terminally differentiated cells. Furthermore, CRISPR-based approaches that bypass double stranded breaks, such as Prime editing, are limited to modification or insertion of short sequences. We present Programmable Addition via Site-specific Targeting Elements, or PASTE, which achieves efficient and versatile gene integration at diverse loci by directing insertion with a CRISPR-Cas9 nickase fused to both a reverse transcriptase and serine integrase. Without generating double stranded breaks, we demonstrate integration of sequences as large as ∼36 kb with rates between 10-50% at multiple genomic loci across three human cell lines, primary T cells, and quiescent non-dividing primary human hepatocytes. To further improve PASTE, we discover thousands of novel serine integrases and cognate attachment sites from metagenomes and engineer active orthologs for high-efficiency integration using PASTE. We apply PASTE to fluorescent tagging of proteins, integration of therapeutically relevant genes, and production and secretion of transgenes. Leveraging the orthogonality of serine integrases, we engineer PASTE for multiplexed gene integration, simultaneously integrating three different genes at three genomic loci. PASTE has editing efficiencies comparable to or better than those of homology directed repair or non-homologous end joining based integration, with activity in non-dividing cells and fewer detectable off-target events. For therapeutic applications, PASTE can be delivered as mRNA with synthetically modified guides to programmably direct insertion of DNA templates carried by AAV or adenoviral vectors. PASTE expands the capabilities of genome editing via drag-and-drop gene integration, offering a platform with wide applicability for research, cell engineering, and gene therapy.

**One Sentence Summary:** A new technology combining CRISPR-mediated genome editing and site-specific integrases enables efficient programmable gene integration at any targeted genomic locus without double-strand DNA breaks, leading to broad applications in basic science research, cell engineering, and gene therapy.

## Main Text

Programmable genome insertion is integral to both gene therapy and basic research. Existing methods to insert long DNA sequences are either inefficient or rely on cellular mechanisms for double-strand break (DSB) repair. These technologies require programmable nucleases, such as CRISPR-Cas9, for (*1–3*) induction of repair pathways such as non-homologous end joining (NHEJ) (*4*) — as with the homology-independent targeted insertion (HITI) (*5*) technology — or homology directed repair (HDR) (*6–9*). However, DSB-based approaches are limited, as genome damage causes undesirable outcomes, including insertions/deletions, translocations, and activation of p53 (*10, 11*); NHEJ can generate off-target insertions at unintended DSBs (*12*); HDR has low efficiency in non-dividing cells and many cell types *in vivo*; and HDR requires long DNA templates that are labor-intensive to produce (*13*). Newer genome editing technologies such as base editing (*14–16*) and prime editing (*17*) avoid creating DSBs, but these technologies cannot install all genome edits. Base editing is limited to specific nucleotide transition changes (C→T, G→A, A→G, and T→C) and prime editing, based on a fusion of Cas9 from *Streptococcus pyogenes* (SpCas9) to an engineered Moloney Murine Leukemia Virus (M-MLV) reverse transcriptase, generates only nucleotide edits or small insertions (less than ∼50 nucleotides) or deletions (less than ∼80 nucleotides) (*17*).

Natural transposable element systems, which include several families of integrases and transposases, provide efficient routes for genome integration, but lack the programmability of CRISPR effector nucleases. Transposases insert varying copies of a donor sequence into cells at loosely defined sites, such as TA dinucleotides, resulting in semi-random gene insertion throughout the genome (*18*). In contrast, site-specific integrases, such as large serine phage integrases, efficiently integrate their DNA cargo into sequence-defined landing sites that are ∼30-50 nucleotides long (*19*) and have be used to insert therapeutic transgenes at naturally occurring pseudo-sites in the human genome in pre-clinical models (*20*). Targeted integration can also be achieved by a two-step approach involving prior insertion of integrase landing sites at a desired location using HDR (*21*). However, the inefficiency of two-step integration with HDR and the risks associated with DSBs have limited this approach in mammalian cells. Furthermore, a major issue limiting clinical application of certain integrases, such as phiC31, is that chromosomal rearrangements between pseudo-sites can occur, leading to a significant DNA damage response (*22, 23*).

Engineered systems to direct integrases, recombinases, or transposases to genomic sites for integration of gene cargos without DNA cleavage rely on fusions with programmable DNA binding proteins. Approaches fusing either zinc finger, transcription activator-like effector (TALE), or catalytically inactive Cas9 programmable DNA binding proteins to transposases (*24–28*) or recombinases (*29–35*) have been reported in mammalian cells, but their integration efficiency is low at genomic loci. Moreover, transposase fusions are hindered by excessive promiscuity and off-target insertions, while recombinase fusions have limited targets in the genome due to intrinsic sequence restrictions. As an alternative to engineered systems, natural CRISPR-associated Tn7-like transposons (*36*), either recruited by the Type V-K effector protein, Cas12k (*37*), or a Type I-F CRISPR–Cas system (*38*) have demonstrated high activity in bacterial cells. Despite this promise, these systems have not yet been successfully applied in eukaryotic contexts.

To overcome the current limitations of gene integration approaches, we married advances in programmable CRISPR-based gene editing with site-specific integrases to overcome dependence on DNA cleavage. By fusing Cas9, reverse transcriptases, and large serine integrases, we are able to programmably integrate cargos of up to ∼36 kb in a single-delivery reaction with efficiencies up to ∼55% in a diversity of cell types, including primary human hepatocytes and T cells. This approach, termed PASTE (Programmable Addition via Site-specific Targeting Elements), is easily retargeted to new genes, able to be delivered with a single dose of plasmids, and functional in non-dividing and primary cells. By profiling thousands of guide designs in a pooled screen, we determine guide rules for optimal programming to loci. We further engineered orthogonal integration with PASTE, simultaneously introducing three genes at three separate loci by tuning the central dinucleotide of the associated landing site. Through genome wide sequencing, we show that PASTE is much more specific than HITI, with higher purity of insertion than HDR and HITI. To improve upon PASTE, we mine bacterial genomes and metagenomes for novel integrases and apply them to high efficiency integration as part of the PASTE system. Moreover, we show diverse templates are compatible with PASTE, including AAV and adenovirus, allowing for drag-and-drop DNA integration of viruses and other DNA templates, a feature important for therapeutic applications. As a novel integration tool, PASTE opens multiple applications for gene insertion and tagging in biomedical research and therapeutic development.

## Results

### PASTE combines CRISPR-based genome editing and site-specific integration

We envisioned a programmable integration system coupling a CRISPR-based targeting approach with efficient insertion via serine integrases, which typically insert sequences containing an AttP attachment site into a target containing the related AttB attachment site. By using programmable genome editing to place integrase landing sites at desired locations in the genome, this system would guide the direct activity of the associated integrase to the specific genomic site. As prime editors have been reported to insert 44 bp sequences (*17*), we hypothesized that the ∼46 bp AttB landing site of serine integrases could be incorporated into the prime editing guide RNA (pegRNA) design and be copied into the genome via reverse transcription and flap repair (Fig. 1a-b). This “beacon” would serve as a target for an integrase, which could either be supplied in *trans* or directly fused to the Cas9 protein for additional recruitment. By simultaneously delivering a circular double-strand DNA template containing the AttP attachment site, we hypothesized that the DNA cargo could be directly integrated at the desired target site in a single-delivery reaction (Fig. 1a, b).

**Figure 1:**
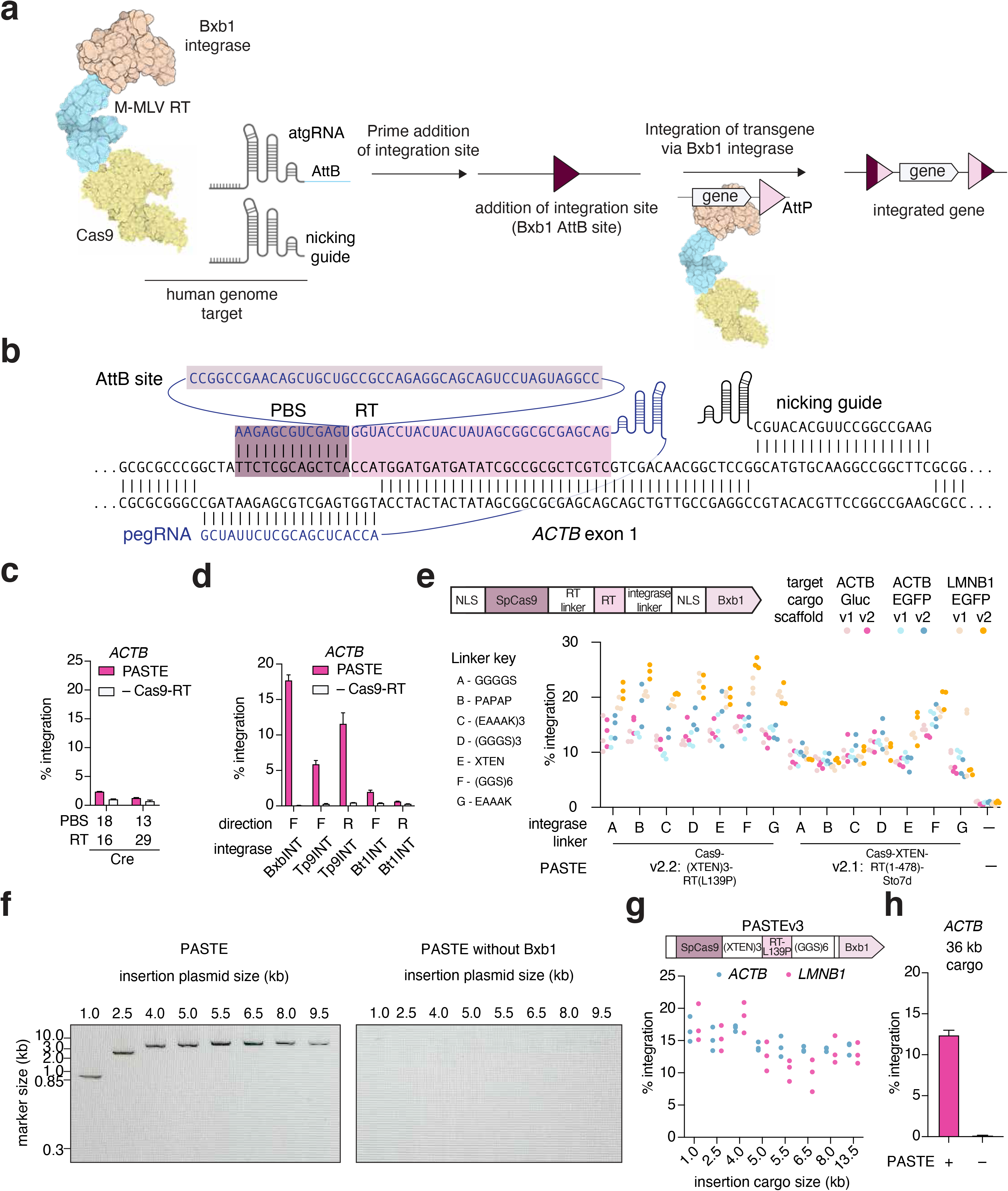
PASTE editing allows for programmable gene insertion independent of DNA repair pathways. a) Schematic of programmable gene insertion with PASTE. The PASTE system involves insertion of landing sites via Cas9-directed reverse transcriptases, followed by landing site recognition and integration of cargo via Cas9-directed integrases. b) Schematic of PASTE insertion at the *ACTB* locus, showing guide and target sequences. c) Comparison of *GFP* cargo integration efficiency between BxbINT and Cre recombinase at the 5’ end of the *ACTB* locus. d) Comparison of PASTE integration efficiency of GFP with a panel of integrases targeting the 5’ end of the *ACTB* locus. Both orientations of landing sites are profiled (F, forward; R, reverse). e) Optimization of PASTE constructs with a panel of linkers and RT modifications for EGFP integration at the *ACTB* and *LMNB1* loci with different payloads. f) Gel electrophoresis showing complete insertion by PASTE for multiple cargo sizes. g) Effect of cargo size on PASTE insertion efficiency at the endogenous *ACTB* and *LMNB1* targets. Cargos were transfected with fixed molar amounts. h) PASTE insertion of 36 kb cargo template at the *ACTB* locus. Data are mean (n= 3) ± s.e.m.

First, we assayed pegRNAs containing different AttB length truncations, hereafter referred to as attachment site-containing guide RNA (atgRNA) and we could insert sequences up to 56 bp at the beta-actin (*ACTB*) gene locus, with higher efficiency at lengths below 31 bp (Ext. Data Fig. S1a-b). Given that prime has been reported to insert LoxP sites to serve as beacons for Cre-based insertion (*17*), we tested a Cre-based integration approach with co-expression of PE2 and a Cre recombinase; however, tyrosine recombinases showed inefficient insertion (Fig 1c). Given the high efficiency of serine recombinases (*39*), we evaluated a panel of multiple enzymes, including Bxb1 (hereafter referred to as BxbINT), TP901 (hereafter referred to as (Tp9INT), and phiBT1 (hereafter referred to as Bt1INT) phage serine integrases (Supplementary Table 1), and we could insert all landing sites tested, with efficiencies between 10-30% (Ext. Data Fig. 1c). To test the complete system, we combined all components and delivered them in a single transfection: the prime editing vector, the atgRNA, a nicking guide for stimulating repair of the other strand, a mammalian expression vector for the corresponding integrase or recombinase and a 969 bp minicircle (*40*) DNA cargo encoding green fluorescent protein (*GFP*) (Fig. 1d). We compared *GFP* integration rates among the four integrases and recombinases and found that BxbINT integrase showed the highest integration rate (∼15%) at the targeted *ACTB* locus and required the nicking guide for optimal performance (Fig. 1d; Ext. Data Fig. 1d; 2a-b). This combined system, termed PASTEv1, resulted in programmable efficient insertion of the EGFP transgene.

We next hypothesized that we could improve PASTE editing through a series of protein and guide engineering efforts. We tested modified scaffold designs (atgRNAv2) for increased stabilization and expression from PolIII promoters (*41*), improving both atgRNA landing site insertion and overall PASTE efficiency (Ext. Data Fig. 2c). To optimize the other potential bottlenecks for PASTE’s complex activity, we screened a panel of protein modifications, including alternative reverse transcriptase fusions and mutations; various linkers between the Cas9, reverse transcriptase, and integrase domains; and reverse transcriptase and BxbINT domain mutants (Fig. 1e and Ext. Data Fig. 2c-f, Supplementary tables 2 and 3). A number of protein modifications, including an 48 residue XTEN linker between the Cas9 and reverse transcriptase, as well as the fusion of MMuLV to the Sto7d DNA binding domain or mutation of L139P (*42*) improved editing efficiency (Ext. Data Fig. 2d-f). When these top modifications were combined with a (GGGGS)_3_ linker between the reverse transcriptase- and BxbINT, they produced up to ∼30% gene integration, highlighting the importance of directly recruiting the integrase to the target site (Fig. 1e and Ext. Data Fig. 2h). We refer to this optimized construct, SpCas9-(XTEN-48)-RT(L139P)-(GGGGS)_3_-BxbINT, as PASTEv2. Further pairing of PASTEv2 with atgRNAv2 was termed PASTEv3, and was used for subsequent experiments. PASTEv3 achieved precise integration of templates as large as ∼36,000 bp with ∼10-20% integration efficiency at the *ACTB* and *LMNB1* loci (Fig. 1f-h and Ext. Data Fig. 3a-e), with complete integration of the full-length cargo confirmed by Sanger sequencing (Ext. Data Fig. 3f-g).

### atgRNA and integration site parameters influence PASTE efficiency

To optimize PASTEv3, we explored the impact of atgRNA and integrase parameters on integration efficiency. Relevant atgRNA parameters for PASTE include the primer binding site (PBS), reverse transcription template (RT), and AttB site lengths, as well as the relative locations and efficacy of the atgRNA spacer and nicking guide (Ext. Data Fig. 4a). We tested a range of PBS and RT lengths at two loci, *ACTB* and lamin B1 (*LMNB1*), and found that rules governing efficiency varied between loci, with shorter PBS lengths and longer RT designs having higher editing at the *ACTB* locus (Ext. Data Fig. 4b) and longer PBS and shorter RT designs performing better at *LMNB1* (Ext. Data Fig. 4c). These differences may be related to locus-dependent efficiency of priming and resolution of flap insertion observed in other prime editing applications (*17*). The length of the AttB landing site must balance two conflicting factors: the higher efficiency of prime editing for smaller inserts (*17*) and reduced efficiency of Bxb1 integration at shorter AttB lengths (*43*). We evaluated AttB lengths at *ACTB, LMNB1,* and nucleolar phosphoprotein p130 *(NOLC1)*, finding that the optimal AttB length was locus dependent. At the *ACTB* locus, long AttB lengths could be inserted (Fig. S1a) and overall PASTE efficiencies for the insertion of *GFP* were highest for long AttB lengths (Ext. Data Fig. 4d). In contrast, intermediate AttB lengths had higher overall integration efficiencies (> 20%) at *LMNB1* (Ext. Data Fig. S4e) and *NOLC1* (Ext. Data Fig. S4f), indicating that the increased efficiency of installing shorter AttB sequences overcame the reduction of BxbINT integration at these sites. We tested a panel of shorter RT and PBS guides at *ACTB* and *LMNB1* loci in comparison to our previous optimized guides and found that while shorter RT and PBS sequences did not increase editing at *ACTB* (Ext. Data Fig. 4g), they had improved editing at *LMNB1* (Ext. Data Fig. 4h) Moreover, manual design of a variety of atgRNA to different targets had varying levels of performance and editing outcomes at seven different gene loci (*ACTB*, *SUPT16H*, *SRRM2*, *NOLC1*, *DEPDC4*, *NES*, and *LMNB1*) (Ext. Data Fig. 4i).

To develop thorough rules for design, we tested atgRNA designs in high-throughput via pooled library screening (Fig. 2a). Using pooled oligo synthesis and cloning, we generated a library of 10,580 atgRNA designs for 11 spacers across 8 genes (ACTB, LMNB1, NOLC1, SUPT16H, DEPDC4, NES, CFTR, and SERPINA1). For each spacer/target pair, we were able to evaluate PBS lengths between 5-19 bp, RT lengths between 6-36 bp (increments of 2 bases), and AttB lengths of 38, 40, 43, and 46 bp, generating a distribution of edits across each target (Fig. 2b and Extended Data 1). Across the screen, every gene had atgRNAs with significant editing (Fig 2b-c). Upon analyzing the results, we found that more editing was generally found at a per target basis for shorter AttBs and that a wider range of RT and PBS lengths were permissible, although the exact optimal combinations differed across genes (Fig. 2d and Ext. Data Fig. 5). Across the 12 targets, RTs longer than 20 bp tended to yield higher editing, whereas PBS lengths could be between 5-19 bp without any clear trend. To validate the screen, we tested a panel of top predicted atgRNAs and found that they were all capable of higher efficiency AttB insertion (Fig. 2e) and PASTE integration (Fig. 2f) than manually designed atgRNAs derived from our arrayed screening of parameters.

**Figure 2:**
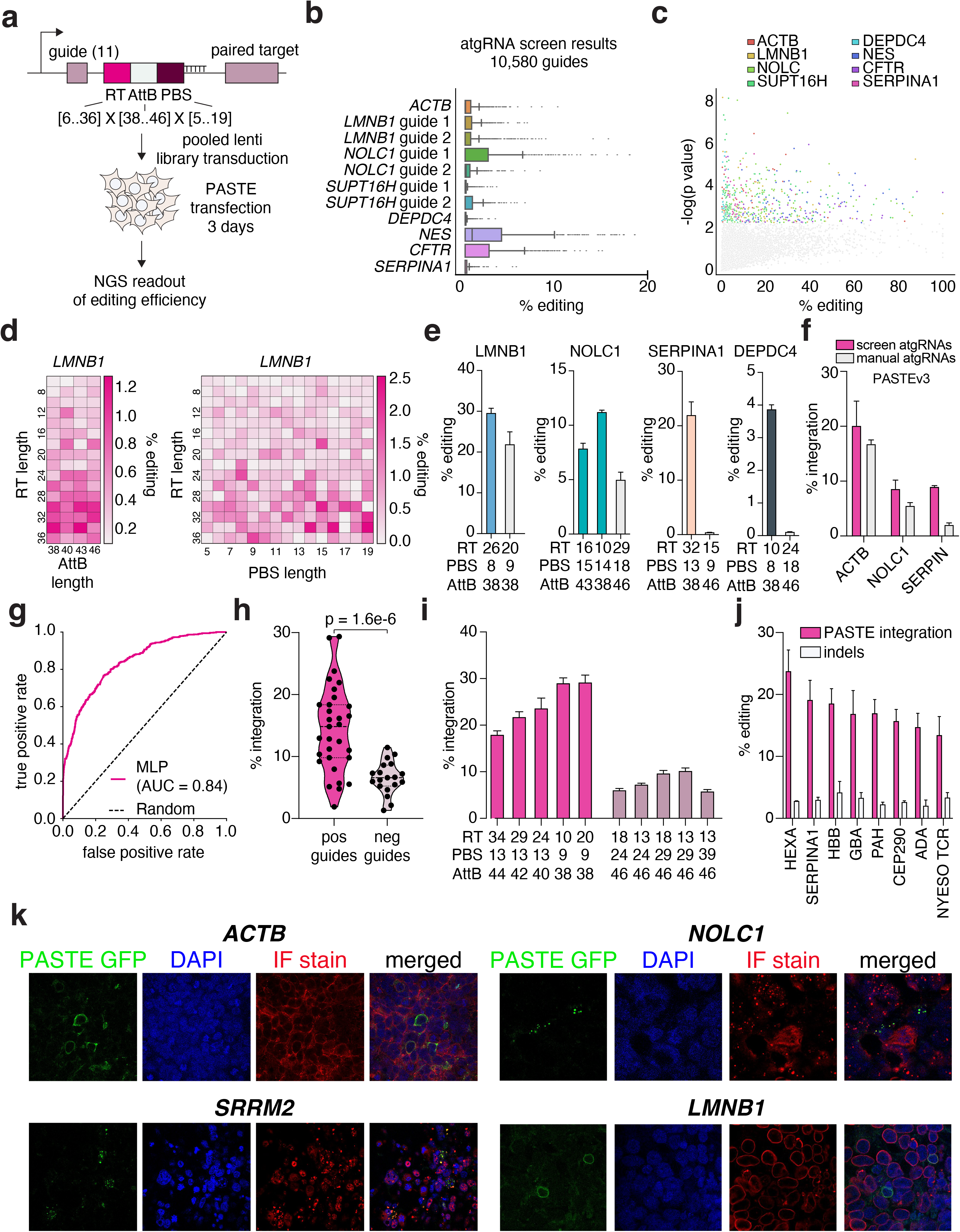
Evaluating design rules for efficient PASTE insertion at endogenous genomic loci. a) Schematic of pooled oligo library design for high-throughput screening of atgRNA designs at endogenous gene targets. b) Box plots depicting the editing rates of AttB addition at the different endogenous targets across 10,580 different atgRNA designs. Box indicates between 25th and 75th percentiles, whiskers indicate 1.5 times interquartile-range. c) Scatter plot depicting AttB site editing versus significance of the editing (–log(p-value)) as measured by a Student’s T-test against a no Cas9-RT control. d) Heatmaps depicting percent AttB site editing for LMNB1 guide 1 across different RT, PBS, and AttB lengths. e) Top atgRNA hits from the screen are compared for AttB site addition against manually designed atgRNAs (grey bars). f) PASTEv3 efficiency for insertion of an EGFP cargo at different endogenous targets is compared between screen validated atgRNAs and manually designed atgRNAs. g) Accuracy results by 5-fold cross validation of a MLP classifier trained on data from the 10,580 atgRNAs. h) PASTE integration rates of previously evaluated atgRNAs predicted by the MLP classifier to be efficient (pos. guides) or not efficient (neg. guides). i) PASTE integration rates of top atgRNAs predicted to be efficient (dark pink) or not efficient (light pink) by the MLP classifier. Solid line indicates median, dotted lines indicate 25th and 75th percentiles. j) PASTE integration rates and indel formation for integration of eight therapeutically relevant payloads at the *ACTB* locus. k) Endogenous protein tagging with *GFP* via PASTE by in-frame endogenous gene tagging at four loci (*ACTB*, *SRRM2*, *NOLC1*, and *LMNB1*). Immunofluorescence images of representative cells are shown. Cells have nuclear DAPI staining and antibody staining of the labeled proteins to show correlation to the endogenous PASTE tagging signal. Data are mean (n= 3) ± s.e.m.

To build an explicit predictive model for designing atgRNAs for PASTEv3, we trained a classifier using a kmer-based multi-layer perceptron for modeling the effect of an atgRNA sequence on the final editing rate of AttB insertion. Feature optimization and model training, had high accuracy (AUC = .84, Fig. 2g) and scoring of atgRNAs not seen by the model against *LMNB1*, *NOLC1*, and *ACTB* revealed clear differences in efficiency between guides nominated by the model and those rejected (Fig. 2h-i). Because our screening results have shown that rational design rules are difficult to generalize across gene targets, we release this prediction model as a guide design tool via a software package (https://github.com/abugoot-lab/pegRNA_rank) that simply receives as input a user’s target sequence and produces a list of atgRNAs rank-ordered by the predicted efficiency score.

The PE3 version of prime editing combines PE2 and an additional nicking guide to bias resolution of the flap intermediate towards insertion. To test the importance of nicking guide selection on PASTE editing, we tested editing at *ACTB* and *LMNB1* loci with two nicking guide positions. Suboptimal nicking guide positions reduced PASTE efficiency up to 30% (Ext. Data Fig. 6a-b) in agreement with the 75% reduction of PASTE efficiency in the absence of nicking guide (Ext. Data Fig. 1d, 6c). We also found, as expected, that the atgRNA spacer sequence was necessary for PASTE editing, and substitution of the spacer sequence with a non-targeting guide eliminated editing (Ext. Data Fig. 6d).

### PASTE is efficient at tagging multiple endogenous genes

Because PASTE does not require homology or sequence similarity on cargo plasmids, integration of diverse cargo sequences is modular and easily scaled across different loci. We tested seven different gene cargos: consisting of the common therapeutic genes *CEP290, HBB, PAH, GBA, ADA, SERPINA1,* and the NYESO T-cell receptor at the *ACTB* locus with PASTEv3, and a subset of these cargos at *LMNB1*. These cargos, which varied in size from 969 bp to 4906 bp, had integration frequencies between 15% and 22% depending on the gene and insertion locus, with minimal indel formation (Fig. 2j and Ext. Data Fig. 6e). We next tested if PASTE could make precise insertions for in-frame protein tagging or expressing cargo without disruption of endogenous gene expression. As BxbINT leaves residual sequences in the genome (termed AttL and AttR) after cargo integration, we hypothesized that these genomic scars could serve as protein linkers. We positioned the frame of the AttR sequence through strategic placement of the AttP on the minicircle cargo, achieving a suitable protein linker, GGLSGQPPRSPSSGSSG. Using this linker, we tagged four genes (*ACTB*, *SRRM2*, *NOLC1*, and *LMNB1*) with *GFP* using PASTE. To assess correct gene tagging, we compared the subcellular location of GFP with the tagged gene product by immunofluorescence. For all four targeted loci, GFP co-localized with the tagged gene product as expected, indicating successful tagging (Fig. 2k).

### PASTE efficiencies exceed DSB-based insertion methods

To benchmark PASTE against other gene integration methods, we compared PASTE to DSB-dependent gene integration using either NHEJ (i.e. HITI) (*5*) or HDR (*6, 7*) pathways (Fig. 3a). PASTE had equivalent or better gene insertion efficiencies than HITI (Fig. 3b). On a panel of 7 different endogenous targets, PASTE exceeded HITI editing at 6 out of 7 genes, with similar efficiency for the 7th gene (Fig. 3c). Importantly, as DSB generation can lead to insertions or deletions (indels) as an alternative and undesired editing outcome, we assessed the indel frequency by next-generation sequencing, finding significantly fewer indels generated with PASTE than HITI in both HEK293FT and HepG2 cells (Fig. 5b and Ext. Data Fig. 7a), showcasing the high purity of gene integration outcomes with PASTE due to the lack of DSB formation. We also compared PASTE to previously validated HDR constructs at the N-terminus of ACTB and LMNB1 for EGFP tagging, finding that although PASTE had similar efficiency at the ACTB locus and lower efficiency at the LMNB1 locus, it generated significantly fewer indels than HDR (Fig. 3d). Notably, both HDR and HITI generate more indels than desired on-target integrations at the ACTB locus.

**Figure 3:**
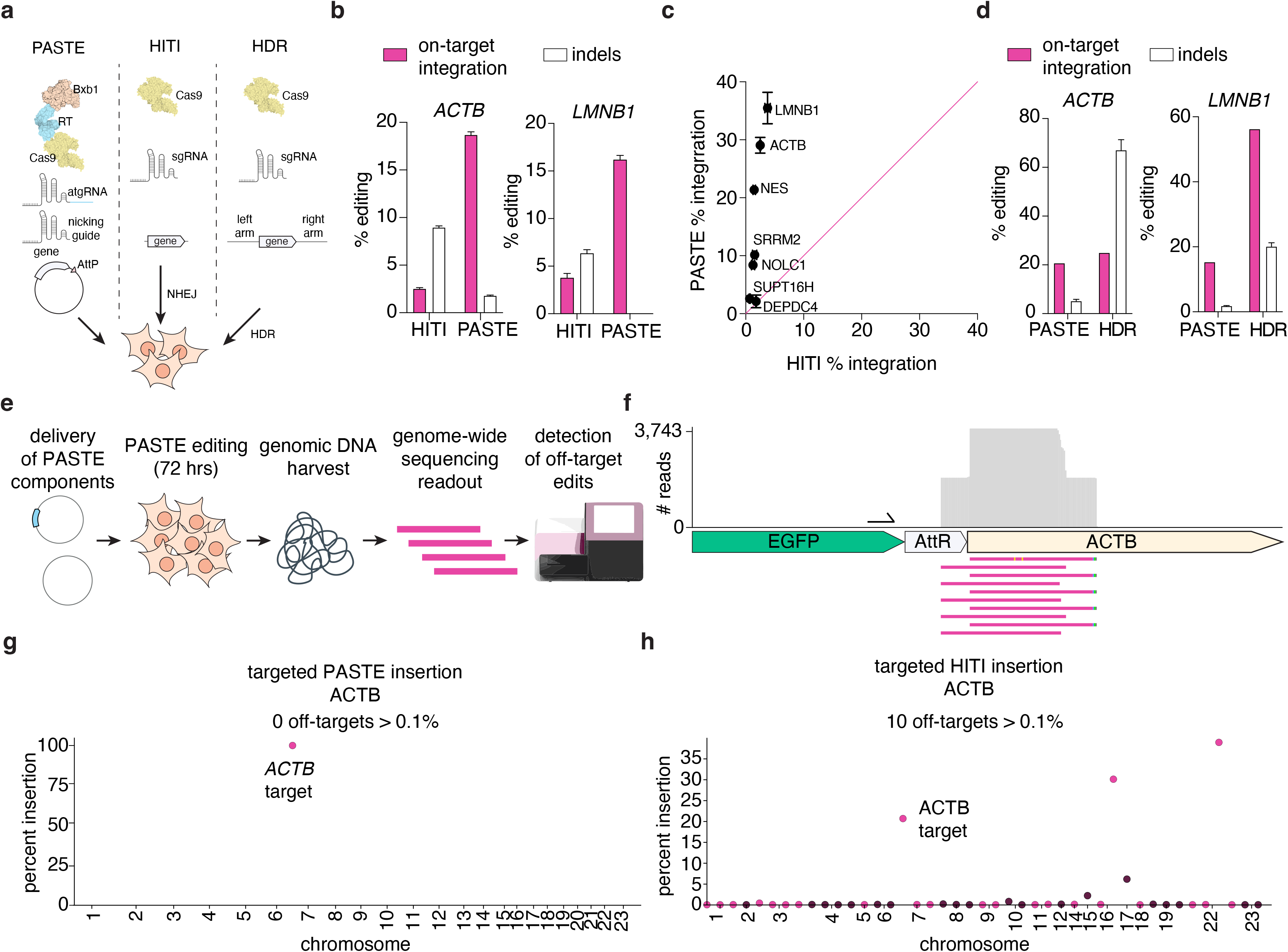
Characterization of genome-wide PASTE specificity and purity of integration compared to other integration approaches. a) Schematic of PASTE, HITI, and HDR gene integration approaches. b) Comparison of byproduct indel generation by PASTE and HITI at the *ACTB* and *LMNB1* target sites. Shown is the on-target EGFP integration rate observed compared to byproduct indels. c) *GFP* integration efficiency at a panel of genomic loci by PASTE compared to insertion rates by homology-independent targeted integration (HITI). d) Integration of a *GFP* template by PASTE at the *ACTB* and *LMNB1* loci compared to homology-directed repair (HDR) at the same target. Quantification is by single-cell clone counting. Integration efficiency is compared to the rate of byproduct indel generation. e) Schematic of next-generation sequencing method to assay genome-wide off-target integration sites by PASTE and HITI. f) Alignment of reads at the on-target *ACTB* site using our unbiased genome-wide integration assay, showing expected on-target PASTE integration outcomes. g) Manhattan plot of averaged integration events for multiple single-cell clones with PASTE editing. The on-target site is at the *ACTB* gene on chromosome 7 (labeled). Number of off-targets with greater than 0.1% integration is shown. h) Manhattan plot of averaged integration events for multiple single-cell clones with HITI editing. The on-target site is at the *ACTB* gene on chromosome 7 (labeled). Number of off-targets with greater than 0.1% integration is shown. Data are mean (n= 3) ± s.e.m.

**Figure 4:**
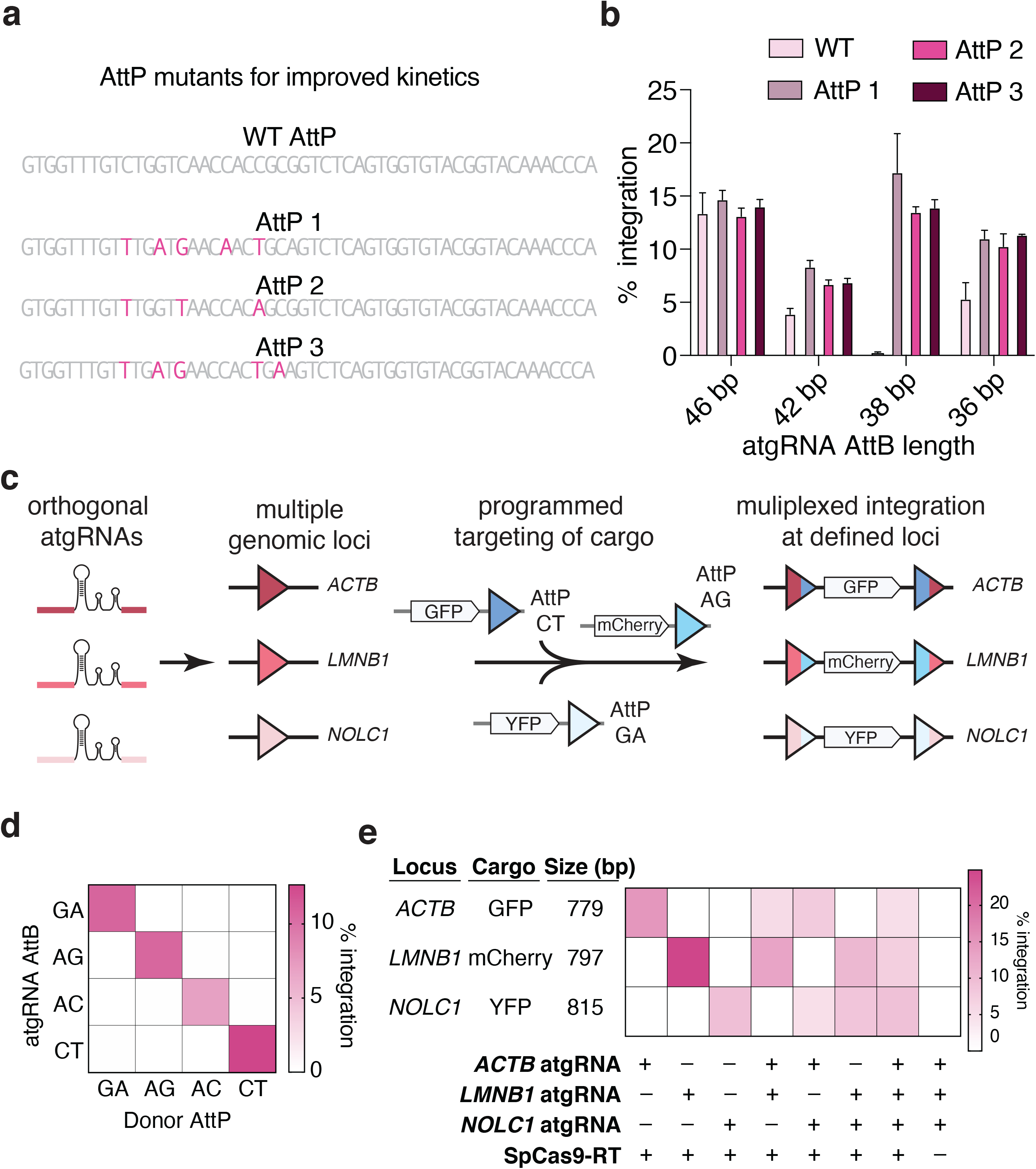
Multiplexed and orthogonal gene insertion with PASTE. a) Schematic of AttP mutations tested for improving integration efficiency. b) Integration efficiencies of wildtype and mutant AttP sites across a panel of AttB lengths. c) Schematic of multiplexed integration of different cargo sets at specific genomic loci. Three fluorescent cargos (GFP, mCherry, and YFP) are inserted orthogonally at three different loci (*ACTB, LMNB1, NOLC1*) for in-frame gene tagging. d) Orthogonality of top 4 AttB/AttP dinucleotide pairs evaluated for GFP integration with PASTE at the *ACTB* locus. e) Efficiency of multiplexed PASTE insertion of combinations of fluorophores at *ACTB, LMNB1,* and *NOLC1* loci. Data are mean (n= 3) ± s.e.m.

**Figure 5:**
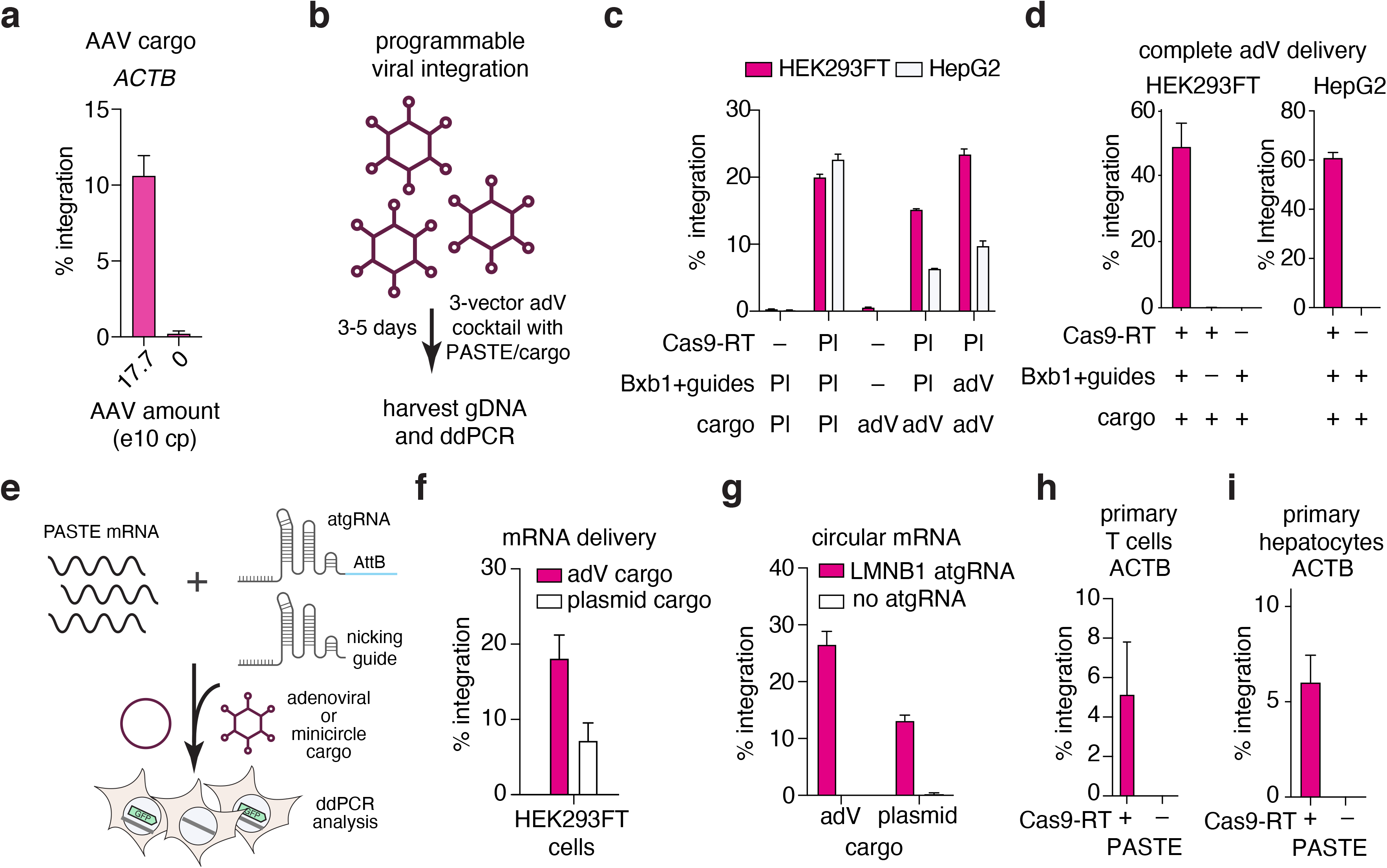
PASTE is compatible with multiple delivery methods and is active in primary cell types. a) Insertion templates delivered via AAV transduction. PASTE editing machinery was delivered via transfection, and templates were co-delivered via AAV dosing at levels indicated. b) Schematic of AdV delivery of the complete PASTE system with three viral vectors. c) Integration efficiency of AdV delivery of integrase, guides, and cargo in HEK293FT and HepG2 cells. BxbINT and guide RNAs or cargo were delivered either via plasmid transfection (Pl), AdV transduction (AdV), or omitted (-). SpCas9-RT was only delivered as plasmid or omitted. d) AdV delivery of all PASTE components in HEK293FT and HepG2 cells. e) Schematic of mRNA and synthetic guide delivery of PASTE components f) Delivery of PASTE system components with mRNA and synthetic guides, paired with either AdV or plasmid cargo. g) Delivery of circular mRNA with synthetic guides and either AdV or plasmid cargo. h) PASTE editing efficiency with single vector designs in primary human T cells. i) PASTE editing efficiency with single vector designs in primary human hepatocytes. Data are mean (n= 3) ± s.e.m.

### Off-target characterization of PASTE and HITI gene integration

As off-target editing is a critical consideration for novel genome editing technologies, we explored the specificity of PASTE at specific sites through two hypotheses: 1) off-targets generated by BxbINT integration into pseudo-AttB sites in the human genome and 2) off-targets generated via guide- and Cas9-dependent editing in the human genome. While BxbINT lacks documented integration into the human genome at pseudo-attachment sites (*44*), we computationally identified potential sites with partial similarity to the natural BxbINT AttB core sequence. We tested for BxbINT integration by ddPCR across these sites and found no off-target activity (Ext. Data Fig. 7b-g). To assay Cas9 off-targets for our *ACTB* atgRNA, we identified two potential off-target sites via computational prediction and found no off-target integration for PASTE (Ext. Data Fig. 7c), but substantial off-target activity by HITI at one of the sites (Ext. Data Fig. 7d). While PASTE is shown to be specific for our targets, Cas9-based off-target analysis should be performed for each new PASTE target to ensure specificity.

As computationally predicted sites may not account for all possible off-targets, we additionally evaluated genome-wide off-targets due to either Cas9 or BxbINT through tagging and PCR amplification of insert-genomic junctions (Fig. 3e). We isolated single cell clones for conditions with PASTE editing and negative controls missing PE2, and deep sequencing of insert-genomic junctions from these clones showed all reads aligning to the on-target *ACTB* site, confirming no off-target genomic insertions (Fig. 3ef and Ext. Data Fig. 7h). We also used this genome-wide pipeline to analyze HITI off-targets using the same ACTB guide and HITI EGFP insertion template and found substantial off-target activity across the genome with 42 different sites identified across 20 chromosomes and with 10 of these off-targets having greater than 0.1% integration. Moreover, the on-target ACTB edit was only 20.7% of the reads identified with two other off-targets having higher efficiency. These results show that linear template-based integration approaches have significant off-target activity and highlight the benefits of using circular templates with a dual-nicking PASTE system.

Expression of reverse transcriptases and integrases involved in PASTE may have detrimental effects on cellular health. To determine the extent of these effects, we transfected the complete PASTE system, the corresponding guides and cargo with only PE2, and the corresponding guides and cargo with only BxbINT, and compared them to both GFP control transfections and guides without protein expression via transcriptome-wide RNA sequencing. We found that, while BxbINT expression in the absence of Prime editing had several significant off-targets, the complete PASTE system had only one differentially regulated gene with more than a 1.5-fold change (Ext. Data Fig. 7i-j). Genes upregulated by BxbINT overexpression included stress response genes, such as TENT5C and DDIT3, but these changes were not seen in the expression of the PASTE system (Ext. Data Fig. 7i-j), potentially due to differences in expression when BxbINT is linked to the PASTE construct.

### AttB engineering for multiplexed gene integration and enhanced editing activity

To optimize PASTE efficiency, we profiled attachment site mutants for optimization of integration kinetics of BxbINT, especially for shorter AttB sites that have reduced integration efficiency. Testing a panel of different AttP sequences (Fig. 4a), including ones previously shown to affect BxbINT integration, we found AttP sequence variants that substantially improved the integration rate, especially for the 38 bp AttB at ACTB (Fig. 4b and Ext. Data Fig. 8a).

The central dinucleotide of BxbINT is intimately involved in the association of AttB and AttP sites for integration (*43*), and changing the matched central dinucleotide sequences can modify integrase activity and provide orthogonality for insertion of two genes (*45*). We hypothesized that expanding the set of AttB/AttP dinucleotides could enable multiplexed gene insertion with PASTE, using orthogonal atgRNA combinations (Fig 4c). To find optimal AttB/AttP dinucleotides for PASTE insertion, we profiled the efficiency of *GFP* integration at the *ACTB* locus with PASTE across all 16 dinucleotide AttB/AttP sequence pairs. We found several dinucleotides with integration efficiencies greater than the wild-type GT sequence (Ext. Data Fig 8b). Importantly, a majority of dinucleotides had 75% editing efficiency or greater compared to wild-type AttB/AttP efficiency, implying that these dinucleotides could be potential orthogonal channels for multiplexed gene insertion with PASTE.

Next, we explored the specificity of matched and unmatched AttB/AttP dinucleotide interactions. We comprehensively profiled the interactions between all dinucleotide combinations in a scalable fashion using a pooled assay to compare AttB/AttP integration (Ext. Data Fig. 8c). By barcoding 16 AttP dinucleotide plasmids with unique identifiers, co-transfecting this AttP pool with the BxbINT integrase expression vector and a single AttB dinucleotide acceptor plasmid, and sequencing the resulting integration products, we measured the relative integration efficiencies of all possible AttB/AttP pairs (Ext. Data Fig. 8d). We found that dinucleotide specificity varied wildly, with some dinucleotides (GG) exhibiting strong self-interaction with negligible crosstalk, and others (AA) showing minimal self-preference. Sequence logos of AttP preferences (Ext. Data Fig. 8e) reveal that dinucleotides with C or G in the first position have stronger preferences for AttB dinucleotide sequences with shared first bases, while other AttP dinucleotides, especially those with an A in the first position, have reduced specificity for the first AttB base. Informed by the efficiency and specificity of the central dinucleotides, we tested GA, AG, AC, and CT dinucleotide atgRNAs for *GFP* integration at *ACTB*, either paired with their corresponding AttP cargo or mis-paired with the other three dinucleotide AttP sequences. We found that all four of the tested dinucleotides efficiently integrated cargo only when paired with the corresponding AttB/AttP pair, with no detectable integration across mispaired combinations (Fig. 4d).

Selecting the three top dinucleotide attachment site pairs (CT, AG, and GA), we designed atgRNAs that target *ACTB* (CT), *LMNB1* (AG), and *NOLC1* (GA) and corresponding minicircle cargo containing GFP (CT), mCherry (AG), and YFP (GA). Upon co-delivering these reagents to cells, we found that we could achieve single-plex, dual-plex, and trip-plex editing of all possible combinations of these atgRNAs and cargo in the range of 5%-25% integration (Fig. 4e).

A useful application for multiplexed gene integration is for labeling different proteins to visualize intracellular localization and interactions within the same cell. We used PASTE to simultaneously tag *ACTB* (GFP) and *NOLC1* (mCherry) or *ACTB* (GFP) and *LMNB1* (mCherry) in the same cell. We observed that no overlap of GFP and mCherry fluorescence and confirmed that tagged genes were visible in their appropriate cellular compartments, based on the known subcellular localizations of the *ACTB*, *NOLC1* and *LMNB1* protein products (Ext. Data Fig. 8f).

### Directed programmable integration of viruses for therapeutic payload delivery and expression

To explore compatibility of PASTEv3 with therapeutically relevant delivery modalities, we explored whether components of the PASTE system could be delivered with either adenovirus-associated viral (AAV) or adenoviral (AdV) vectors. Testing AAV-delivered cargo with an AttP-containing payload in conjunction with other PASTE components delivered via transfection, we found above 10% integration of the viral payload in a dose dependent fashion (Fig 5a, Ext. Data Fig. S9a-b). This revealed that the AAV genome could serve as a suitable template for serine integrase-mediated insertion, agreeing with reports of AAV genome circularization in cells (*46*).

In order to package the complete PASTE system in viral vectors, we utilized an AdV vector, an emerging approach for clinical delivery of large cargo (*47*) (Fig. 5b). We first evaluated whether adenovirus could deliver a suitable template for BxbINT-mediated insertion along with plasmids for SpCas9-RT-BxbINT and guide expression, or AdV delivery of guides and BxbINT with plasmid delivery of SpCas9-RT, finding that we could achieve 10-20% integration of the ∼36 kb adenovirus genome carrying EGFP in HEK293FT and HepG2 cells (Fig. 5c and Ext. Data. Fig. 9c). Upon packaging and delivering the cargo and PASTE system components across 3 AdV vectors, we found that the complete PASTE system (Cas9-reverse transcriptase, integrase and guide RNAs, and cargo) could be substituted by adenoviral delivery, with integration of up to ∼50-60% with viral-only delivery in HEK293FT and HepG2 cells (Fig. 5d).

To further demonstrate PASTE would be amenable for in vivo delivery, we developed an mRNA version of the PASTE protein components as well as chemically-modified synthetic atgRNA and nicking guide against the LMNB1 target (Fig. 5e). Electroporation of the mRNA and guides along with delivery of the template via adenovirus or plasmid yielded high efficiency integration up to ∼23% (Fig. 5e-f and Ext. Data Fig. 9d). As we hypothesized more sustained BxbINT expression would allow for integration into newly placed AttB sites in the genome, we tested circular mRNA expression(*48*) and found that this boosted the efficiency of integration to ∼30% (Fig. 5g).

### PASTE efficiency in non-dividing and primary cells

As PASTE does not rely on DSB repair pathways that are only active in dividing cells, we tested PASTE activity in non-dividing cells by transfecting either Cas9 and HDR templates or PASTE into HEK293FT cells and arresting cell division (*49*) via aphidicolin treatment (Ext. Data Fig. 9e). In this model of blocked cell division, we found that PASTE maintained *GFP* gene integration activity greater than 20% at the *ACTB* locus whereas HDR-mediated integration was abolished (Ext. Data Fig. 9g). To evaluate the size limits for therapeutic transgenes, we evaluated insertion of cargos up to 13.3 kb in length in both dividing and aphidicolin treated cells, and found insertion efficiency greater than 10% (Ext. Data Fig. 9hf), enabling insertion of ∼99.7% of all full-length human cDNA transgenes (*50*). To overcome reduction of large insert delivery to cells due to potential delivery inefficiencies, we found that delivering larger DNA amounts of insert could significantly improve gene integration efficiency (Ext. Data Fig. 9i)

We also expanded PASTE editing to additional cell types, testing PASTE in the K562 lymphoblast line, primary human T cells, and primary human hepatocytes. We found that PASTE had ∼15% gene integration activity in K562 cells and around 5% efficiency in primary human T cells (Fig. 5h and Ext. Data Fig. 9j). In addition, in non-dividing quiescent human primary hepatocytes, we found that PASTE was capable of ∼5% gene integration at the ACTB locus (Fig. 5i), consistent with the non-dividing activity we observed with the aphidicolin-treated HEK293FT cells.

### Using PASTE for programmable gene integration results in production and secretion of therapeutic transgenes

Programmable gene integration provides a modality for expression of therapeutic protein products, and we tested protein production of therapeutically relevant proteins Alpha-1 antitrypsin (encoded by *SERPINA1)* and Carbamoyl phosphate synthetase I (encoded by *CPS1),* involved in the diseases Alpha-1 antitrypsin deficiency and CPS1 deficiency, respectively. By tagging gene products with the luminescent protein subunit HiBiT (*51*), we could independently assess transgene production and secretion in response to PASTE treatment (Ext. Data Fig. 10a). We transfected PASTE with *SERPINA1* or *CPS1* cargo in HEK293FT cells and a human hepatocellular carcinoma cell line (HepG2) and found efficient integration at the *ACTB* locus ( Ext. Data Fig. 10b-c). This integration resulted in robust protein expression, intracellular accumulation of transgene products), and secretion of proteins into the media (Ext. Data Fig. 10d-g).

### Discovery and development of novel integrases for PASTE integration

As we found that integrase choice can have implications for integration activity (Fig. 1c-d), we decided to mine bacterial and metagenomic sequences for new phage associated serine integrases (Fig. 6a). Exploring over 10 TB worth of data from NCBI, JGI, and other sources, we found 27,399 novel integrases (Fig. 6b-c, Ext. Data Fig 10h, and Extended Data 2) and annotated their associated attachment sites using a novel repeat finding algorithm that could predict potential 50 bp attachment sites with high confidence near phage boundaries. Analysis of the integrases sequences revealed that they fell into four distinct clusters: INTa, INTb, INTc, and INTd with diverse domain architectures (Fig. 6c). About half of integrases (14,771) derive from metagenomic sequences, presumably from pro-phages, and 13,693 of the integrases specifically derive from human microbiome metagenomic samples. An initial screen of integrase activity using a reporter system revealed that a number of the integrases were highly active in HEK293FT cells with more activity than BxbINT, a member of the INTa family (Fig. 6d). Using the predicted 50 bp sequences encoded in atgRNAs along with minicircles containing the complementary AttP sites, we found that these integrases were compatible with PASTE, but performed less effectively than BxbINTa-based PASTE (Fig. 6e). We hypothesized that this reduction in performance of the new integrases was due to their longer 50 bp AttB sequences and so we explored truncations of these AttBs in the hopes of finding more minimal attachment sites. Truncation screening on integrase reporters revealed that AttB truncations of all the integrases, including as short as 34 bp, were still active and many had more activity than BxbINTa (Fig. 6f). Upon porting these new shorter AttBs to atgRNAs for PASTE, we found that a number of integrases had more activity in the PASTE system than BxbINT-based PASTE at the ACTB locus, including the integrase from *B. cereus* (BceINTc), N191352_143_72 stool sample from China (SscINTd), and N684346_90_69 stool sample from adult in China (SacINTd), while others like the integrase from *B. cytotoxicus* (BcytINTd) and *S. lugdunensis* (SluINTd) did not (Fig. 6d,g-h). We next fused BceINTc to SpCas9-MLV-RT^L139P^, termed PASTEv4, and found that it performed better than BxbINTa-based PASTE across a number of endogenous gene loci (Fig. 6i and Ext. Data Fig. 10i).

**Figure 6:**
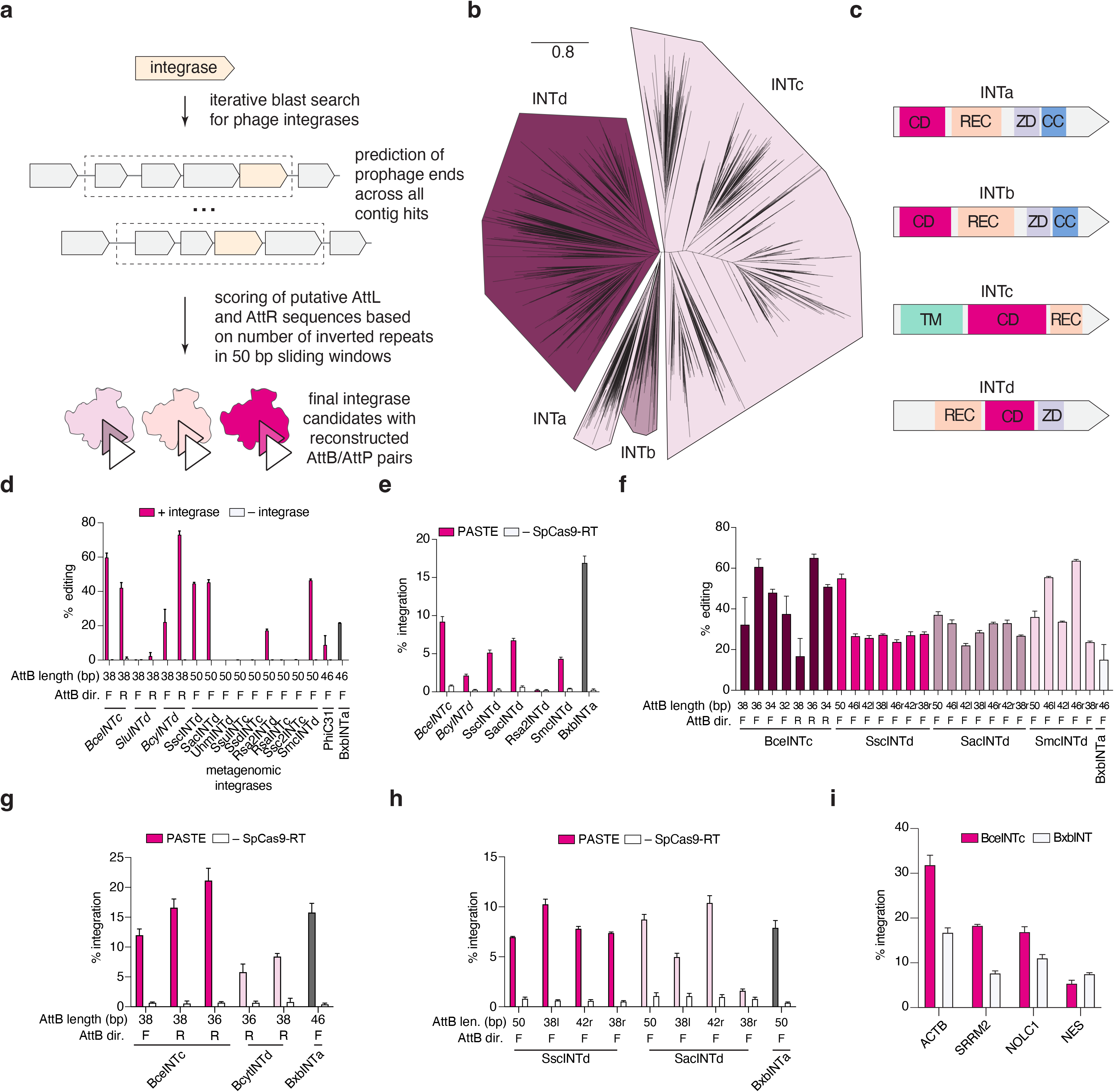
Discovery of novel phage-derived integrases for programmable gene integration with PASTE. a) Schematic of integrase discovery pipeline from bacterial and metagenomic sequences. b) Phylogenetic tree of discovered integrases showing distinct subfamilies. c) Domain architecture of the four integrase sub-families. d) Screening novel integrase integration activity using reporters in HEK293FT cells compared to BxbINT and phiC31. e) PASTE integration activity with most active integrases compared to BxbINT. f) Characterization of integrase integration activity with truncated attachment sites using reporters in HEK293FT cells. g) PASTE integration activity with BceINT and BcyINT with truncated attachment sites compared to BxbINT. h) PASTE integration activity with SscINT and SacINT with truncated attachment sites compared to BxbINT. j) Integration of EGFP at different endogenous gene targets for PASTE with either BceINT or BxbINT. Data are mean (n= 3) ± s.e.m.

## Discussion

We develop PASTE, via engineering of Cas9, reverse transcriptase, and integrase linkers to create a fusion protein capable of efficient integration (5-50%) of diverse cargos at precisely defined target locations within the human genome with small, stereotyped scars that can serve as protein linkers. We demonstrate the versatility of PASTE for gene tagging, gene replacement, gene delivery, and protein production and secretion. Through extensive characterization of integrase attachment sites, we engineer multiplexed gene integration with PASTE, enabling applications such as the specific fusion of three different endogenous genes with three different fluorescent cargos. Overall, we show PASTE insertions at 9 different endogenous sites with 20 different cargos ranging in size from 779 bp to 36,000 bp, which would enable insertion of greater than 99.7% of human cDNAs. Utilizing pooled screening approaches for testing thousands of guide hypotheses in parallel, we define rules for PBS, RT, AttB/AttP lengths, and sequence characteristics to maximize efficiency and specificity. In agreement with previous studies of serine integrases and prime editing, we find no off-target activity with PASTE. Furthermore, via metagenomic mining, we discover thousands of putative integrase/attachment site combinations, and engineer multiple novel integrase orthologs with improved activity and reduced attachment site requirements to further optimize the activity of PASTE, generating a PASTEv4 system using the BceINT integrase. As therapeutic use of certain integrases like phiC31 suffers from concerns of genome rearrangement due to pseudo-sites in the human genome (*22, 23*), use of novel integrases in PASTE offers greater clinical potential because of the absence of pseudo-sites in the human genome (*44*), resulting in no detectable off-target genomic integration. Moreover, in contrast to transposase-based integration systems (*37*), PASTE integration is stereotyped, allowing for precise design of integration and predictable gene fusions. As PASTE does not rely on HDR, it can function in non-dividing cells, including in primary hepatocytes and T-cells.

Programmable insertion is a fundamental tool for genetics, as tagging of gene products with fluorophores or degradation tags enables new modalities for measuring or perturbing specific genes of interest. Extension of these concepts to large scale insertion screens in diverse cells *in vitro* and *in vivo* is further facilitated by PASTE, as the ease of programmability is amenable to pooled screening, where multiple genes can be tagged simultaneously and evaluated for phenotypic or expression effects in throughput. Sequence insertion can also be used for integration of mutant genes to understand variants of unknown function, to develop models to explore mechanisms of disease, or to test potential therapeutic interventions. PASTE enables therapeutic correction of genetic disease through insertion of full length, functional genes at native loci, a viable strategy for both treating recessive loss of function mutations that cover 4,122 genetic diseases (*52*) and overcoming dominant negative mutations. Current genome editing approaches for diseases such as cystic fibrosis or leber’s congenital amaurosis (*53, 54*) are limited, as systems must be tailored for specific mutations (*55, 56*), requiring unique genome editing therapies for each subset of the patient population. Programmable insertion of the wild-type gene at the endogenous location would address all potential patient mutations, serving as a blanket therapy. Beyond direct correction of hereditary disease, gene insertion provides a promising avenue for cell therapies, and efficient integration of engineered transgenes, such as chimeric antigen receptors, at specific loci can produce significantly improved therapeutic products in comparison to random integration (*57*).

By providing efficient, multiplexed integration of transgenes in dividing and non-dividing cells, the PASTE platform builds upon fundamental developments in both integrase and CRISPR biology to expand the scope of genome editing and enable new applications across basic biology and therapeutics.

## Acknowledgments

We would like to thank B. Desimone, F. Chen, Y. Cha, A. Serj-Hansen, G. Feng, J. Wilde, M. Calos, T. Aida, J. Joung, and M. Mittens for helpful discussions; P. Reginato, D. Weston, and E. Boyden for MiSeq instrumentation; S. Jacobs and A. Ainbinder for digital-droplet PCR instrumentations; S. Bhatia and S. March Riera for hepatocyte assistance; G. Paradis and M. Griffin for flow cytometry assistance; and J. Crittenden for editing the manuscript.

## Author contributions

O.O.A and J.S.G conceived the study. O.O.A., and J.S.G. designed and participated in all experiments. E.I.I., M.Y., and C.S. led many of the experiments and assay readouts. R.N.K. helped with cell culture, clonings, plasmid sequencing, and next-generation sequencing. M.Y. and C.S. helped with digital-droplet PCR, sequencing experiments, and cloning. L.V. helped with various PASTE editing experiments. W.Z. synthesized mRNA and performed the electroporation experiments. J.L. performed the computational mining to uncover novel integrases and annotated these new systems. K.J. performed the ML modeling of the pooled atgRNA screening and developed a guide design software package. N.R., L.Z., and C.A.V. developed synthetic atgRNA guides. K.H., J.A.W, A.P.K, and A.E.Z. synthesized synthetic nicking guides. O.O.A., and J.S.G. wrote the manuscript with help from all authors.

## Data and materials availability

Sequencing data are available at Sequence Read Archive under BioProject accession number PRJNA700575. Expression plasmids are available from Addgene under UBMTA; support information and computational tools are available via the Abudayyeh-Gootenberg lab website (https://www.abugootlab.org/).

---

**Extended Data Figure 1:**
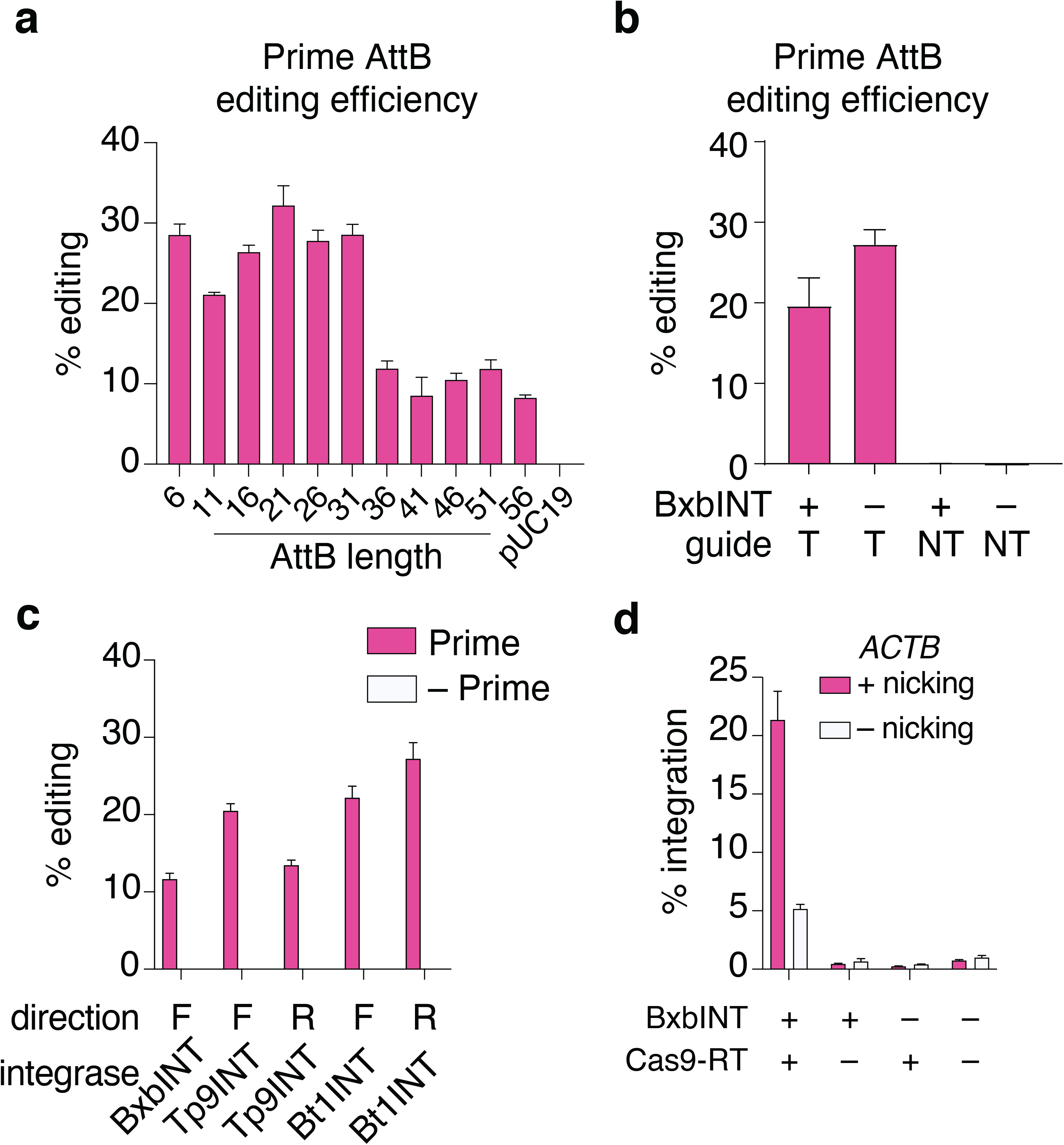
Evaluation of prime editing activity for diverse AttB sequences. a) Prime editing efficiency for the insertion of different length BxbINT AttB sites at *ACTB*. b) Prime editing efficiency for this insertion of a BxbINT AttB site at *ACTB* with targeting and non-targeting guides. c) Prime editing efficiency for the insertion of different integrases’ AttB sites at *ACTB*. Both orientations of landing sites are profiled (F, forward; R, reverse). d) PASTE editing efficiency for the insertion of EGFP at *ACTB* with and without a nicking guide. Data are mean (n= 3) ± s.e.m.

**Extended Data Figure 2:**
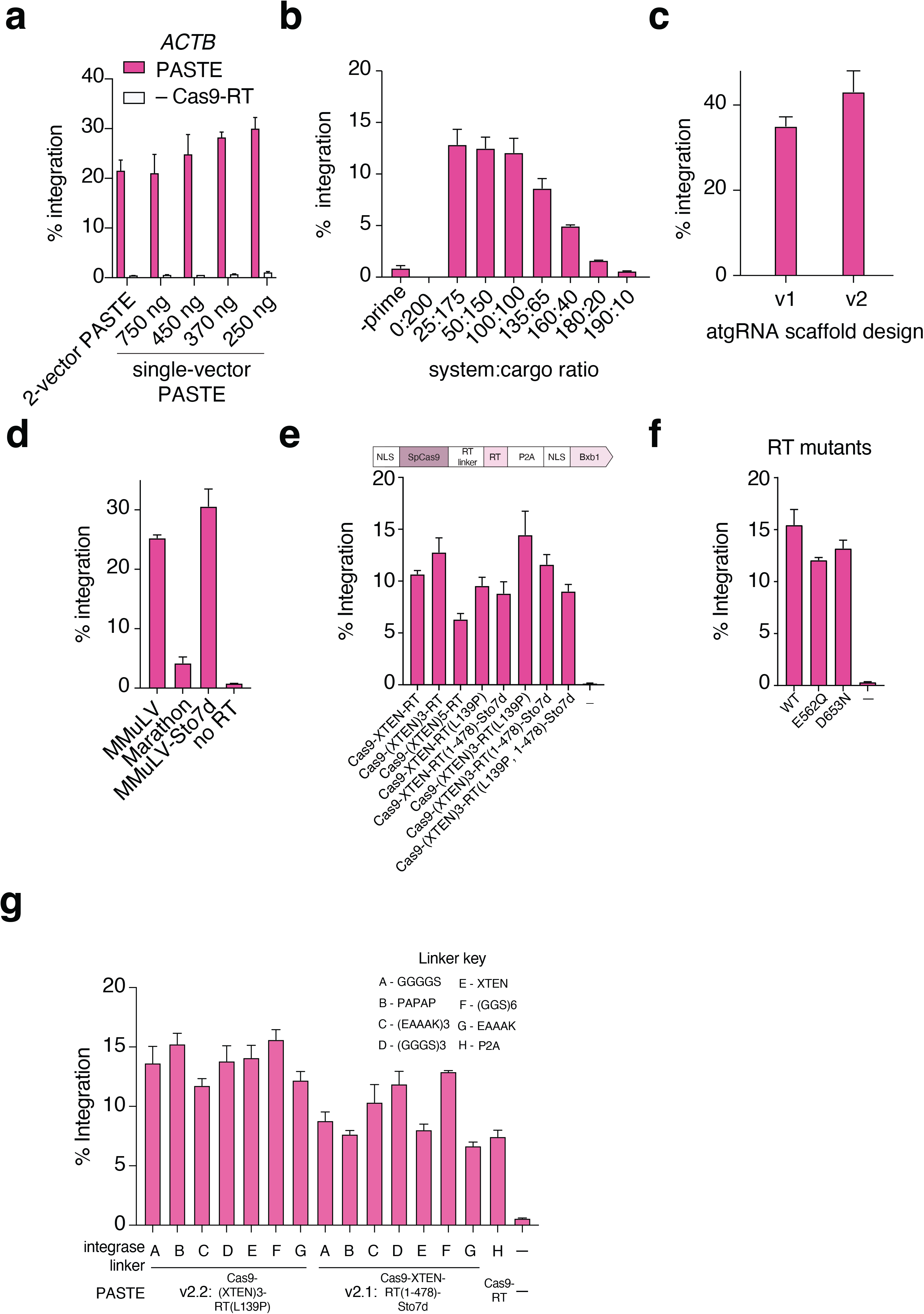
Optimization of PASTE editing by dosage titration and protein optimization. a) PASTE integration efficiency of EGFP at *ACTB* measured with different doses of a single-vector delivery of components. b) PASTE integration efficiency of EGFP at *ACTB* measured with different ratios of a single-vector delivery of components to the EGFP template vector. c) PASTE efficiency at the *ACTB* target compared between atgRNAs containing either the v1 or v2 scaffold designs. d) PASTE integration efficiency of EGFP at *ACTB* with different RT domain fusions. e) PASTE integration efficiency of EGFP at *ACTB* with different RT domain fusions and linkers. f) PASTE integration efficiency of EGFP at *ACTB* with mutant RT domains. g) Optimization of PASTE constructs with a panel of linkers and RT modifications for Gluc integration at the *ACTB* locus using atgRNAs with the v2 scaffold. Data are mean (n= 3) ± s.e.m.

**Extended Data Figure 3:**
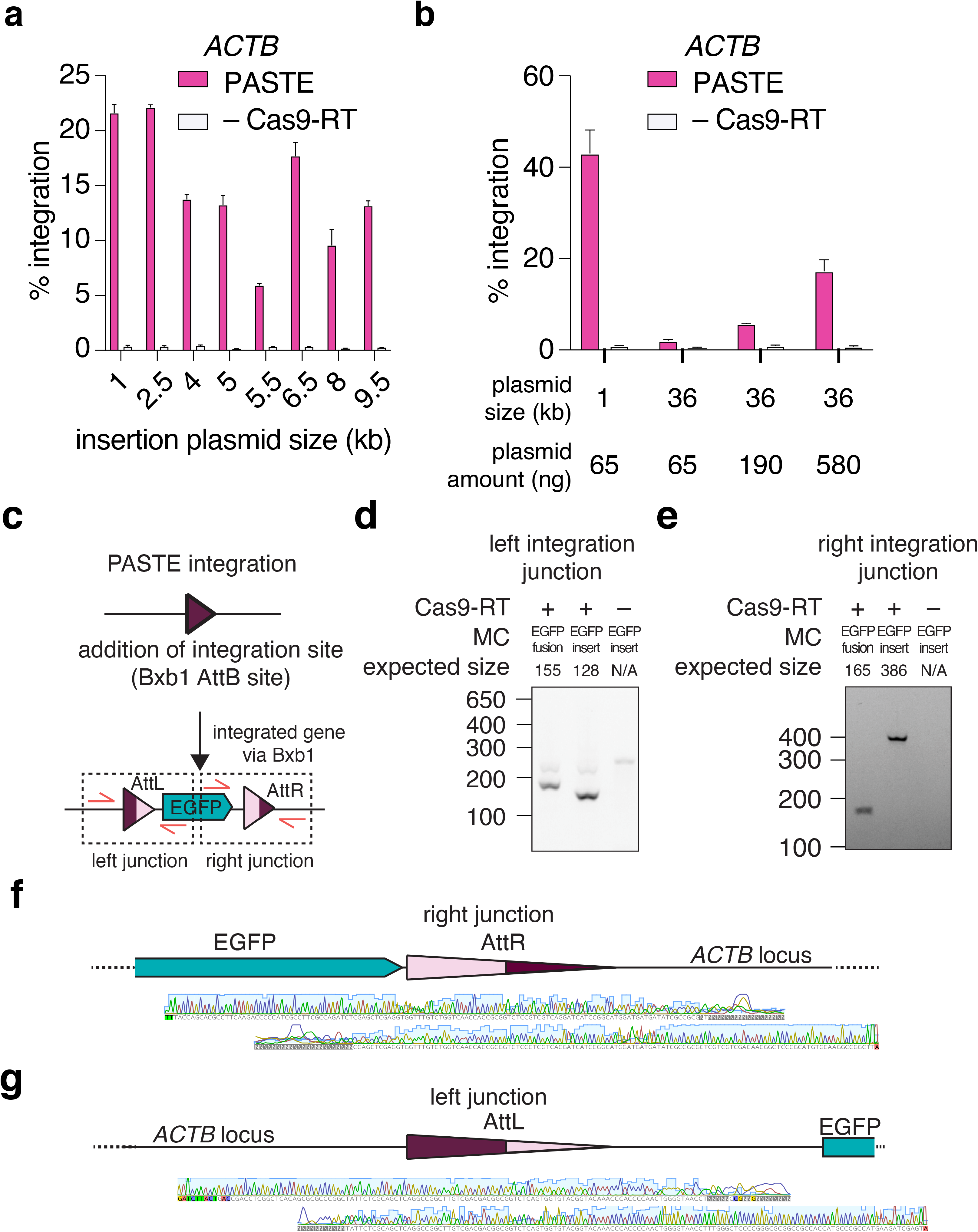
Characterization of PASTE payload sizes and integration junctions. a) PASTE integration efficiency at the ACTB locus of varying sized cargos transfected at a fixed DNA amount and variable molar ratio. b) PASTE integration efficiency at the ACTB locus of varying sized cargos transfected at a variable DNA amounts. c) Schematic of PASTE integration, including resulting AttR and AttL sites that are generated and PCR primers for assaying the integration junctions. d) PCR and gel electrophoresis readout of left integration junction from PASTE insertion of GFP at the ACTB locus. Insertion is analyzed for in-frame and out-of-frame GFP integration experiments as well as for a no prime control. Expected sizes of the PCR fragments are shown using the primers shown in the schematic in subpanel A. e) PCR and gel electrophoresis readout of right integration junction from PASTE insertion of GFP at the ACTB locus. Insertion is analyzed for in-frame and out-of-frame GFP integration experiments as well as for a no prime control. Expected sizes of the PCR fragments are shown using the primers shown in the schematic in subpanel A. f) Sanger sequencing shown for the right integration junction for an in-frame fusion of GFP via PASTE to the N-terminus of ACTB. g) Sanger sequencing shown for the left integration junction for an in-frame fusion of GFP via PASTE to the N-terminus of ACTB. Data are mean (n= 3) ± s.e.m.

**Extended Data Figure 4:**
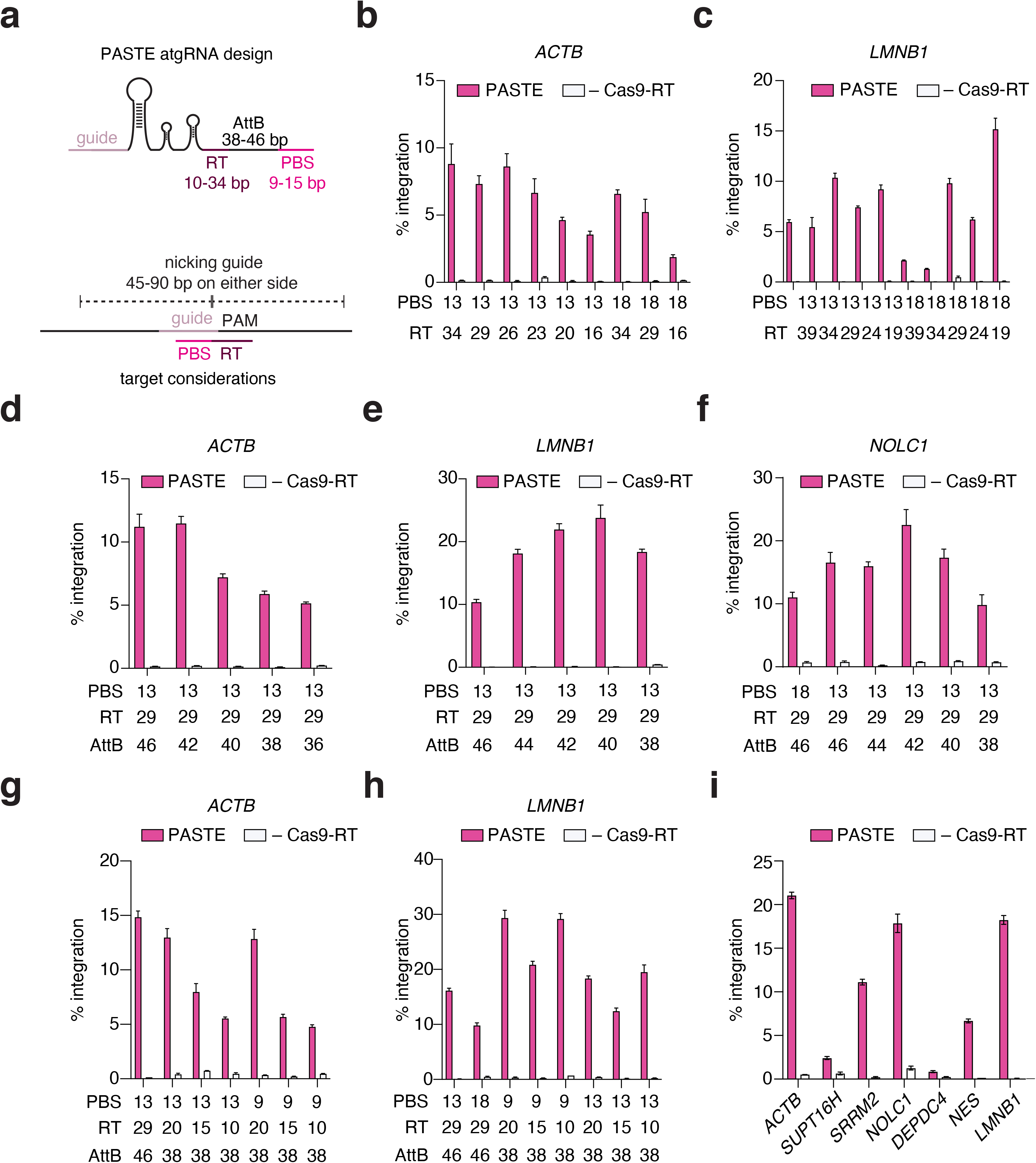
Validation of design rules for efficient PASTE insertion at endogenous genomic loci. a) Schematic of various parameters that affect PASTE integration of ∼1 kb GFP insert. On the atgRNA, the PBS, RT, and AttB lengths can alter the efficiency of AttB insertion. Nicking guide selection also affects overall gene integration efficiency. b) The impact of PBS and RT length on PASTE integration of *GFP* at the *ACTB* locus. c) The impact of PBS and RT length on PASTE integration of GFP at the *LMNB1* locus. d) The impact of AttB length on PASTE integration of *GFP* at the *ACTB* locus. e) The impact of AttB length on PASTE integration of GFP at the *LMNB1* locus. f) The impact of AttB length on PASTE integration of *GFP* at the *NOLC1* locus. g) The impact of minimal PBS, RT, and AttB lengths on PASTE integration efficiency of *GFP* at the *ACTB* locus. h) The impact of minimal PBS, RT, and AttB lengths on PASTE integration efficiency of *GFP* at the *LMNB1* locus. i) PASTE integration efficiency of EGFP at varying endogenous loci. Data are mean (n= 3) ± s.e.m.

**Extended Data Figure 5:**
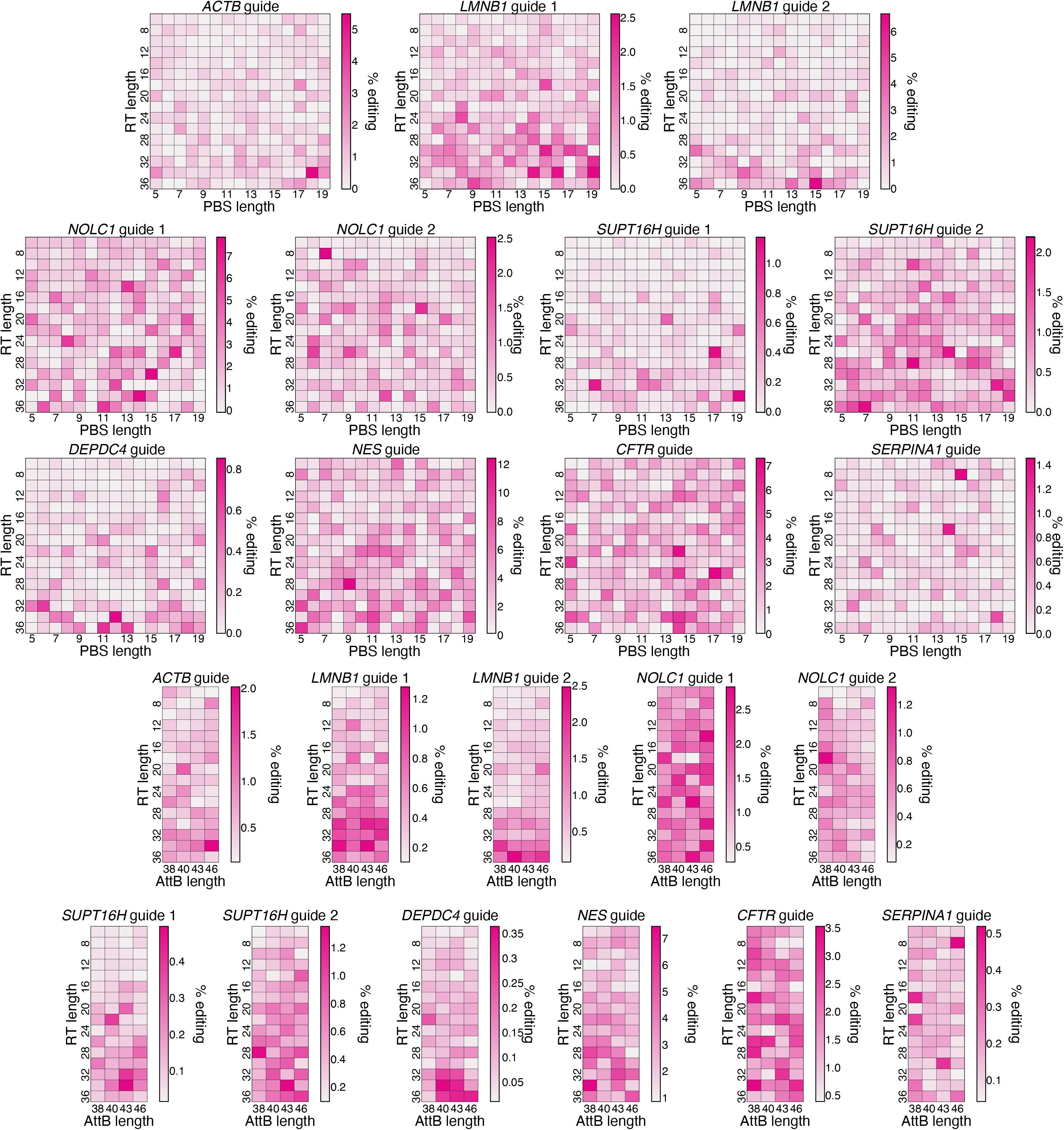
Heatmaps depicting the effect of PBS, RT, and AttB lengths on atgRNA efficiency of attachment site editing from high-throughput pooled screening of 10,580 guides targeting a variety of loci.

**Extended Data Figure 6:**
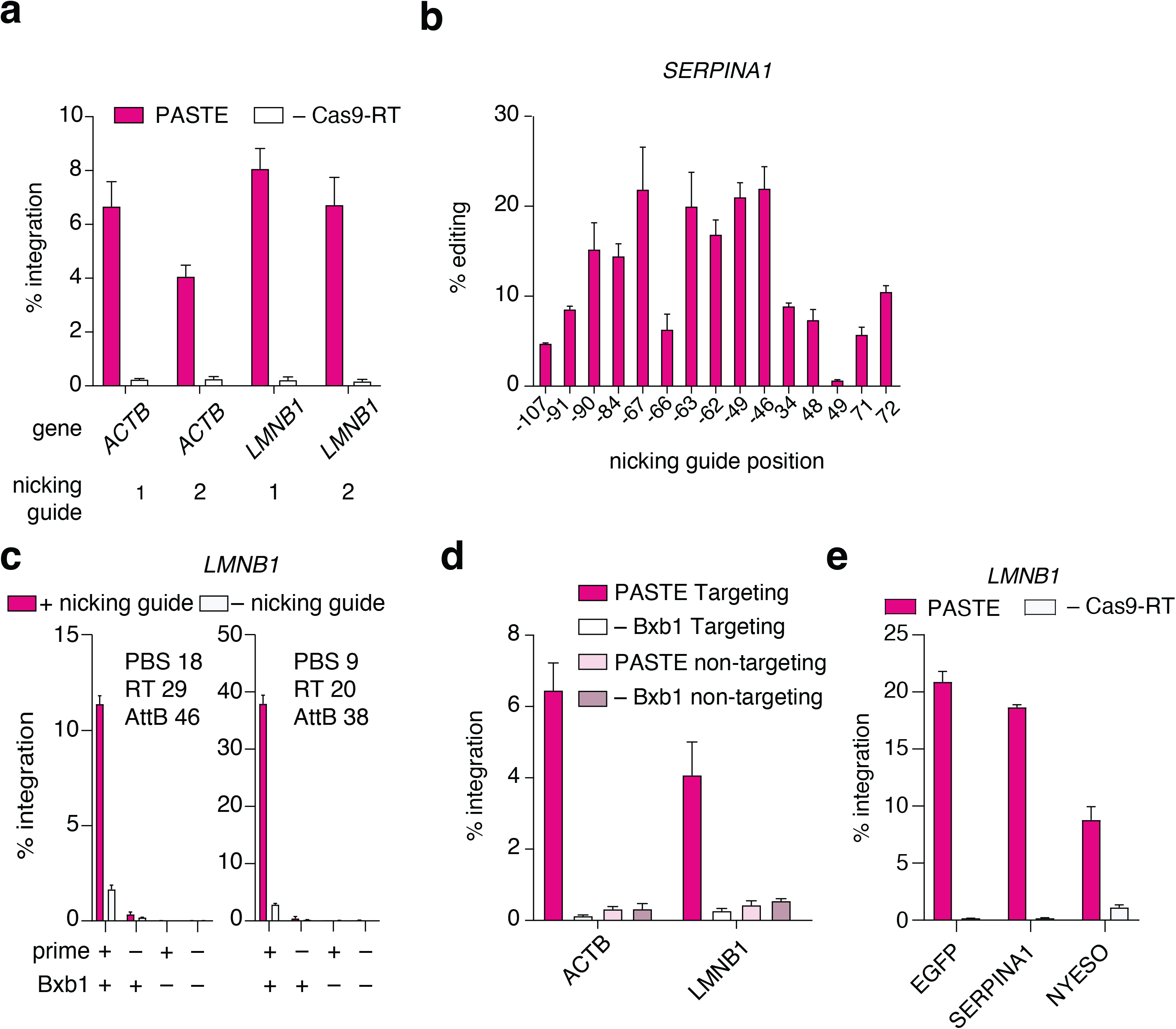
Effect of nicking guides on insertion of diverse cargos. a) PASTE insertion efficiency at *ACTB* and *LMNB1* loci with two different nicking guide designs. b) Attachment site editing at the SERPINA1 locus with a panel of different nicking guides at varying distances. c) Effect of nicking guides on PASTE integration efficiency at the LMNB1 locus with two different atgRNA designs. d) PASTE editing efficiency at *ACTB* and *LMNB1* with target and non-targeting spacers and matched atgRNAs with and without BxbINT expression. e) Integration of a panel of different gene cargo at LMNB1 locus via PASTE. Data are mean (n= 3) ± s.e.m.

**Extended Data Figure 7:**
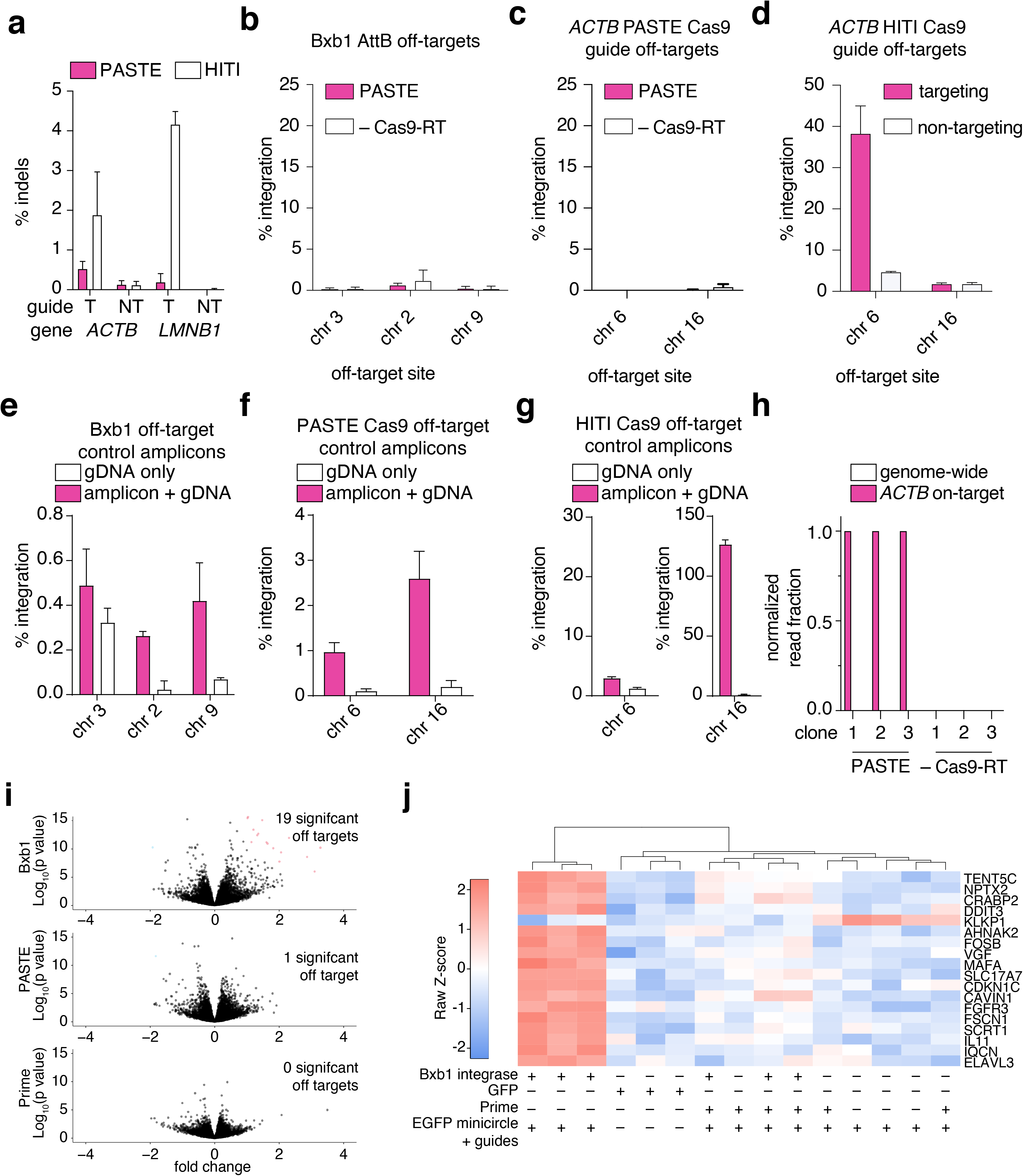
Further characterization of PASTE specificity and effects on cellular transcriptome. a) Comparison of indel rates generated by PASTE and HITI mediated insertion of EGFP at the ACTB and LMNB1 loci in HepG2 cells. b) GFP integration activity at predicted BxbINT off-target sites in the human genome. c) GFP integrations activity at predicted PASTE *ACTB* Cas9 guide off-target sites. d) GFP integration activity at predicted HITI *ACTB* Cas9 guide off-target sites. e) Validation of ddPCR assays for detecting editing at predicted BxbINT off-target sites using synthetic amplicons. f) Validation of ddPCR assays for detecting editing at predicted PASTE ACTB Cas9 guide off-target sites using synthetic amplicons. g) Validation of ddPCR assays for detecting editing at predicted HITI ACTB Cas9 guide off-target sites using synthetic amplicons. h) Analysis of on-target and off-target integration events across 3 single-cell clones for PASTE and 3 single-cell clones for no prime condition. i) Volcano plots depicting the fold expression change of sequenced mRNAs versus significance (p-value). Each dot represents a unique mRNA transcript and significant transcripts are shaded according to either upregulation (red) or downregulation (blue). Fold expression change is measured against ACTB-targeting guide-only expression (including cargo). j) Top significantly upregulated and downregulated genes for BxbINT-only conditions. Genes are shown with their corresponding Z-scores of counts per million (cpm) for BxbINT only expression, GFP-only expression, PASTE targeting ACTB for EGFP insertion, Prime targeting ACTB for EGFP expression without BxbINT, and guide/cargo only. Data are mean (n= 3) ± s.e.m.

**Extended Data Figure 8:**
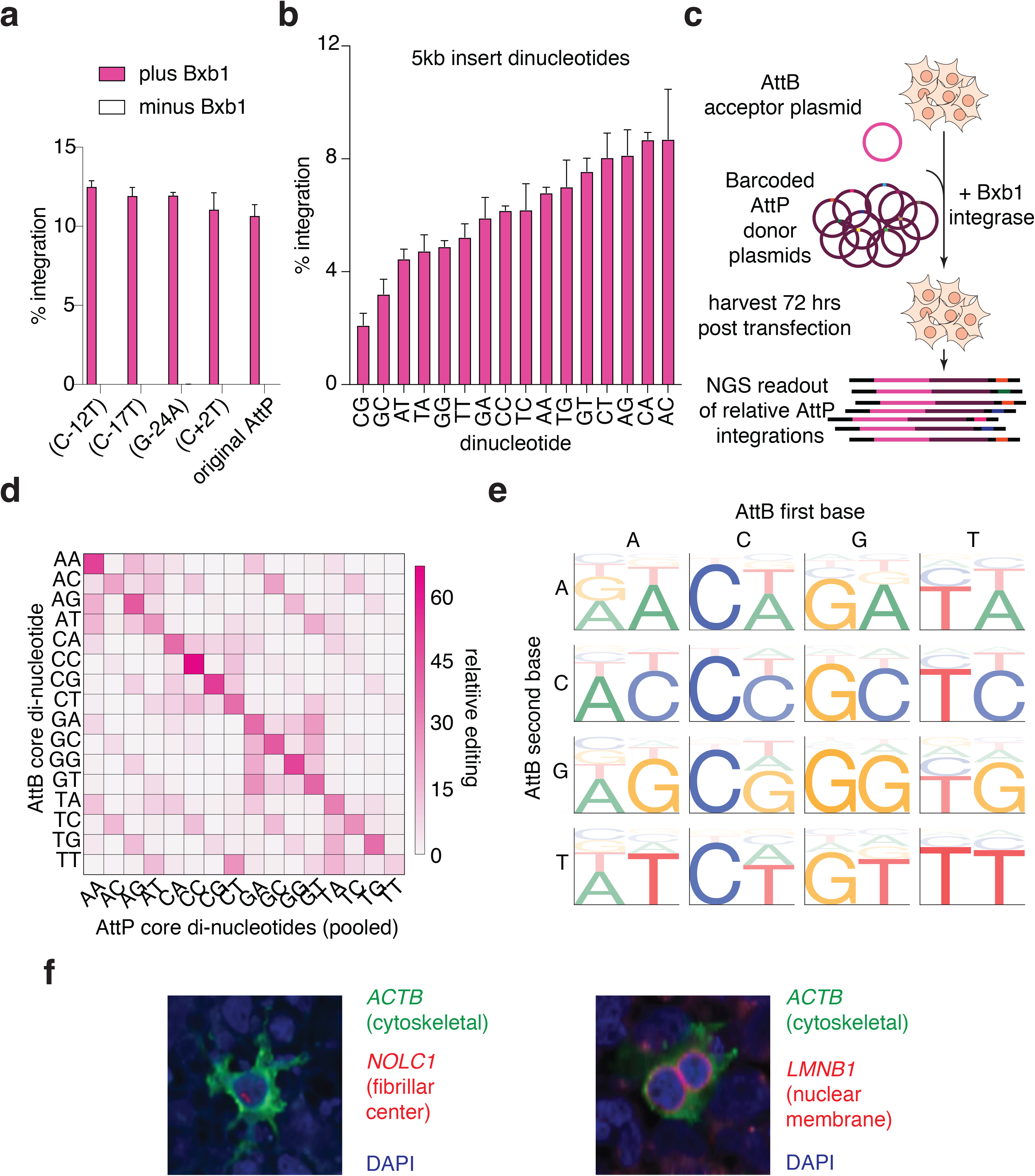
Additional characterization of AttP mutants for improved editing and multiplexing. a) AttP single mutants are characterized for PASTE EGFP integration at the ACTB locus. b) Characterization of integration of a 5 kb payload at the ACTB locus with all 16 possible dinucleotides for AttB/AttP pairs between the atgRNA and minicircle. c) Schematic of the pooled AttB/AttP dinucleotide orthogonality assay. Each AttB dinucleotide sequence is co-transfected with a barcoded pool of all 16 AttP dinucleotide sequences and BxbINT, and relative integration efficiencies are determined by next generation sequencing of barcodes. All 16 AttB dinucleotides are profiled in an arrayed format with AttP pools. d) Relative insertion preferences for all possible AttB/AttP dinucleotide pairs determined by the pooled orthogonality assay. e) Orthogonality of BxbINT dinucleotides as measured by a pooled reporter assay. Each web logo motif shows the relative integration of different AttP sequences in a pool at a denoted AttB sequence with the listed dinucleotide. f) Representative fluorescence images of multiplexed PASTE gene tagging of *ACTB, LMNB1,* and *NOLC1*. Data are mean (n= 3) ± s.e.m.

**Extended Data Figure 9:**
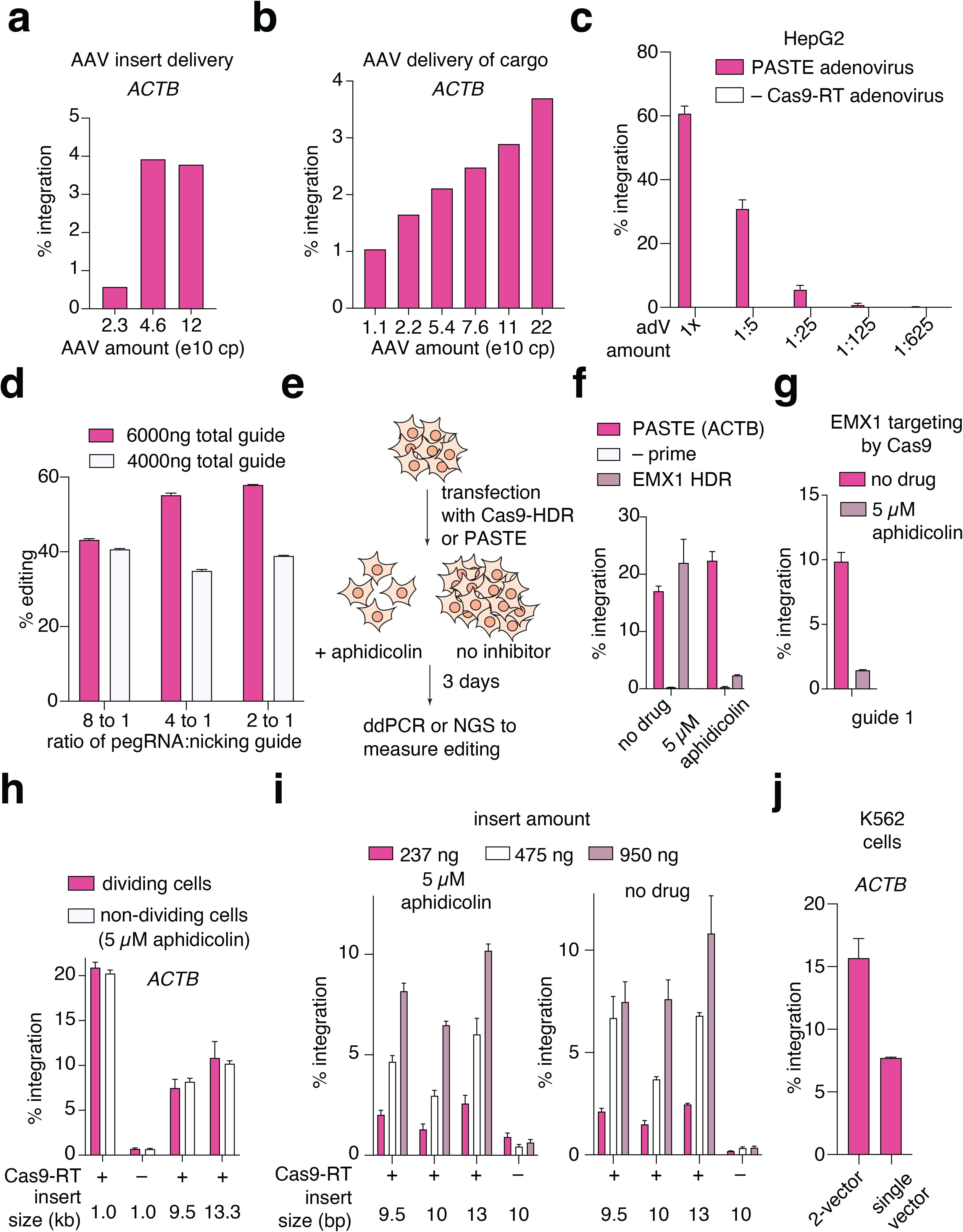
Evaluation of viral templates for PASTE and characterization of editing in non-dividing cells. a) PASTE integration of GFP at the ACTB locus with the GFP template delivered via AAV in HEK293FT cells. b) PASTE integration of GFP at the ACTB locus with the GFP template delivered via AAV at different doses in HEK293FT cells. c) PASTE integration of GFP at the ACTB locus with the GFP template delivered via AAV at different doses in HepG2 cells. d) Attachment site editing efficiency at the LMNB1 locus using PASTE delivered as mRNA with synthetic atgRNA and nicking guides. e) Schematic of PASTE performance in the presence of cell cycle inhibition. Cells are transfected with plasmids for insertion with PASTE or Cas9-induced HDR and treated with aphidicolin to arrest cell division. Efficiency of PASTE and HDR are read out with ddPCR or amplicon sequencing, respectively. f) Editing efficiency of single mutations by HDR at EMX1 locus with two Cas9 guides in the presence or absence of cell division read out with amplicon sequencing. g) HDR mediated editing of the EMX1 locus is significantly diminished in non-dividing HEK293FT cells blocked by 5 µM aphidicolin treatment. h) Integration efficiency of various sized GFP inserts up to 13.3 kb at the *ACTB* locus with PASTE in the presence or absence of cell division. i) Effect of insert minicircle DNA amount on PASTE-mediated insertion at the *ACTB* locus in dividing and non-dividing HEK293FT cells blocked by 5 µM aphidicolin treatment. j) PASTE efficiency of EGFP integration at the ACTB locus in K562 cells. Data are mean (n= 3) ± s.e.m.

**Extended Data Figure 10:**
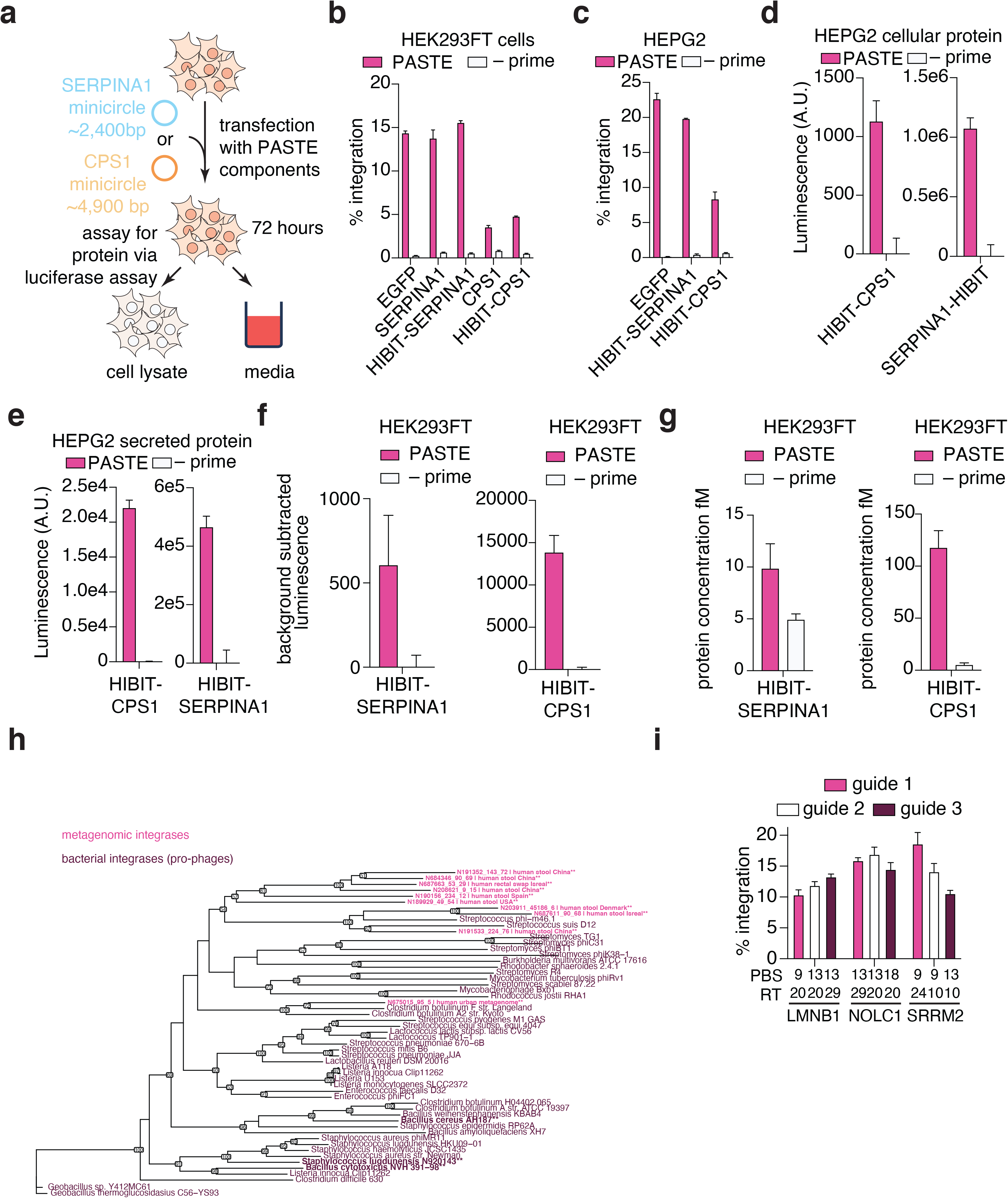
Therapeutic applications of PASTE and further characterization of novel integrases. a) Schematic of protein production assay for PASTE-integrated transgene. *SERPINA1* and *CPS1* transgenes are tagged with HIBIT luciferase for readout with both ddPCR and luminescence. b) Integration efficiency of SERPINA1 and CPS1 transgenes in HEK293FT cells at the *ACTB* locus. c) Integration efficiency of SERPINA1 and CPS1 transgenes in HepG2 cells at the *ACTB* locus. d) Intracellular levels of SERPINA1-HIBIT and CPS1-HIBIT in HepG2 cells. e) Secreted levels of SERPINA1-HIBIT and CPS1-HIBIT in HepG2 cells. f) Integration of SERPINA1 and CPS1 genes that are HIBIT tagged as measured by a protein expression luciferase assay. g) Integration of SERPINA1 and CPS1 genes that are HIBIT tagged as measured by a protein expression luciferase assay normalized to a standardized HIBIT ladder, enabling accurate quantification of protein levels. h) Phylogenetic tree of a smaller subset of bacterial integrases (pro-phages) and metagenomic integrases identified by our computational discovery pipeline. i) Panel of atgRNAs for integration of EGFP at three endogenous targets using PASTE with BceINT. Data are mean (n= 3) ± s.e.m.

## Materials and methods

### Cloning of atgRNAs, nicking guides

atgRNA and nicking guides were cloned by Golden Gate assembly of PCR products. Guide products were amplified by PCR (KAPA HiFi HotStart DNA Polymerase, Roche) off of the Cas9 sgRNA scaffold, with the forward primer containing spacer sequences and the reverse primer containing desired PBS, RT and AttB insertion sequences, in the case of the atgRNA. PCR products were purified by gel extraction (Monarch gel extraction kit, NEB), and assembled in a Golden Gate assembly containing 6.25 ng pU6-atgRNA-GG-acceptor (Addgene #132777), purified PCR product (approximately 2-to-4-fold molar excess), 0.125 µL Fermentas Eco31I (Thermo Fisher Scientific), 0.0625 µL T7 DNA ligase (Enzymatics),0.0625µL of 20 mg/mL Bovine Serum Albumin (NEB), 2x Reaction Ligation Buffer (Enzymatics) and water for 6.25µL total reaction volume. Reactions were incubated between 37°C and 20°C for 5 minutes each for a total of 15 cycles. 2μL of assembled reactions were transformed into 20 μL of competent Stbl3 generated by Mix and Go! competency kit (Zymo) and plated on agar plates supplemented with appropriate antibiotics. After overnight growth at 37°C, colonies were picked into Terrific Broth (TB) media (Thermo Fisher Scientific) and incubated with shaking at 37°C for 24 hours. Cultures were harvested using QIAprep Spin Miniprep Kit (Qiagen) according to the manufacturer’s instructions. All guides used in experiments are summarized in supplementary table 4 and all AttB sequences used in the paper are listed in supplementary table 5.

### Cloning of PASTE and cargo constructs

Expression constructs for Cas9-RT fusions, RT mutants, integrase and recombinases, and Bxb1 mutants were cloned into for mammalian expression via Gibson cloning using Hifi Assembly mix (NEB) according to manufacturer’s instructions. All enzyme expression plasmids used in mammalian experiments are summarized in supplementary table 6. Sequences of linkers used are listed in supplementary table 2, and sequences of Bxb1 and RT mutants are listed in supplementary table 3. For cloning of minicircle cargo plasmids, the Bxb1 or equivalent integrase/recombinase AttP sites and the cargo sequence were introduced into a minicircle parental plasmid with Gibson cloning using Hifi Assembly mix (NEB) according to manufacturer’s instructions. The parental plasmid was digested in order the sequences to be cloned between the bacterial attB and attP sites recognized by the ZYCY10P3S2T E. *coli* Minicircle Strain (Systems Bioscience). All transgene and cargo plasmids used in experiments are summarized in supplementary table 7.

For all Gibson clonings, 2 μL of assembled reactions were transformed into 20 μL of competent Stbl3 generated by Mix and Go! competency kit (Zymo) and plated on agar plates supplemented with appropriate antibiotics. After growth overnight at 37°C, colonies were picked into Terrific Broth (TB) media (Thermo Fisher Scientific) and incubated with shaking at 37°C for 24 hours. Cultures were harvested using QIAprep Spin Miniprep Kit (Qiagen) according to the manufacturer’s instructions.

For screening novel integrases discovered computationally, gene fragments were synthesized by Twist Biosciences. These genes were then cloned into separate expression vectors for comparing activity on reporters in mammalian cells and top integrases were cloned into PASTE vectors fused to SpCas9-RT constructs.

### Minicircle production

For the production of the minicircle plasmids containing only the integrase AttP site and the transgene sequence, the parental plasmid was transformed into the ZYCY10P3S2T E. coli Minicircle Strain (System Biosciences, catalog #: MN900A-1) overnight at 37°C. The next day a colony was picked into TB media containing Kanamycin antibiotic and grown for approximately 12 hours of incubation at 32°C in an incubator shaker. When the OD_600_ reached 4-6, the induction media was added in 1:1 ratio to the sample. For the preparation of 100.5 mL induction media, 100 mL of Lysogeny Broth media (Thermo Fisher Scientific) were mixed with 400 µL of 10 M Sodium Hydroxide Solution (Sigma Aldrich) and 100 µL of 20% L-Arabinose (Sigma Aldrich). The induced bacterial culture was then incubated at 32°C in the shaker for 4-5 hours. After spinning down at 5000 x g for 15 minutes, the media was removed leaving only the cell pellet at the bottom of the tube. For the purification of the DNA plasmid, an endotoxin-free plasmid midiprep DNA purification took place using NucleoBond Xtra Midi EF kit (Takara Bio) following the manufacturer’s protocol. Minicircle digestion was then confirmed using restriction enzymes and subsequent gel electrophoresis that allowed for interpretation of the minicircle and parent plasmid fractions in the purified DNA.

### Mammalian cell culture

HEK293FT cells (American Type Culture Collection (ATCC) - CRL32156) were cultured in Dulbecco’s Modified Eagle Medium with high glucose, sodium pyruvate, and GlutaMAX (Thermo Fisher Scientific), additionally supplemented with 10% (v/v) fetal bovine serum (FBS) and 1× penicillin-streptomycin (Thermo Fisher Scientific). For puromycin selection, HEK293FT cells were replated at a 1:3 dilution one day post-transfection into media supplemented with 1 µg/mL final concentration puromycin (Thermo Fisher Scientific). HEPG2 cells (American Type Culture Collection (ATCC – HB8065) were seeded in Eagle’s Minimum Essential Medium (Thermo Fisher Scientific), additionally supplemented with 10% (v/v) FBS, at 37°C and 5% CO2. Adherent cells were maintained at confluency below 80-90% at 37°C and 5% CO2. K562 cells (American Type Culture Collection (ATCC) - CCL-243) were cultured in Gibco Roswell Park Memorial Institute 1640 Medium (Thermo Fisher Scientific), additionally supplemented with 10% (v/v) FBS, and they were maintained at 37°C and 5% CO2. Primary human peripheral blood CD8+ T cells (Stemcell Technologies - #70027) were expanded using fresh complete ImmunoCult™-XF T Cell Expansion Medium (Stemcell Technologies - #10981) additionally supplemented with cytokines (Human Recombinant IL-2; Stemcell Technologies - #78036). To stimulate the cells, 25 µL/mL of ImmunoCult™ Human CD3/CD28/CD2 T Cell Activator (Stemcell Technologies - #10970) were used. Primary human hepatocytes pooled from 5 donors (Thermo fisher scientific #HMCPP5) were plated on collagen coated 96 well plates and transfected 24 hours post-plating. 96 well plates were coated using Collagen I, Rat Tail (Thermo fisher scientific #A10483-01). Stock Collagen I was diluted to 50 µg/mL with 20 mM acetic acid (# A6283) and added to plates at 5 μg/cm². Plates were incubated at room temperature for 1 hour then rinsed three times with sterile 1X PBS. Thawed hepatocytes were transferred into Hepatocyte Thaw Medium (Thermo fisher Scientific #CM7500) and centrifuged at 100 x g for 10 minutes at room temperature. Pelleted cells were resuspended and plated at 2.5e⁴ using William’s E Medium (Thermo fisher scientific #A1217601) supplemented with Primary Hepatocyte Thawing and Plating Supplements (Thermo fisher scientific #CM3000). Initial media change occurred 6 hours post plating, with subsequent media changes occurring every 24 hours using William’s E Medium supplemented with Primary Hepatocyte Maintenance Supplements (Thermo fisher scientific #CM4000).

### Transfection

Cells were plated at 5-15K the day prior to transfection in a 96-well plate coated with poly-D-lysine (BD Biocoat). HEK293FT and HepG2 cells were transfected with Lipofectamine 2000 (Thermo Fisher Scientific) and GenJet HepG2 reagent (SignaGen, SL100489-HEPG2), respectively, according to manufacturer’s specifications. For PASTE insertions, 100 ng atgRNA guide-encoding plasmid, 250 ng cargo plasmid, 50 ng nicking guide-encoding plasmid, and 375 ng SpCas9-RT-P2A-Bxb1 complex-encoding plasmid were delivered to each well unless otherwise specified. For HITI insertions, 100 ng guide-encoding plasmid, 250 ng cargo plasmid, and 75 ng SpCas9 plasmid were delivered to cells. For HDR insertion of a large EGFP cargo, 100 ng guide RNA guide, 200 ng SpCas9 plasmid, and 250 ng insertion template plasmid were delivered to cells. Cells were replated 72 hours later via limiting dilution in order to isolate clonal outgrowth in a 96-well plate for quantification of fluorescent colonies compared to PASTE. For HDR gene editing at the EMX1 locus for non-dividing cell experiments, 300 ng of a single vector encoding the guide RNA, SpCas9, and HDR editing template were transfected and cells were harvested 72 hours later for analysis by next-generation sequencing. For PASTE experiments with hepatocytes, plasmids were transfected with standard lipofectamine 3000 protocols with 400 ng of total plasmid.

### Plasmid Electroporation

K562 and primary T cells were electroporated using a Lonza 4D-Nucleofector device (Lonza). The SF Cell Line 4D X Kit S (Lonza) was used for K562s and the P3 Primary Cell 4D Kit (Lonza) was used for the unstimulated primary T cells. Approximately 1.5e6 K562 cells were electroporated in a final volume of 20 µL in a 16-well nucleocuvette strip (Lonza). For the T-cell experiments, 7.25e6 primary T cells were electroporated in a final volume of 100 µL in a cuvette.

For the single vector and two vector PASTE systems delivered to K562 cells, 900 ng of prime-Bxb1 complex-encoding plasmid or 800 ng of prime-encoding plasmid and 100 ng of Bxb1-encoding plasmid were electroporated, respectively. For both systems, 250 ng cargo plasmid, 200 ng atgRNA guide-encoding plasmid and 80 ng RNA nicking guide-encoding plasmid were added.

For T-cell electroporations, 990 ng of a guide vector expressing both the atgRNA and nicking guide, 875 ng of the EGFP-containing minicircle plasmid, and 3,150 ng of the PASTE plasmid (SpCas9-RT-P2A-Bxb1) were electroporated.

Electroporations were performed according to the manufacturer’s protocol and after 72 hours the cells were harvested for genomic DNA isolation and digital droplet PCR quantification.

### Cloning of atgRNA efficiency screen library

atgRNA library members were computationally designed to cover corresponding ranges of PBS, RT, and AttB length. Each library member was also paired with a unique barcode, as well as additional padding sequence after the poly-T transcriptional terminator to maintain consistent oligo length. For each of the 12 spacer sequences in the library, corresponding library members were flanked by spacer-specific subpooling binding regions. The 10,580 member library was synthesized as a pool by Twist Biosciences and PCR amplified to generate 12 subpools. Each subpool was Golden Gate cloned into a corresponding backbone containing both the spacer sequence and a 200-300 nucleotide region for targeting. Each library was independently electroporated into Endura electrocompetent cells (Lucigen), plated on agar bioassay plates, and harvested next day for protein purification.

### Pooled Screening of atgRNA efficiency

The complete library was co-transfected with psPAX2 and pMD2.G with Lipofectamine 3000 (Thermo Fisher Scientific) to produce lentivirus for atgRNA library testing. Two days post-transfection, supernatant containing virus was harvested, filtered using 0.45 µm syringe filters, and titer via spinfection, puromycin selection, and Cell Titer Glow viability readout (Promega). After titer, the atgRNA viral library was used to infect 80M HEK239FT cells at a 0.3 MOI to ensure single integration. Post-spinfection, cells were selected for 2 days with puromycin, then allowed to expand and recover without drug for an additional two days, and then were transfected with either PASTE constructs or Bxb1 integrase controls. Three days after transfection, cells were harvested with the Quick gDNA midi kit (Zymo), and the corresponding library region was prepared for sequencing via PCR amplification. Prepared libraries were pair-end sequenced on an Illumina NextSeq 500.

### Computational analysis of the pooled atgRNA screen

Forward reads were trimmed to the corresponding barcode region to extract barcode sequences. Extracted barcodes were paired with corresponding targeted regions in the reverse read, which were trimmed to the region within 20 nucleotides of the putative AttB region. To test for the presence or absence of editing, the region corresponding to the editing target was aligned to either the AttB-insertion outcome or the wildtype outcome, with reads aligning closer to the AttB-insertion outcome being ranked as edited. Editing frequency was then taken as the ratio of edited to total reads for each barcode, with a psuedocount adjustment of 1.

## Multilayer perceptron modelling of atgRNA efficiency

Three different sequence-to-function models are considered for accurate prediction of atgRNA efficiency including simple linear/logistic regression, random forest classifier, and multilayer perceptron (MLP) classifier. After the initial round of screening, we found that MLP classifier performed the best and we decided to move forward with a two hidden layer MLP model built in pytorch. After initial optimization, the MLP classifier contains an input layer of 125 neurons, a first hidden layer of 512 neurons, a second hidden layer of 10 neurons, and an output layer of 2 neurons. RELU is used as the activation function and a dropout rate of 0.1 is applied for each layer. The output layer is transformed by a softmax function to predict probability for each class. To represent atgRNA as a vector, we considered simple one hot vector or k-mer breakdown of atgRNA. We varied the k-mer breakdown from 1 to 7 and found that breaking atgRNA into short 3-nucleotide sequence (3-mer) is the most effective in training MLP. Padding is applied to atgRNA sequences that are shorter than 198 nt with “N” as the padding element to fulfill the input matrix to a uniform size. During the training of the MLP model, we varied the Adam optimizer’s learning rate from 0.0001 to 0.01, batch size from 30 to 100, and epoch number from 10 to 100. We minimized the validation loss in a 5-fold cross validation algorithm with the cross entropy loss as the loss function and chose a learning rate of 0.001, epoch of 50, and batch size of 64 as the final training hyperparameters. ROC_AUC curve is carried out using the sklearn’s roc_auc function. Codes to predict atgRNA efficiency and corresponding setup instructions are available at the following github repositories. (https://github.com/abugoot-lab/atgRNA_rank)

### mRNA and synthetic guides electroporation

Before *in vitro* transcription, the DNA template was linearized by FastDigest MssI restriction enzyme (Thermo Fisher) and purified by QIAprep 2.0 Spin Miniprep Columns (Qiagen). PASTE mRNA (SpCas9-RT-P2A-Bxb1) and Bxb1 mRNA (NLS-Bxb1) were transcribed and poly-A tailed using the HiScribe™ T7 ARCA mRNA Kit (NEB, E2065S) with 50% supplement of 5-Methyl-CTP and Pseudo-UTP (Jena Biosciences), following the manufacturer’s protocol. The mRNA was then cleaned up using the MEGAclear™ Transcription Clean-Up Kit (Thermo Fisher, AM1908). For circularized Bxb1 mRNA, *in vitro* transcription was conducted using the HiScribe™ T7 ARCA mRNA Kit without modified nucleotides or the poly-A tailing step. mRNA was subsequently circularized as previously reported(*48*) and cleaned up again using the MEGAclear™ Transcription Clean-Up Kit. Chemically modified synthetic atgRNA (Integrated DNA Technologies and Synthego Corporation) and nicking guide RNA (Synthego Corporation) were provided by the corresponding parties. HEK293FT cells were electroporated using a Lonza 4D-Nucleofector device and the SF Cell Line 4D-NucleofectorTM X Kit S (Lonza). For each sample, 4000 ng PASTE mRNA and 1000 ng Bxb1 mRNA were mixed with the designated amount of guide RNAs in a total volume of 15 μL SF buffer solution. Cells (2.0e5 per sample) were spun down at 100×*g* for 10 minutes, resuspended in 5 μL SF buffer solution, and added to the 15 μL RNA solution. The 20 μL mixture was placed in one well of the cuvette strip and subject to electroporation using the CM-130 program. Electroporated cells were resuspended in its culture media and incubated at 37°C and 5% CO_2_ for 72 hours before analysis.

### Genomic DNA extraction and purification

DNA was harvested from transfected cells by removal of media, resuspension in 50 µL of QuickExtract (Lucigen), and incubation at 65 °C for 15 min, 68 °C for 15 min, and 98 °C for 10 min. After thermocycling, lysates were purified via addition of 45 µL of AMPure magnetic beads (Beckman Coulter), mixing, and two 75% ethanol wash steps. After purification, genomic DNA was eluted in 25 μL water.

### Genome editing quantification by digital droplet polymerase chain reaction (ddPCR)

To quantify PASTE and HITI editing efficiency by digital droplet PCR, 24 µL solutions were prepared in a 96-well plate containing 1) 12 µL 2x ddPCR Supermix for Probes (Bio-Rad) 2) primers for amplification of the integration junction at 250 nM-900 nM, 3) FAM probe for detection of the integration junction amplicon at 250 nM 4) 1.44 µL RPP30 HEX reference mix (Bio-Rad) 5) 0.12 µL FastDigest restriction enzyme for degradation of primer off-targets (Thermo Fisher) and 6) Sample DNA at 1-10 ng/µL. All primers and probes used for ddPCR are listed in supplementary table 8. 20 µL of reaction mix was transferred to a Dg8 Cartridge (Bio-Rad) and loaded into a QX2000 droplet generator (Bio-Rad). 40 µL droplets suspended in ddPCR droplet reader oil were transferred to a new 96-well plate and thermocycled according to manufacturer’s specifications. Lastly, the 96-well plate was transferred to a QX200 droplet reader (Bio-Rad) and the generated data were analyzed using Quantasoft Analysis Pro to quantify DNA editing.

### Genome editing quantification by targeted deep sequencing

To quantify integration of AttB/AttP pairs in the Bxb1 orthogonality assay and genome editing for prime editing and HDR integration at the *EMX1* locus, target regions were PCR amplified and analyzed by deep sequencing. Genomic DNA samples were isolated using 50 µL of QuickExtract (Lucigen) per well, and target regions were PCR amplified with NEBNext High-Fidelity 2X PCR Master Mix (NEB) based on the manufacturer’s protocol. PCR amplicon primers are listed in supplementary table 9. Barcodes and adapters for Illumina sequencing were added in a subsequent PCR amplification. Amplicons were pooled and prepared for sequencing on a MiSeq (Illumina). Reads were demultiplexed and analyzed with appropriate pipelines. To analyze the Bxb1 orthogonality assay, AttP barcodes were extracted and normalized to overall barcode counts, and WebLogos were generated with LogoMaker (*55*). To analyze prime and HDR editing, amplicons were analyzed using custom scripts to analyze the relative number of reads with the inserted sequence.

### Genome-wide off-target integration and novel integrase integration quantification by UMI TN5 and next generation sequencing

To quantify the off-target integration of cargo payloads by PASTE and HITI throughout the human genome, single cell clones were harvested three days post transfection with QuickExtract (Lucigen) and purified using AMPure magnetic beads (Beckman Coulter) according to the manufacturer’s protocol. Cellular gDNA was eluted in water and normalized to 6.25 ng/µL. A 2x Tn5 dialysis buffer was prepared with the following components according to Picelli et al 2014: 100 HEPES-KOH at pH 7.2, 0.2 M NaCl, 0.2 mM EDTA, 2 mM DTT, 0.2% Triton X-100, 20% glycerol). TN5 was assembled with equimolar pre annealed mosaic-end double stranded oligonucleotides by incubating the following components at RT for one hour: 25 µL 100uM (final concentration, see supplementary figure 10) oligonucleotide mix in TE, 80 µL 100% glycerol, 24 µL 2x TN5 dialysis buffer, 72 µL Tn5 at A280=3.0. 2.9 µL of normalized DNA was incubated with .1 µL of this TN5 oligonucleotide mix and .75 µL of 5x Tris-HCl buffer (50 mM Tris-HCl pH8, 25 mM MgCl2) for 10 minutes at 55°C. 1.875 µL of this TN5 transposition reaction were used as the template in a PCR reaction using SuperFi PCR Master mix platinum (Thermofisher) according to the manufacturer’s protocol. Next, 1 µL for this reaction was used as the template in a next generation sequencing reaction (see protocol below); UMI TN5 primers for genome-wide off target integration detection are listed in supplementary table 10. After NGS barcoding, all samples were diluted 1:1 and pooled; 20 µL of this pool was run on a 1% agarose gel and a smear from 280-800 bp was extracted, purified, and prepared for next generation sequencing on a MiSeq (Illumina).

To compare and quantify the integration efficiency of novel integrases, HEK293T cells were transfected with an atgRNA-expressing plasmid containing the attb site of the punitive integrase along with a minicircle and integrase-expressing plasmid; integration efficiency of the punitive integrase was measured as the integration of the minicircle into the atgRNA vector. To quantify this integration, the above UMI protocol was followed with different primer sets. The mosaic-end double stranded oligonucleotides used in TN5 preparation remained constant, as did the indexing reverse primer used in the SuperFi PCR mix and first round NGS thermocycling steps. The forward primers for these thermocycling steps were changed for ones with homology for the atgRNA acceptor plasmid. These UMI TN5 novel integrase reporter primers can also be found in supplementary table 10.

### Computational identification of Bxb1 and Cas9 Off-targets

To identify potential off-target sites for Bxb1 integration or Cas9 cleavage, similar sequences were identified in the human genome using either BLAST (*56*), for similar AttP sequences, or Cas9 off-target prediction algorithms (*57*). To validate successful amplicon generation by primer sets, positive control off-target amplicons were ordered as oligonucleotides, annealed, and tested by ddPCR. Off target sites are listed in supplementary table 11.

### Imaging

For sample preparation for imaging, cover slips were placed at the bottom of a 24-well plate prior to plating HEK293FT cells. After transfection at ∼70% confluency and incubation period of three days, the media was removed, and the wells were washed with 1 mL PBS pH 7.4 (Thermo Fisher Scientific). The cells were fixed with 500 µL of 4% Pierce Formaldehyde (Thermo Fisher Scientific) for 30 minutes. Another washing with 1 mL PBS pH 7.4 was performed three times. If no immunostaining was to be performed, slides were processed to be mounted.

If immunostaining was to be performed, the cells were blocked in 1 mL 2.5% goat serum (Cell Signaling Technology) and in 0.1% Triton-X (Sigma Aldrich) for 1 hour at room temperature. For the primary stain, the primary antibodies were mixed with 1.25% goat serum and 300 µL were added per well according to the following dilutions: 1:1500 for the anti-ACTB antibody (NB600-501SS, NovusBio), 1:200 for the anti-SRRM2 antibody (NBP2-55697, NovusBio), 1:200 for the anti-NOLC1 antibody (11815-1-AP, ProteinTech), and 1:200 for the anti-LMNB1 antibody (12987-1-AP, ProteinTech). After shaking overnight at 4 °C, the wells were washed three times with 1mL PBS pH 7.4. For the secondary staining, 1:1000 dilution of secondary antibody, either goat anti-mouse IgG Alexa Fluor 568 (Thermo Fisher Scientific, A-11004) or goat anti-rabbit IgG Alexa Fluor 647 (Thermo Fisher Scientific, A21244), were mixed with 1.25% goat serum. After 1 hour at room temperature, another washing step with PBS took place three times and slides were then mounted.

To mount the slides, a drop of ProLong Gold Antifade Mountant with DAPI (Thermo Fisher Scientific) was placed on the top of the slide and the cover slips were removed from the 24-well plate and inverted onto the drop. The coverslips were left to dry for 24 hours protected from the light at room temperature and then sealed with nail polish. For acquisition of images, a scanning laser confocal microscope (Zeiss LSM900) was used with a 40x oil objective and three different filter sets for visualizing EGFP, DAPI, and the immunofluorescence stain (either 568 or 647).

### AAV production and transduction

To produce AAV vectors for delivery of PASTE cargo, HEK293FT cells were transfected in T25 flasks using Lipofectamine 3000 (Thermo Fisher Scientific) with 1.6 µg GFP AAV cargo plasmid, 1.96 µg AAV8 capsid vector, and 4.13 µg AAV helper pAdDeltaF6 plasmid (Addgene #112867) per T25 flask according to manufacturer’s protocol. Two days after transfection, the media containing the loaded viral vector was filtered using a 0.45 μm filter (Sigma Aldrich) and the final product was stored at −80°C. One day after the transfection of PASTE components (PE2, Bxb1, nicking guide, and atgRNA) into HEK293FT cells, AAVs containing the GFP cargo template were introduced directly into the cells according to the indicated volumes. Three days after the transduction, the cells were harvested and ddPCR readout took place.

### AdV production and transduction

Adenoviral vectors were cloned using the AdEasy-1 system obtained from Addgene. Briefly, SpCas9-RT-P2A-Blast, Bxb1 and guide RNAs, and an EGFP carge gene were cloned into separate adenoviral template backbones and recombined to add the full Adenoviral genome with the AdEasy-1 plasmid in BJ5183 E. coli cells. These recombined plasmids were sent to Vector BioLabs for commercial production. For the EGFP cargo vector, it was added at 6.7e6 PFU per well of a 96 well plate of HEK293FT cells and 1.3e6 PFU per well of a 96 well plate of HepG2 cells. For experiments with the three-vector adenoviral delivery of all PASTE components, we used 8e5 PFUs of each viral vector per well of a 96 well plate of HEK293FT cells. For three-vector adenovirus delivery on HepG2 cells, we used 40e6 PFUs of the EGFP cargo vector, 10e6 of the SpCas9-RT-P2A-Blast vector, and 20e6 of the Bxb1and guides vector per well of a 96 well plate.

### Quantification of protein expression

Three days after the transfection of HepG2 cells, the Nano-Glo HiBiT Lytic Detection System (Promega) was used for the quantification of the HiBiT-tagged proteins, SERPINA1 and CPS1, in cell lysates or media. For the preparation of the Nano-Glo HiBiT Lytic Reagent, the Nano-Glo HiBit Lytic Buffer (Promega) was mixed with Nano-Glo HiBiT Lytic Substrate (Promega) and the LgBiT Protein (Promega) according to manufacturer’s protocol. The volume of Nano-Glo HiBiT Lytic Reagent added, was equal to the culture medium present in each well, and the samples were placed on an orbital shaker at 600 rpm for 3 minutes. After incubation of 10 minutes at room temperature, the readout took place with 125 gain and 2 seconds integration time using a plate reader (Biotek Synergy Neo 2). The control background was subtracted from the final measurements.

### Computational discovery of novel integrases

Prokaryotic genomic and metagenomic sequences were retrieved from various public databases and datasets including NCBI, ENA, Ensembl, MetaSUB, MGnify and JGI. Protein coding genes were predicted with Prodigal v2.6.3. The protein sequences were scanned for large serine integrase domains with hmmsearch (HMMER v3.3.2) using Pfam models PF00239, PF07508 and PF13408 with model specific gathering cutoffs. Protein sequences not containing at least a resolvase and recombinase domain were discarded and the remaining sequences were marked as putative large serine integrases. The source contigs of these putative integrases were passed to VirSorter v1.0.6 for prophage boundary prediction. Contigs that were predicted to have a prophage region completely overlapping with their putative integrases were passed to a dynamic programming algorithm to detect kmer matches of 2-18 bp between the 1000 bp around the predicted prophage boundaries and the 100 bp on the other side of the putative integrase gene. Matching kmers sites were then expanded to 50 bp and scanned for inverted repeats. Sites with a high number of di-nucleotide inverted repeats (based on an experimentally derived cutoff) were nominated as putative attachment sites. To build a phylogenetic tree, the putative integrases were deduplicated and aligned with MUSCLE v5.0.1278 and trimmed with trimAl v1.4.1 using the -gappyout method. A tree was generated with FastTree v2.1.11 using the LG+CAT substitution model and visualized with Geneious Prime. Clustering to make clades was done with TreeCluster v1.0.3 using the max_clade method with a threshold of 8.5.

