## Supplementary Information for "Drag-and-drop genome insertion without DNA cleavage with CRISPR-directed integrases"

### Supplementary Table 1. List of integrases used for the development of PASTE.

| Database | SRA accession | bioproject_acc | nucleotide accession | protein accession or ORF ID | internal protein ID | Proposed names | Alternative names | organism/source | description | Sequence | Length | Group |
|---|---|---|---|---|---|---|---|---|---|---|---|---|
| ENA | SRS1205298 | PRJEB26277 | NA | NA | N189929_49_54 | SsuINT | NA | human gut metagenome | stool sample from male in USA | MEKNRAVLYLRLSKEDVDKVNKGDDSSSIKSQRLLLTDFALERGFKIVGVYSDDDESGLYDDRPDFERMMTDAKLDEFDIIIAKTQSRFSRNMEHIEKYLHHDLPNLGIRFIGAVDGVDTESDENKKSRQINGLVNEWYCEDLSKNIRSAFKAKMKDGQFLGSSCPYGYKKDPQNHNHLVVDDYAAKVVQKIFNLYLEGYGKAKIGSILSSEGILIPTLYKKDILKQNYHNSKALDTTQNWSYQTIHTILNNEVYLGHLIQNKVNTMSYKDKNKRILPKEKWIIVRNTHEPIITEEMFQDVQKLQKNRTRSVENIEPNGLFSGLIFCADCKHAMSRKYARRGEKGFVGYVCKTYKTQGKNFCESHSIDYDELEEAVLFSIKNEARSILQQEEIDELRKVQAYDETKSYYEMQLENIKSRMEKIEKYKKKTYDNYMDDLISRDDYKKYVTEYDKEIGGLKQQQELINSKTDLEKEISTQYDEWVEAFINYVDIDKLTREIVIELIEKIEVNKDGSINIYYKFKNPYIS | 527 | INTc |
| ENA | ERS396461 | PRJEB26280 | NA | NA | N190156_234_12 | SssINT | NA | human gut metagenome | stool sample from Spain | MNTVIYARYSAGPRQTDQSIDGQLRVCTEFCKQRGLTVVDTYCDRHISGRTDERPEFQRLIADAKAHKFEAVVVYKTDRFARNKYDSAIYKRELRRNGIQIFYAAEAIPEGPEGIILESLMEGLAEYYSAELAQKIKRGLNESALKCQSLGSGRPLGYTVDEQKHFQIDPESSQAVKTIFEMYIKGESNAAICDYLNARGLRTSQGNLFNKNSINRIIKNRKYIGEYRYNDIVVEGGMPAIISKETFCMAQAEMERRRTHRAPVSPKAEYLLAGKLFCGHCKGPMQGVSGTGKSGNKWYYYYCANTRGKERTCDKKQVSRDRLEKAVVDFTVRYILQENVLEELSKKVYAAQERQNNTASEIAFYEKKLAENKKAIANILRAIESGAMTQALPARLQELENEQTVIQGELSYLKGARLAFTEDQILFALLQHLDPRPGESERDYHRRIITDFVSEVYLYDDRMLIYFNISSADGKLKHADLSAIESGVFDAGLISSSSRASSFSTRCALI | 510 | INTd |
| ENA | ERS1015837 | PRJEB26832 | NA | NA | N191352_143_72 | SscINT | NA | human gut metagenome | stool sample from China | MNEKNLEIGAAYIRVSTDDQTELSPDAQLRVILEAAKKDGIIIPQEFVFMEDRGRSGRRADNRPEFQRMISTARQNPSPFRYLYLWKFSRFARNQEESAFYKGILRKKCGVTIKSVSEPIMEGMFGRLVEMIIEWSDEFYSVNLSGEVLRGMTQKALEHGYQLTPCLGYDAVGHGRPYVINEEQYQIVEFIHRSFFDGKDMTWIAREANRRGYHTRRGNPFDTRAVRIILTNSFYVGLVKWNDVTFQGTHECRESVTSVFSANQERLNRIHRPRGRRQASSCKHWLSGLLKCSICGASLGYNQTKDLTKRGHAFQCWKYTKGIHPGSCSVSSLKAEAAVLESLQMILETGEVEYTYEQREKHLDDNKLTLIQKSLERLDTKELRIREAYESGIDTLDEFKTNKARLQRERDQLMEELEELHSQEEPEDVPGKEILIERIQNVYDLLQSPDVDNDDKGNAVRSIIKKIVYIKESKTFCFYYYV | 482 | INTd |
| ENA | ERS1289677 | PRJEB26924 | NA | NA | N191533_224_76 | Ssc2INT | NA | human gut metagenome | stool sample from China | MERTIKVIQPGTVKIPTKKRVAAYARVSSGKDAMLHSLSAQVSYYSNMIQQKNEWSYVGIYADEAITGTKDRRVEFNRLIQDCTDGKIDMIITKSISRFARNTLTMLEVVRKLKNINVDVYFEKENIHSISGDGELMLTILASFAQEESRSVSENCKWRIRKGFEQGELINLRFLYGYRINKGKIEIYEKEAEIVRMIFDDYLNGEGCTRIGNKLRKMKVNKLRGGMWNSERVVDIIKNEKYTGNALLQKKYVKDH | 406 | INTc |

| | | | | | | | | | | | | |
|---|---|---|---|---|---|---|---|---|---|---|---|---|
| | | | | | | | | | | LSKKLVRNKGILTQYYAEGTHPAIIDIKTFEIAQKIMEANRTKFQGKCGSNRYLFTSKIECGICGKNYRHKDREGKSTWVCANHLKYGNSRCIAKPLNEEKLKKLINEALELKYFDEEIFIRNIKRIKVTGNQTIEFILKDGKVIEEGMI | | |
| ENA | ERS2655827 | PRJEB28245 | NA | NA | N203911_45186_6 | SsdINT | NA | human gut metagenome | stool sample from Denmark | MKKIKIDRAIQERPATRKQTRNEKIRQSLTEHVDVQVIPAITDREGYEKPKLRVCAYCRVSTDMDTQALSYELQVQNYTDYIRGNDEWRFAGIYADRGISGTSLKHRDEFNRMIEDCKAGKIDLIITKAVTRFARNVLDCISTIRMLKQLEHPVAVYFETERINTLDTTSETYLGLISLFAQGESESKSESLKWSYIRRWKRGTGIYPAWSLLGYEMGEDGKWQIVEAEAELVRIIYDMYLNGYSSPQIAEILTRSGVPTATNQTVWSSGGVLGILRNEKYCGNVLCQKTMTVDVFSHKAIKNTGQKTQYFIEGHHDPIILRSDWDRVQQMIDEKYYRKRRGRRTKPRIVLKGCLAGFTQIDLDWDEDDIARIFYSTTPAAEVATPAMADHIEIIKVKGEN | 401 | INTc |
| ENA | SRS294942 | PRJEB30046 | NA | NA | N208621_9_15 | SmcINT | NA | human gut metagenome | sample from 72-year-old male from China | MKTAAAYIRVSTDDQVEYSPDSQIKLIRDYAKRNDYILPDEFIFRDDGISGKSAKHRPEFTKMIALAKSPEHPFDAILVWKFSRFARNQEESIVFKNILRKIGVEVRSVSEPISEDPFGSLVERIIEWTDEYYIINLSGEVKRGMLEKISRGQPVVPPPVGYKMENGQYIPDENAHFIKEIFEAYAAGEGARHIAQRLAAQGCLTKRGNPIDNRFVDYVLHNPVYIGKLRWSVNSHAASSRHYDSADIIVFDGTHEPLISSELWESVQKRLHEVKTLYPKYQRREQPVSFMLKGLVRCSSCGSTLCYCRTSEPSLQCHSYARGSCRQSHSINIATANEAVIKGLQLAVDKLDFAIAPAKPHYSADAPGTNKLLAAEYKKMERIKAAYANGTDTLEEYAANKKKISAEIARLEAELQQESNVKPINKKAFAKRVSEIIKYISDPHNSEAAKNQALRTVISYIIFDRAATTFNIIFHF | 476 | INTd |
| MetaSUB | NA | NA | NA | NA | N675015_95_5 | UhmINT | NA | urban human microbiome | NA | MKIAIYARKSKYSPTGESVENQIQLCKEYLQAKYKSETLEIDEYKDEGYSGGNTNRPDFKKLIAQIEDYDMLICYRLDRISRNVADFSSTLTLLQNNKCDFVSIKEQFDTTSPMGRAMIYISSVFAQLERETIAERIRDNMMELAKMGRWLGGTIPMGFDSEPITFIDENMKERSMTKLIPNVEELKVIELIYEKYLQLGSMGKVVTYLLQNNIKTKKGKDFTLGSIKVILTNPIYVKANQEVVNHLKTQGITICGDVDGKKALLTYNKTTGISNDVGTKTIVKDKSEWIAAVANHKGIIPADKWLQAQNIKDKNKDSFPALGRSNTTIASRVLRCDKCESTMGVTHGHINPVTGKKHYYYNCTLKKRSKGVRCDNKPAKAAEVDEAILITLENMFKAKSSIIDNLKAKNKARRIEMISSNRVDVINKIIEDKTKQIDNLVNKLSLDDDLTDILFKKIKGLKAEIKELEDELLTLTSDNIKLNEDEVVLDFTEKLLEKCSIIRTLDILEQQQIVDALIPLVTWNGDTEVLNIYPLGSPELELKEAESKKK | 550 | INTd |
| Segata-Pasolli | NA | PRJNA422434 | NA | NA | N684346_90_69 | SacINT | NA | human gut metagenome | stool sample from adult in China | MKEKVSERKTGAIYIRVSTDKQEELSPDAQLRLLLDYAKKDSIDVPKEYIFQDNGISGRKANKRPAFQNMIALAKSKEHPIDTIIVWKFSRFARNQEESIVYKSLLKKNNVDVVSVSEPLIDGPFGSLIERIIEWMDEYYSIRLSGEVMRGMTQNAMRGHYQSDAPIGYTSPGDKKPPVINPDTVQIPLMIKDMFLSGSTQLQIARKLNDSGYRTKRGNLWDARGVRYVLENPFYIGKSRWNYTERGRRLKPADEVIYADGNWEALWDEDTFKEIQKRLALNMRKSKSRDISAAKHWLSGLLICSSCGGTLAFGGAHNMRGFQCWKYSKGFCSESHYISTGPIEKMVLEYLEAVMHSPALSYTVISSSSVDASSKLSDLERQLQKIDAKEKRIKAAYLNEIDTLEEYKANKTALEEERRTVEKEIEELTLSDVKYSKEDLDKKMKQNISDLLRVLRDESADYIQKGNMMRNVVDHIVFNRKNTSLDVFLKLVV | 493 | INTd |
| Segata- | ERR113 | PRJEB1 | NA | NA | N687611_90_6 | RsaINT | NA | human gut metagenom | rectal swab from adult in | MKITKKQPLRPRGRSEDKRQSTKNVIRDAYINGPQKEVQIIPAKRDMEAETEKKKLRVCAYCRVSTDEDTQASSYELQVQNYTRMIRENPEWEFAGIFADEGI | 404 | INTc |

| | | | | | | | | | | | | |
|---|---|---|---|---|---|---|---|---|---|---|---|---|
| Pasolli | 6864 | 1532 | | | 8 | | | e | Isreal | SGTSVLHREHFLEMIEKCKAGEIDLIITKQVSRFARNVLDSLNYIFMLRKLDPPVGVYFETEKLNTLDKSSDMVITVLSLVAQSESEQKSNSLKWSFKRRRAQGLGIYPSWALLGYRLDDEKNWEIVEDEADIVRTIYSLYLDGYSSTQIAELLTKSGIPTVKGLSVWSSGSVLGILKNEKFCGDALCQKTVTIDFFTHKSVKNNGIEPQYFVEGHHIPIIEKNDWLLAQQIRKERRYRKRRSTHRKPRIVVKGALSGFMIVDTSWDEEYVDSLLISATQKPEPAPVIAEEDENFIVIEKE | | |
| Segata-Pasolli | ERR1136737 | PRJEB11532 | NA | NA | N687663_53_29 | Rsa2INT | NA | human gut metagenome | rectal swab from adult in Isreal | MADIQPVKNGALYIRVSTHLQEELSPDAQKRLLMEYAEAHNIIVLKEHIYIDSGISGRSARQRPQFNNMIAEAKSKEHPFDVILVWKYSRFARNQEESIVYKSMLKRENVDVISVSEPISDDPFGSLIERIIEWMDEYYSIRLSGEVSRGMAENAMRGNYQARPPLGYRIPGYRQTPVIVPEEAELIQLIFDLYTEKKMGIFEIVRYLNEHGYQTGHKKPFQRRSVTYILKNPTYIGKTIWNQHDQDHKLRDKSEWIIADGKHEPIISKEQFDKAQKRIESTYKPAYRKPTSVCHHWLSSLLKCSSCGRTLVVKRTASKKKDRMYVNFQCYGYQKGICNTNQSISAIKLEPVIMHALEDAMTSGKIHFDVLNPTTLDSSQKQQFLTRLNEIEKKEERIKRAYRDGIDTLEEYKENKSIIQTEKEMLLKKIEHIEEPALSPEEAKPIMMDRIKNVYEIITNPDIGMEEKNKAARSIIEKIVFDRATGSVNIFFYLAHCP | 498 | INTd |
| NCBI | NA | NA | NC_002656.1 | NP_075302.1 | NA | BxbINT | Bxb1 integrase | Mycobacterium phage Bxb1 | NA | MRALVVIRLSRVTDATTSPERQLESCQQLCAQRGWDVVGVAEDLDVSGAVDPFDRKRRPNLARWLAFEEQPFDVIVAYRVDRLTRSIRHLQQLVHWAEDHKKLVVSATEAHFDTTTPFAAVVIALMGTVAQMELEAIKERNRSAAHFNIRAGKYRGSLPPWGYLPTRVDGEWRLVPDPVQRERILEVYHRVVDNHEPLHLVAHDLNRRGVLSPKDYFAQLQGREPQGREWSATALKRSMISEAMLGYATLNGKTVRDDDGAPLVRAEPILTREQLEALRAELVKTSRAKPAVSTPSLLLRVLFCAVCGEPAYKFAGGGRKHPRYRCRSMGFPKHCGNGTVAMAEWDAFCEEQVLDLLGDAERLEKVWVAGSDSAVELAEVNAELVDLTSLIGSPAYRAGSPQREALDARIAALAARQEELEGLEARPSGWEWRETGQRFGDWWREQDTAAKNTWLRSMNVRLTFDVRGGLTRTIDFGDLQEYEQHLRLGSVVERLHTGMS* | 501 | INTa |
| NCBI | NA | NA | NC_002747.1 | NP_112664.1 | NA | Tp9INT | TP901-1 integrase | Lactococcus phage TP901-1 | NA | MTKKVAIYTRVSTTNQAEEGFSIDEQIDRLTKYAEAMGWQVSDTYTDAGFSGAKLERPAMQRLINDIENKAFDTVLVYKLDRLSRSVRDTLYLVKDVFTKNKIDFISLNESIDTSSAMGSLFLTILSAINEFERENIKERMTMGKLGRAKSGKSMMWTKTAFGYYHNRKTGILEIVPLQATIVEQIFTDYLSGISLTKLRDKLNESGHIGKDIPWSYRTLRQTLDNPVYCGYIKFKDSLFEGMHKPIIPYETYLKVQKELEERQQQTYERNNNPRPFQAKYMLSGMARCGYCGAPLKIVLGHKRKDGSRTMKYHCANRFPRKTKGITVYNDNKKCDSGTYDLSNLENTVIDNLIGFQENNDSLLKIINGNNQPILDTSSFKKQISQIDKKIQKNSDLYLNDFITMDELKDRTDSLQAEKKLLKAKISENKFNDSTDVFELVKTQLGSIPINELSYDNKKKIVNNLVSKVDVTADNVDIIFKFQLA* | 486 | INTd |
| NCBI | NA | NA | NC_004664.2 | NP_813744.2 | NA | Bt1INT | PhiBT integrase | Streptomyces virus phiBT1 | NA | MSPFIAPDVPEHLLDTVRVFLYARQSKGRSDGSDVSTEAQLAAGRALVASRNAQGGARWVVAGEFVDVGRSGWDPNVTRADFERMMGEVRAGEGDVVVVNELSRLTRKGAHDALEIDNELKKHGVRFMSVLEPFLDTSTPIGVAIFALIAALAKQDSDLKAERLKGAKDEIAALGGVHSSSAPFGMRAVRKKVDNLVISVLEPDEDNPDHVELVERMAKMSFEGVSDNAIATTFEKEKIPSPGMAERRATEKRLASIKARRLNGAEKPIMWRAQTVRWILNHPAIGGFAFERVKHGKAHINVIRRDPGGKPLTPHTGILSGSKWLELQEKRSGKNLSDRKPGAEVEPTLLSGWRFLGCRICGGSMGQSQGGRKRNGDLAEGNYMCANPKGHGGLSVKRS | 595 | INTa |

| | | | | | | | | | | | | |
|---|---|---|---|---|---|---|---|---|---|---|---|---|
| | | | | | | | | | | ELDEFVASKVWARLRTADMEDEHDQAWIAAAAERFALQHDLAGVADERREQQAHLDNVRRSIKDLQADRKAGLYVGREELETWRSTVLQYRSYEAECTTRLAELDEKMNGSTRVPSEWFSGEDPTAEGGIWASWDVYERREFLSFFLDSVMVDRGRHPETKKYIPLKDRVTLKWAELLKEEDEASEATERELAAL* | | |
| NCBI | NA | NA | NC_011658.1 | WP_000286206.1 | NA | BceINT | NA | Bacillus cereus AH187 | NA | MYPYDVPDYAGSYRPESLDVCIYLRKSRKDVEEERRAIEEGSSYNALERHRKRLFAIAKAENHNIIDIFEEVASGESIQERPQMQQLLRKLEGNEIDGVLVIDLDRLGRGDMLDAGMIDRAFRYSSTKIITPTDVYDPDDESWELVFGIKSLISRQELKSITKRLQNGRIDSVKEGKHIGKKPPYGYLKDENLRLYPDPEKAWIVKKIFELMCDGKGRQMIAAELDRLGIDPPVTKRGAWDSSTITSIIKNEVYTGVIVWGKFKHKKRNGKYTRHKNPQEKWIMYENAHEPIISKELFDAANEAHSSRHKPAVITSKKLTNPLAGILKCKLCGYTMLIQTRKDRPHNYLRCNNPACKGKQKQSVFNLVEEKLLYSLQQIVDEYQAQKVEEVEIDDSKLISFKEKAIISKEKELKELQAQKGNLHDLLEQGIYTVEIFLERQKNLVERITSIENDIEVLQKEIETEQIKEHNKTEFIPALKTVIESYHKTTNIELKNQLLKTILSTVTYYRHPDWKTNEFEIQVYFKIS* | 529 | INTc |
| NCBI | NA | NA | NC_009674.1 | WP_012095429.1 | NA | BcyINT | NA | Bacillus cytotoxicus NVH 391-98 | NA | MYPYDVPDYAGSAVGIYIRVSTQEQASEGHSIESQKKKLASYCEIQGWDDYRFYIEEGISGKNTNRPKLKLLMEHIEKGKINILLVYRLDRLTRSVIDLHKLLNFLQEHGCAFKSATETYDTTTANGRMSMGIVSLLAQWETENMSERIKLNLEHKVLVEGERVGAIPYGFDLSDDEKLVKNEKSAILLDMVERVENGWSVNRIVNYLNLTNNDRNWSPNGVLRLLRNPALYGATRWNDKIAENTHEGIISKERFNRLQQILADRSIHHRRDVKGTYIFQGVLRCPVCDQTLSVNRFIKKRKDGTEYCGVLYRCQPCIKQNKYNLAIGEARFLKALNEYMSTVEFQTVEDEVIPKKSEREMLESQLQQIARKREKYQKAWASDLMSDDEFEKLMVETRETYDECKQKLESCEDPIKIDETYLKEIVYMFHQTFNDLESEKQKEFISKFIRTIRYTVKEQQPIRPDKSKTGKGKQKVIITEVEFYQS* | 487 | INTd |
| NCBI | NA | NA | NC_017353.1 | WP_014533238.1 | NA | SluINT | NA | Staphylococcus lugdunensis N920143 | NA | MYPYDVPDYAGSKVAIYTRVSSAEQANEGYSIHEQKKKLISYCEIHDWNEYKVFTDAGISGGSMKRPALQKLMKHLSSFDLVLVYKLDRLTRNVRDLLDMLEEFEQYNVSFKSATEVFDTTSAIGKLFITMVGAMAEWERETIRERSLFGSRAAVREGNYIREAPFCYDNIEGKLHPNEYAKVIDLIVSMFKKGISANEIARRLNSSKVHVPNKKSWNRNSLIRLMRSPVLRGHTKYGDMLIENTHEPVLSEHDYNAINNAISSKTHKSKVKHHAIFRGALVCPQCNRRLHLYAGTVKDRKGYKYDVRRYKCETCSKNKDVKNVSFNESEVENKFVNLLKSYELNKFHIRKVEPVKKIEYDIDKINKQKINYTRSWSLGYIEDDEYFELMEEINATKKMIEEQTTENKQSVSKEQIQSINNFILKGWEELTIKDKEELILSTVDKIEFNFIPKDKKHKTNTLDINNIHFKFS* | 473 | INTd |

**Supplementary Table 2.** Linker sequences

| Description | Sequence (5'-3') | Amino acid sequence |
|---|---|---|
| A - P2A | GGAAGCGGAGCTACTAACTTCAGCCT GCTGAAGCAGGCTGGCGACGTGGAGG AGAACCCTGGACCT | GSGATNFSLLKQAGDVEENPGP |
| B - (GGGS)3 | GGGGGAGGAGGTTCTGGAGGCGGAGG CTCCGGAGGCGGAGGGTCA | GGGGSGGGGSGGGGS |
| C - GGGGS | GGAGGTGGCGGGAGC | GGGGS |
| D - PAPAP | CCCGCACCAGCGCCT | PAPAP |
| E - (EAAAK)3 | GAGGCAGCTGCCAAGGAAGCCGCT GCCAAGGAGGCGGCCGCAAAG | EAAAKEAAAKEAAAK |
| F - XTEN | AGTGGGAGCGAGACCCCTGGGACT AGCGAGTCAGCTACACCCGAAAGC | SGSETPGTSESATPES |
| G - (GGS)6 | GGGGGGTCAGGTGGATCCGGCGG AAGTGGCGGATCCGGTGGATCTGG CGGCAGT | GGSGGSGGSGGSGGSGGS |
| H - EAAAK | GAAGCTGCTGCTAAG | EAAAK |

**Supplementary Table 3.** Sequences of RT mutants

| Description | Forward Sequence (5'-3') |
|---|---|
| RT_mut_L139P | ttgagcgggCCCccaccgt |
| RT_mut_E562Q | cagcgggctCAGctgatagca |
| RT_mut_D653N | cggatggctAACcaagcggcc |
| RT(1-478)_Sto7d fusion [MMulv sequence, Sto7d sequence] | atgactcactatcaggccttgcttttggacacggaccgggtccagttcggaccggtggtagccctgaacccggctacgctgctccc actgcctgaggaagggctgcaacacaactgccttgatGGGACAGGTGGCGGTGGTGTCACCGTCAA GTTCAAGTACAAGGGTGAGGAACTTGAAGTTGATATTAGCAAAATCAAGAAG GTTTGGCGCGTTGGTAAAATGATATCTTTTACTTATGACGACAACGGCAAGAC AGGTAGAGGGGCAGTGTCTGAGAAAGACGCCCCCAAGGAGCTGTTGCAAATG TTGGAAAAGTCTGGGAAAAAGtctggcggctcaaaaagaaccgccgacggcagcgaattcgagcccaaga agaagaggaaagtc |

**Supplementary Table 4.** Sequences of atgRNAs, sgRNAs and nicking guides. Spacers are labeled in blue, RT regions in green, AttB sites in red, and PBS in orange. Unless otherwise denoted, the AttB is for Bxb1.

| Description | Sequence (5’-3’) |
| --- | --- |
| ACTB N-term PBS 13 RT 29 AttB 46 atgRNA | GCTATTCTCGCAGCTCACCAgttttagagctagaaatagcaagttaaaataaggctagtccgttatcaacttgaaaaagtggcaccgagtcggtgcGACGAGCGCGGCGATATCATCATCCATGGccggatgatcctgacgacggagaccgccgtcgtcgacaagccggccTGAGCTGCGAGAA |
| ACTB N-term PBS 13 RT 29 AttB 46 atgRNA with v2 scaffold | GCTATTCTCGCAGCTCACCAgtttgagagctatgctggaaacagcatagcaagttcaaataaggctagtccgttatcaacttgaaaaagtggcaccgagtcggtgcGACGAGCGCGGCGATATCATCATCCATGGccggatgatcctgacgacggagaccgccgtcgtcgacaagccggccTGAGCTGCGAGAA |
| ACTB N-term PBS_13_RT_29_with TP901-1 minimal AttB f atgRNA | GCTATTCTCGCAGCTCACCAgttttagagctagaaatagcaagttaaaataaggctagtccgttatcaacttgaaaaagtggcaccgagtcggtgcGAGTCGGTGCGACGAGCGCGGCGATATCATCATCCATGGcacaattaacatctcaatcaaggtaaaTGCTTGAGCTGCGAGAA |
| ACTB N-term PBS_13_RT_29_with TP901-1 minimal AttB rc atgRNA | GCTATTCTCGCAGCTCACCAgttttagagctagaaatagcaagttaaaataaggctagtccgttatcaacttgaaaaagtggcaccgagtcggtgcGAGTCGGTGCGACGAGCGCGGCGATATCATCATCCATGGagcatttaccttgattgagatgttaattgtgTGAGCTGCGAGAA |
| ACTB N-term PBS_13_RT_29_with PhiBT1 minimal AttB f atgRNA | GCTATTCTCGCAGCTCACCAgttttagagctagaaatagcaagttaaaataaggctagtccgttatcaacttgaaaaagtggcaccgagtcggtgcGAGTCGGTGCGACGAGCGCGGCGATATCATCATCCATGGcaggtttttgacgaaagtgatccagatgatccagTGAGCTGCGAGAA |
| ACTB N-term PBS_13_RT_29_with PhiBT1 minimal AttB rc atgRNA | GCTATTCTCGCAGCTCACCAgttttagagctagaaatagcaagttaaaataaggctagtccgttatcaacttgaaaaagtggcaccgagtcggtgcGAGTCGGTGCGACGAGCGCGGCGATATCATCATCCATGGctggatcatctggatcactttcgtcaaaaacctgTGAGCTGCGAGAA |
| ACTB N-term Nicking guide 1 +48 guide | GAAGCCGGCCTTGCACATGCgttttagagctagaaatagcaagttaaaataaggctagtccgttatcaacttgaaaaagtggcaccgagtcggtgc |
| ACTB N-term PBS_18_RT_16_with_Lox71_Cre atgRNA | GAAGCCGGCCTTGCACATGCgttttagagctagaaatagcaagttaaaataaggctagtccgttatcaacttgaaaaagtggcaccgagtcggtgcATATCATCATCCATGGtaccgttcgtatagcatacattatacgaagttatTGAGCTGCGAGAATAGCC |
| ACTB N-term PBS_13_RT_29_with_Lox71_Cre atgRNA | GAAGCCGGCCTTGCACATGCgttttagagctagaaatagcaagttaaaataaggctagtccgttatcaacttgaaaaagtggcaccgagtcggtgcGACGAGCGCGGCGATATCATCATCCATGGtaccgttcgtatagcatacattatacgaagttatTGAGCTGCGAGAA |
| ACTB N-term PBS 13 RT 34 atgRNA | GCTATTCTCGCAGCTCACCAgttttagagctagaaatagcaagttaaaataaggctagtccgttatcaacttgaaaaagtggcaccgagtcggtgcTCGACGACGAGCGCGGCGATATCATCATCCATGGccggatgatcctgacgacggagaccgccgtcgtcgacaagccggccTGAGCTGCGAGAA |
| ACTB N-term PBS 13 RT 26 atgRNA | GCTATTCTCGCAGCTCACCAgttttagagctagaaatagcaagttaaaataaggctagtccgttatcaacttgaaaaagtggcaccgagtcggtgcGAGCGCGGCGATATCATCATCCATGGccggatgatcctgacgacggagaccgccgtcgtcgacaagccggccTGAGCTGCGAGAA |

| | |
|---|---|
| ACTB N-term PBS 13 RT 23 atgRNA | GCTATTCTCGCAGCTCACCAgttttagagctagaaatagcaagttaaaataaggctagtccgttatcaacttgaaaaagtggcaccgagtcggtgcCGCGGCGATATCATCATCCATGGccggatgatcctgacgacggagaccgccgtcgtcgacaagccggccTGAGCTGCGAGAA |
| ACTB N-term PBS 13 RT 20 atgRNA | GCTATTCTCGCAGCTCACCAgttttagagctagaaatagcaagttaaaataaggctagtccgttatcaacttgaaaaagtggcaccgagtcggtgcGGCGATATCATCATCCATGGccggatgatcctgacgacggagaccgccgtcgtcgacaagccggccTGAGCTGCGAGAA |
| ACTB N-term PBS 13 RT 16 atgRNA | GCTATTCTCGCAGCTCACCAgttttagagctagaaatagcaagttaaaataaggctagtccgttatcaacttgaaaaagtggcaccgagtcggtgcATATCATCATCCATGGccggatgatcctgacgacggagaccgccgtcgtcgacaagccggccTGAGCTGCGAGAA |
| ACTB N-term PBS 18 RT 34 atgRNA | GCTATTCTCGCAGCTCACCAgttttagagctagaaatagcaagttaaaataaggctagtccgttatcaacttgaaaaagtggcaccgagtcggtgcTCGACGACGAGCGCGGCGATATCATCATCCATGGccggatgatcctgacgacggagaccgccgtcgtcgacaagccggccTGAGCTGCGAGAATAGCC |
| ACTB N-term PBS 18 RT 29 atgRNA | GCTATTCTCGCAGCTCACCAgttttagagctagaaatagcaagttaaaataaggctagtccgttatcaacttgaaaaagtggcaccgagtcggtgcGACGAGCGCGGCGATATCATCATCCATGGccggatgatcctgacgacggagaccgccgtcgtcgacaagccggccTGAGCTGCGAGAATAGCC |
| ACTB N-term PBS 18 RT 16 atgRNA | GCTATTCTCGCAGCTCACCAgttttagagctagaaatagcaagttaaaataaggctagtccgttatcaacttgaaaaagtggcaccgagtcggtgcATATCATCATCCATGGccggatgatcctgacgacggagaccgccgtcgtcgacaagccggccTGAGCTGCGAGAATAGCC |
| LMNB1 N-term PBS 13 RT 39 atgRNA | GCTGTCTCCGCCGCCCGCCAgttttagagctagaaatagcaagttaaaataaggctagtccgttatcaacttgaaaaagtggcaccgagtcggtgcCTGCCCATCCGCGGCGGCACGGGGGTCGCAGTCGCCATGccggatgatcctgacgacggagaccgccgtcgtcgacaagccggccCGGGCGGCGGAGA |
| LMNB1 N-term PBS 13 RT 34 atgRNA | GCTGTCTCCGCCGCCCGCCAgttttagagctagaaatagcaagttaaaataaggctagtccgttatcaacttgaaaaagtggcaccgagtcggtgcCATCCGCGGCGGCACGGGGGTCGCAGTCGCCATGccggatgatcctgacgacggagaccgccgtcgtcgacaagccggccCGGGCGGCGGAGA |
| LMNB1 N-term PBS 13 RT 29 atgRNA | GCTGTCTCCGCCGCCCGCCAgttttagagctagaaatagcaagttaaaataaggctagtccgttatcaacttgaaaaagtggcaccgagtcggtgcGCGGCGGCACGGGGGTCGCAGTCGCCATGccggatgatcctgacgacggagaccgccgtcgtcgacaagccggccCGGGCGGCGGAGA |
| LMNB1 N-term PBS 13 RT 24 atgRNA | GCTGTCTCCGCCGCCCGCCAgttttagagctagaaatagcaagttaaaataaggctagtccgttatcaacttgaaaaagtggcaccgagtcggtgcGGCACGGGGGTCGCAGTCGCCATGccggatgatcctgacgacggagaccgccgtcgtcgacaagccggccCGGGCGGCGGAGA |
| LMNB1 N-term PBS 13 RT 19 atgRNA | GCTGTCTCCGCCGCCCGCCAgttttagagctagaaatagcaagttaaaataaggctagtccgttatcaacttgaaaaagtggcaccgagtcggtgcGGGGGTCGCAGTCGCCATGccggatgatcctgacgacggagaccgccgtcgtcgacaagccggccCGGGCGGCGGAGA |
| LMNB1 N-term PBS 18 RT 39 atgRNA | GCTGTCTCCGCCGCCCGCCAgttttagagctagaaatagcaagttaaaataaggctagtccgttatcaacttgaaaaagtggcaccgagtcggtgcCTGCCCATCCGCGGCGGCACGGGGGTCGCAGTCGCCATGccggatgatcctgacgacggagaccgccgtcgtcgacaagccggccCGGGCGGCGGAGACAGCG |
| LMNB1 N-term PBS 18 RT 34 atgRNA | GCTGTCTCCGCCGCCCGCCAgttttagagctagaaatagcaagttaaaataaggctagtccgttatcaacttgaaaaagtggcaccgagtcggtgcCATCCGCGGCGGCACGGGGGTCGCAGTC |

| | |
|---|---|
| | GCCATGccggatgatcctgacgacggagaccgccgtcgtcgacaagccggccCGGGCGGCGGAGACAGCG |
| LMNB1 N-term PBS 18 RT 29 atgRNA | GCTGTCTCCGCCGCCCGCCAgttttagagctagaaatagcaagttaaaataaggctagtccgttatcaacttgaaaaagtggcaccgagtcggtgcGCGGCGGCACGGGGGTCGCAGTCGCCATGccggatgatcctgacgacggagaccgccgtcgtcgacaagccggccCGGGCGGCGGAGACAGCG |
| LMNB1 N-term PBS 18 RT 24 atgRNA | GCTGTCTCCGCCGCCCGCCAgttttagagctagaaatagcaagttaaaataaggctagtccgttatcaacttgaaaaagtggcaccgagtcggtgcGGCACGGGGGTCGCAGTCGCCATGccggatgatcctgacgacggagaccgccgtcgtcgacaagccggccCGGGCGGCGGAGACAGCG |
| LMNB1 N-term PBS 18 RT 19 atgRNA | GCTGTCTCCGCCGCCCGCCAgttttagagctagaaatagcaagttaaaataaggctagtccgttatcaacttgaaaaagtggcaccgagtcggtgcGGGGGTCGCAGTCGCCATGccggatgatcctgacgacggagaccgccgtcgtcgacaagccggccCGGGCGGCGGAGACAGCG |
| LMNB1 N-term Nicking guide 1 +46 | GCGTGGTGGGGCCGCCAGCGgttttagagctagaaatagcaagttaaaataaggctagtccgttatcaacttgaaaaagtggcaccgagtcggtgc |
| ACTB N-term PBS 13 RT 29 AttB 42 atgRNA | GCTATTCTCGCAGCTCACCAgttttagagctagaaatagcaagttaaaataaggctagtccgttatcaacttgaaaaagtggcaccgagtcggtgcGACGAGCGCGGCGATATCATCATCCATGGggatgatcctgacgacggagaccgccgtcgtcgacaagccggTGAGCTGCGAGAA |
| ACTB N-term PBS 13 RT 29 AttB 40 atgRNA | GCTATTCTCGCAGCTCACCAgttttagagctagaaatagcaagttaaaataaggctagtccgttatcaacttgaaaaagtggcaccgagtcggtgcGACGAGCGCGGCGATATCATCATCCATGGgatgatcctgacgacggagaccgccgtcgtcgacaagccgTGAGCTGCGAGAA |
| ACTB N-term PBS 13 RT 29 AttB 38 atgRNA | GCTATTCTCGCAGCTCACCAgttttagagctagaaatagcaagttaaaataaggctagtccgttatcaacttgaaaaagtggcaccgagtcggtgcGACGAGCGCGGCGATATCATCATCCATGGatgatcctgacgacggagaccgccgtcgtcgacaagccTGAGCTGCGAGAA |
| ACTB N-term PBS 13 RT 29 AttB 36 atgRNA | GCTATTCTCGCAGCTCACCAgttttagagctagaaatagcaagttaaaataaggctagtccgttatcaacttgaaaaagtggcaccgagtcggtgcGACGAGCGCGGCGATATCATCATCCATGGtgatcctgacgacggagaccgccgtcgtcgacaagcTGAGCTGCGAGAA |
| LMNB1 N-term PBS 13 RT 29 AttB 44 atgRNA v2 | GCTGTCTCCGCCGCCCGCCAgttttagagctagaaatagcaagttaaaataaggctagtccgttatcaacttgaaaaagtggcaccgagtcggtgcGCGGCGGCACGGGGGTCGCAGTCGCCATGcggatgatcctgacgacggagaccgccgtcgtcgacaagccggcCGGGCGGCGGAGA |
| LMNB1 N-term PBS 13 RT 29 AttB 42 atgRNA v2 | GCTGTCTCCGCCGCCCGCCAgttttagagctagaaatagcaagttaaaataaggctagtccgttatcaacttgaaaaagtggcaccgagtcggtgcGCGGCGGCACGGGGGTCGCAGTCGCCATGggatgatcctgacgacggagaccgccgtcgtcgacaagccggCGGGCGGCGGAGA |
| LMNB1 N-term PBS 13 RT 29 AttB 40 atgRNA v2 | GCTGTCTCCGCCGCCCGCCAgttttagagctagaaatagcaagttaaaataaggctagtccgttatcaacttgaaaaagtggcaccgagtcggtgcGCGGCGGCACGGGGGTCGCAGTCGCCATGgatgatcctgacgacggagaccgccgtcgtcgacaagccgCGGGCGGCGGAGA |
| LMNB1 N-term PBS 13 RT 29 AttB 38 atgRNA v2 | GCTGTCTCCGCCGCCCGCCAgttttagagctagaaatagcaagttaaaataaggctagtccgttatcaacttgaaaaagtggcaccgagtcggtgcGCGGCGGCACGGGGGTCGCAGTCGCCATGatgatcctgacgacggagaccgccgtcgtcgacaagccCGGGCGGCGGAGA |
| NOLC1 N-term PBS 18 RT 29 AttB 46 atgRNA | GCGTATTGCCTGGAGGATGGgttttagagctagaaatagcaagttaaaataaggctagtccgttatcaacttgaaaaagtggcaccgagtcggtgcGAACCACGCGGCGAATGCCGGCGTCCGCCccggatgatcctgacgacggagaccgccgtcgtcgacaagccggccTCCTCCAGGCAATACGCG |
| NOLC1 N-term PBS 13 | GCGTATTGCCTGGAGGATGGgttttagagctagaaatagcaagttaaaataaggctagtccgttat |

| | |
|---|---|
| RT 29 AttB 46 atgRNA | caacttgaaaaagtggcaccgagtcggtgcGAACCACGCGGCGAATGCCGGCGTCCGCCccggatgatcctgacgacggagaccgccgtcgtcgacaagccggccTCCTCCAGGCAAT |
| NOLC1 N-term PBS 13 RT 29 AttB 44 atgRNA | GCGTATTGCCTGGAGGATGGgttttagagctagaaatagcaagttaaaataaggctagtccgttatcaacttgaaaaagtggcaccgagtcggtgcGAACCACGCGGCGAATGCCGGCGTCCGCCcggatgatcctgacgacggagaccgccgtcgtcgacaagccggcTCCTCCAGGCAAT |
| NOLC1 N-term PBS 13 RT 29 AttB 42 atgRNA | GCGTATTGCCTGGAGGATGGgttttagagctagaaatagcaagttaaaataaggctagtccgttatcaacttgaaaaagtggcaccgagtcggtgcGAACCACGCGGCGAATGCCGGCGTCCGCCggatgatcctgacgacggagaccgccgtcgtcgacaagccggTCCTCCAGGCAAT |
| NOLC1 N-term PBS 13 RT 29 AttB 40 atgRNA | GCGTATTGCCTGGAGGATGGgttttagagctagaaatagcaagttaaaataaggctagtccgttatcaacttgaaaaagtggcaccgagtcggtgcGAACCACGCGGCGAATGCCGGCGTCCGCCgatgatcctgacgacggagaccgccgtcgtcgacaagccgTCCTCCAGGCAAT |
| NOLC1 N-term PBS 13 RT 29 AttB 38 atgRNA | GCGTATTGCCTGGAGGATGGgttttagagctagaaatagcaagttaaaataaggctagtccgttatcaacttgaaaaagtggcaccgagtcggtgcGAACCACGCGGCGAATGCCGGCGTCCGCCatgatcctgacgacggagaccgccgtcgtcgacaagccTCCTCCAGGCAAT |
| NOLC1 nicking guide -43 | GAGCCGAGCACGAGGGATACgttttagagctagaaatagcaagttaaaataaggctagtccgttatcaacttgaaaaagtggcaccgagtcggtgc |
| ACTB N-term PBS 13 RT 20 AttB 38 atgRNA | GCTATTCTCGCAGCTCACCAgttttagagctagaaatagcaagttaaaataaggctagtccgttatcaacttgaaaaagtggcaccgagtcggtgcGGCGATATCATCATCCATGGatgatcctgacgacggagaccgccgtcgtcgacaagccTGAGCTGCGAGAA |
| ACTB N-term PBS 13 RT 15 AttB 38 atgRNA | GCTATTCTCGCAGCTCACCAgttttagagctagaaatagcaagttaaaataaggctagtccgttatcaacttgaaaaagtggcaccgagtcggtgcTATCATCATCCATGGatgatcctgacgacggagaccgccgtcgtcgacaagccTGAGCTGCGAGAA |
| ACTB N-term PBS 13 RT 10 AttB 38 atgRNA | GCTATTCTCGCAGCTCACCAgttttagagctagaaatagcaagttaaaataaggctagtccgttatcaacttgaaaaagtggcaccgagtcggtgcTCATCCATGGatgatcctgacgacggagaccgccgtcgtcgacaagccTGAGCTGCGAGAA |
| ACTB N-term PBS 9 RT 20 AttB 38 atgRNA | GCTATTCTCGCAGCTCACCAgttttagagctagaaatagcaagttaaaataaggctagtccgttatcaacttgaaaaagtggcaccgagtcggtgcGGCGATATCATCATCCATGGatgatcctgacgacggagaccgccgtcgtcgacaagccTGAGCTGCG |
| ACTB N-term PBS 9 RT 15 AttB 38 atgRNA | GCTATTCTCGCAGCTCACCAgttttagagctagaaatagcaagttaaaataaggctagtccgttatcaacttgaaaaagtggcaccgagtcggtgcTATCATCATCCATGGatgatcctgacgacggagaccgccgtcgtcgacaagccTGAGCTGCG |
| ACTB N-term PBS 9 RT 10 AttB 38 atgRNA | GCTATTCTCGCAGCTCACCAgttttagagctagaaatagcaagttaaaataaggctagtccgttatcaacttgaaaaagtggcaccgagtcggtgcTCATCCATGGatgatcctgacgacggagaccgccgtcgtcgacaagccTGAGCTGCG |
| LMNB1 N-term PBS 13 RT 20 AttB 38 atgRNA | GCTGTCTCCGCCGCCCGCCAgttttagagctagaaatagcaagttaaaataaggctagtccgttatcaacttgaaaaagtggcaccgagtcggtgcCGGGGGTCGCAGTCGCCATGatgatcctgacgacggagaccgccgtcgtcgacaagccCGGGCGGCGGAGA |
| LMNB1 N-term PBS 13 RT 15 AttB 38 atgRNA | GCTGTCTCCGCCGCCCGCCAgttttagagctagaaatagcaagttaaaataaggctagtccgttatcaacttgaaaaagtggcaccgagtcggtgcGTCGCAGTCGCCATGatgatcctgacgacggagaccgccgtcgtcgacaagccCGGGCGGCGGAGA |
| LMNB1 N-term PBS 13 RT 10 AttB 38 atgRNA | GCTGTCTCCGCCGCCCGCCAgttttagagctagaaatagcaagttaaaataaggctagtccgttatcaacttgaaaaagtggcaccgagtcggtgcAGTCGCCATGatgatcctgacgacggagaccgccgtcgtcgacaagccCGGGCGGCGGAGA |
| LMNB1 N-term PBS 9 | GCTGTCTCCGCCGCCCGCCAgttttagagctagaaatagcaagttaaaataaggctagtccgttat |

| | |
|---|---|
| RT 20 AttB 38 atgRNA | caacttgaaaaagtggcaccgagtcggtgcCGGGGGTCGCAGTCGCCATGatgatcctgacgacggagaccgccgtcgtcgacaagccCGGGCGGCG |
| LMNB1 N-term PBS 9 RT 15 AttB 38 atgRNA | GCTGTCTCCGCCGCCCGCCAgttttagagctagaaatagcaagttaaaataaggctagtccgttatcaacttgaaaaagtggcaccgagtcggtgcGTCGCAGTCGCCATGatgatcctgacgacggagaccgccgtcgtcgacaagccCGGGCGGCG |
| LMNB1 N-term PBS 9 RT 10 AttB 38 atgRNA | GCTGTCTCCGCCGCCCGCCAgttttagagctagaaatagcaagttaaaataaggctagtccgttatcaacttgaaaaagtggcaccgagtcggtgcAGTCGCCATGatgatcctgacgacggagaccgccgtcgtcgacaagccCGGGCGGCG |
| SUPT16H N-term PBS 13 RT 24 Bxb1-GT_Initial length | GAGAAGCGGCGTCCGGGGCTAgttttagagctagaaatagcaagttaaaataaggctagtccgttatcaacttgaaaaagtggcaccgagtcggtgcTCTTTGTCCAGAGTCACAGCCATAccggatgatcctgacgacggagaccgccgtcgtcgacaagccggccCCCCGGACGCCGC |
| SRRM2 N-term PBS 13 RT 24 Bxb1 Initial length | GGGCACGGGGCCATGTACAAgttttagagctagaaatagcaagttaaaataaggctagtccgttatcaacttgaaaaagtggcaccgagtcggtgcGGCGTCGGCAGCCCGATCCCGTTGccggatgatcctgacgacggagaccgccgtcgtcgacaagccggccTACATGGCCCCGT |
| DEPDC4 N-term PBS 18 RT 24 Bxb1 Initial length | GTGTCAGGTGGGGCGGGGCTAgttttagagctagaaatagcaagttaaaataaggctagtccgttatcaacttgaaaaagtggcaccgagtcggtgcGCTGGCTCCTCCCCTGGCACCATAccggatgatcctgacgacggagaccgccgtcgtcgacaagccggccCCCCGCCCCACCTGACAC |
| NES N-term PBS 13 RT 29 Bxb1 Initial length | GAGTGGGTCAGACGAGCAGGAgttttagagctagaaatagcaagttaaaataaggctagtccgttatcaacttgaaaaagtggcaccgagtcggtgcCGACTCCTCCCCCATGCAGCCCTCCATCccggatgatcctgacgacggagaccgccgtcgtcgacaagccggccTGCTCGTCTGACC |
| SUPT16H nicking guide -53 | GCAGCCACCCGCTCTCGGCCCgttttagagctagaaatagcaagttaaaataaggctagtccgttatcaacttgaaaaagtggcaccgagtcggtgc |
| SRRM2 N-term nicking guide 1 +87 | GTGTAGTCAGGCCGCTCACCCgttttagagctagaaatagcaagttaaaataaggctagtccgttatcaacttgaaaaagtggcaccgagtcggtgc |
| DEPDC4 N-term Nicking guide 1 +59 | GCTGACAAGTCTACGGAACCTgttttagagctagaaatagcaagttaaaataaggctagtccgttatcaacttgaaaaagtggcaccgagtcggtgc |
| NES N-term Nicking guide 2 +79 | GCTCCTCCAGCGCCTTGACCgttttagagctagaaatagcaagttaaaataaggctagtccgttatcaacttgaaaaagtggcaccgagtcggtgc |
| HITI_ACTB_guide | GCTATTCTCGCAGCTCACCAgttttagagctagaaatagcaagttaaaataaggctagtccgttatcaacttgaaaaagtggcaccgagtcggtgc |
| HITI_SUPTH16_guide | AGAAGCGGCGTCCGGGGCTAgttttagagctagaaatagcaagttaaaataaggctagtccgttatcaacttgaaaaagtggcaccgagtcggtgc |
| HITI_SRRM2_guide | GGGCACGGGGCCATGTACAAgttttagagctagaaatagcaagttaaaataaggctagtccgttatcaacttgaaaaagtggcaccgagtcggtgc |
| HITI_NOLC1_guide | GCGTATTGCCTGGAGGATGGgttttagagctagaaatagcaagttaaaataaggctagtccgttatcaacttgaaaaagtggcaccgagtcggtgc |
| HITI_DEPDC4_guide | TGTCAGGTGGGGCGGGGCTAgttttagagctagaaatagcaagttaaaataaggctagtccgttatcaacttgaaaaagtggcaccgagtcggtgc |
| HITI_NES_guide | AGTGGGTCAGACGAGCAGGAgttttagagctagaaatagcaagttaaaataaggctagtccgtta |

| | |
|---|---|
| | tcaacttgaaaaagtggcaccgagtcggtgc |
| HITI_LMNB1_guide | GCTGTCTCCGCCGCCCGCCAgttttagagctagaaatagcaagttaaaataaggctagtccgttatcaacttgaaaaagtggcaccgagtcggtgc |
| HDR Cas9 ACTB guide | GCTATTCTCGCAGCTCACCAgttttagagctagaaatagcaagttaaaataaggctagtccgttatcaacttgaaaaagtggcaccgagtcggtgc |
| HDR Cas9 LMNB1 guide | GGGGTCGCAGTCGCCATGGCgttttagagctagaaatagcaagttaaaataaggctagtccgttatcaacttgaaaaagtggcaccgagtcggtgc |
| ACTB N-term PBS 13 RT 29 AttB original length atgRNAs for dinucleotides | GCTATTCTCGCAGCTCACCAgttttagagctagaaatagcaagttaaaataaggctagtccgttatcaacttgaaaaagtggcaccgagtcggtgcGACGAGCGCGGCGATATCATCATCCATGGccggatgatcctgacgacggagXXcgccgtcgtcgacaagccggccTGAGCTGCGAGAA<br><br>XX: CG, GC, AT, TA, GG, TT, GA, AG, CC, TC, CT, AA, TG, GT, CA, AC |
| ACTB N-term PBS 13 RT 29 atgRNA with AttB 46 GT for fusion | GCTATTCTCGCAGCTCACCAgttttagagctagaaatagcaagttaaaataaggctagtccgttatcaacttgaaaaagtggcaccgagtcggtgcGACGAGCGCGGCGATATCATCATCCATGccggatgatcctgacgacggagACcgccgtcgtcgacaagccggccTGAGCTGCGAGAA |
| ACTB N-term PBS 13 RT 29 atgRNA with AttB 46 CT for multiplexing | GCTATTCTCGCAGCTCACCAgttttagagctagaaatagcaagttaaaataaggctagtccgttatcaacttgaaaaagtggcaccgagtcggtgcGACGAGCGCGGCGATATCATCATCCATGccggatgatcctgacgacggagAGcgccgtcgtcgacaagccggccTGAGCTGCGAGAA |
| NOLC1 N-term PBS 18 RT 29 atgRNA with AttB 46 GA for multiplexing | GCGTATTGCCTGGAGGATGGgttttagagctagaaatagcaagttaaaataaggctagtccgttatcaacttgaaaaagtggcaccgagtcggtgcGAACCACGCGGCGAATGCCGGCGTCCGCCccggatgatcctgacgacggagTCcgccgtcgtcgacaagccggccTCCTCCAGGCAATACGCG |
| LMNB1 N-term PBS 18 RT 29 atgRNA with AttB 46 AG for multiplexing | GCTGTCTCCGCCGCCCGCCAgttttagagctagaaatagcaagttaaaataaggctagtccgttatcaacttgaaaaagtggcaccgagtcggtgcGCGGCGGCACGGGGGTCGCAGTCGCCATGccggatgatcctgacgacggagCTcgccgtcgtcgacaagccggccCGGGCGGCGGAGACAGCG |
| EMX1 Cas9 guide 1 | GTCACCTCCAATGACTAGGGgttttagagctagaaatagcaagttaaaataaggctagtccgttatcaacttgaaaaagtggcaccgagtcggtgc |
| EMX1 Cas9 guide 2 | GGGCAACCACAAACCCACGAgttttagagctagaaatagcaagttaaaataaggctagtccgttatcaacttgaaaaagtggcaccgagtcggtgc |
| ACTB N-term PBS 13 RT 29 AttB 56 GA atgRNA | GCTATTCTCGCAGCTCACCAgttttagagctagaaatagcaagttaaaataaggctagtccgttatcaacttgaaaaagtggcaccgagtcggtgcGACGAGCGCGGCGATATCATCATCCATGGctatgccggatgatcctgacgacggagtccgccgtcgtcgacaagccggccctagcTGAGCTGCGAGAA |
| ACTB N-term PBS 13 RT 29 AttB 51 GA atgRNA | GCTATTCTCGCAGCTCACCAgttttagagctagaaatagcaagttaaaataaggctagtccgttatcaacttgaaaaagtggcaccgagtcggtgcGACGAGCGCGGCGATATCATCATCCATGGtgccggatgatcctgacgacggagtccgccgtcgtcgacaagccggccctaTGAGCTGCGAGAA |
| ACTB N-term PBS 13 RT 29 AttB 46 GA atgRNA | GCTATTCTCGCAGCTCACCAgttttagagctagaaatagcaagttaaaataaggctagtccgttatcaacttgaaaaagtggcaccgagtcggtgcGACGAGCGCGGCGATATCATCATCCATGGccggatgatcctgacgacggagtccgccgtcgtcgacaagccggccTGAGCTGCGAGAA |
| ACTB N-term PBS 13 RT 29 AttB 41 GA atgRNA | GCTATTCTCGCAGCTCACCAgttttagagctagaaatagcaagttaaaataaggctagtccgttatcaacttgaaaaagtggcaccgagtcggtgcGACGAGCGCGGCGATATCATCATCCATGGggatgatcctgacgacggagtccgccgtcgtcgacaagccgTGAGCTGCGAGAA |

| | |
|---|---|
| ACTB N-term PBS 13 RT 29 AttB 36 GA atgRNA | GCTATTCTCGCAGCTCACCAgttttagagctagaaatagcaagttaaaataaggctagtccgttatcaacttgaaaaagtggcaccgagtcggtgcGACGAGCGCGGCGATATCATCATCCATGGtgatcctgacgacggagtccgccgtcgtcgacaagcTGAGCTGCGAGAA |
| ACTB N-term PBS 13 RT 29 AttB 31 GA atgRNA | GCTATTCTCGCAGCTCACCAgttttagagctagaaatagcaagttaaaataaggctagtccgttatcaacttgaaaaagtggcaccgagtcggtgcGACGAGCGCGGCGATATCATCATCCATGGatcctgacgacggagtccgccgtcgtcgacaTGAGCTGCGAGAA |
| ACTB N-term PBS 13 RT 29 AttB 26 GA atgRNA | GCTATTCTCGCAGCTCACCAgttttagagctagaaatagcaagttaaaataaggctagtccgttatcaacttgaaaaagtggcaccgagtcggtgcGACGAGCGCGGCGATATCATCATCCATGGcctgacgacggagtccgccgtcgtcgTGAGCTGCGAGAA |
| ACTB N-term PBS 13 RT 29 AttB 21 GA atgRNA | GCTATTCTCGCAGCTCACCAgttttagagctagaaatagcaagttaaaataaggctagtccgttatcaacttgaaaaagtggcaccgagtcggtgcGACGAGCGCGGCGATATCATCATCCATGGtgacgacggagtccgccgtcgTGAGCTGCGAGAA |
| ACTB N-term PBS 13 RT 29 AttB 16 GA atgRNA | GCTATTCTCGCAGCTCACCAgttttagagctagaaatagcaagttaaaataaggctagtccgttatcaacttgaaaaagtggcaccgagtcggtgcGACGAGCGCGGCGATATCATCATCCATGGacgacggagtccgccgTGAGCTGCGAGAA |
| ACTB N-term PBS 13 RT 29 AttB 11 GA atgRNA | GCTATTCTCGCAGCTCACCAgttttagagctagaaatagcaagttaaaataaggctagtccgttatcaacttgaaaaagtggcaccgagtcggtgcGACGAGCGCGGCGATATCATCATCCATGGgacggagtccgTGAGCTGCGAGAA |
| ACTB N-term PBS 13 RT 29 AttB 6 GA atgRNA | GCTATTCTCGCAGCTCACCAgttttagagctagaaatagcaagttaaaataaggctagtccgttatcaacttgaaaaagtggcaccgagtcggtgcGACGAGCGCGGCGATATCATCATCCATGGcggagtTGAGCTGCGAGAA |
| ACTB N-term PBS_18_RT_34_with_Lox71_Cre atgRNA | GAAGCCGGCCTTGCACATGCgttttagagctagaaatagcaagttaaaataaggctagtccgttatcaacttgaaaaagtggcaccgagtcggtgcTCGACGACGAGCGCGGCGATATCATCATCCATGGtaccgttcgtatagcatacattatacgaagttatTGAGCTGCGAGAATAGCC |
| ACTB N-term PBS_18_RT_29_with_Lox71_Cre atgRNA | GAAGCCGGCCTTGCACATGCgttttagagctagaaatagcaagttaaaataaggctagtccgttatcaacttgaaaaagtggcaccgagtcggtgcGACGAGCGCGGCGATATCATCATCCATGGtaccgttcgtatagcatacattatacgaagttatTGAGCTGCGAGAATAGCC |
| ACTB N-term PBS_13_RT_34_with_Lox71_Cre atgRNA | GAAGCCGGCCTTGCACATGCgttttagagctagaaatagcaagttaaaataaggctagtccgttatcaacttgaaaaagtggcaccgagtcggtgcTCGACGACGAGCGCGGCGATATCATCATCCATGGtaccgttcgtatagcatacattatacgaagttatTGAGCTGCGAGAA |
| ACTB N-term PBS_13_RT_16_with_Lox71_Cre atgRNA | GAAGCCGGCCTTGCACATGCgttttagagctagaaatagcaagttaaaataaggctagtccgttatcaacttgaaaaagtggcaccgagtcggtgcATATCATCATCCATGGtaccgttcgtatagcatacattatacgaagttatTGAGCTGCGAGAA |
| ACTB N-term Nicking guide 2 +93 guide | CCCCACGATGGAGGGGAAGAgttttagagctagaaatagcaagttaaaataaggctagtccgttatcaacttgaaaaagtggcaccgagtcggtgc |
| LMNB1 N-term Nicking guide 2 +87 guide | CCTTCTCCTGGAGCCGCGACgttttagagctagaaatagcaagttaaaataaggctagtccgttatcaacttgaaaaagtggcaccgagtcggtgc |
| ACTB N-term PBS 13 RT 29 AttB 46 N191352_143_72 integrase | GCTATTCTCGCAGCTCACCAgttttagagctagaaatagcaagttaaaataaggctagtccgttatcaacttgaaaaagtggcaccgagtcggtgcGACGAGCGCGGCGATATCATCATCCATGGcattatatgttcttacagtatggcggcccggattgtaaaaacatataatgTGAGCTGCGAGAA |
| ACTB N-term PBS 13 RT | GCTATTCTCGCAGCTCACCAgttttagagctagaaatagcaagttaaaataaggctagtccgttatca |

| | |
|---|---|
| 29 AttB 46 N684346_90_69 integrase | acttgaaaaagtggcaccgagtcggtgcGACGAGCGCGGCGATATCATCATCCATGGcgttatagggtattacagtatggcggtcggtactgcaataccctataacgTGAGCTGCGAGAA |
| ACTB N-term PBS 13 RT 29 AttB 46 N675015_95_5 integrase | GCTATTCTCGCAGCTCACCAgttttagagctagaaatagcaagttaaaataaggctagtccgttatcaacttgaaaaagtggcaccgagtcggtgcGACGAGCGCGGCGATATCATCATCCATGGtgtatcattttcatatagttagcacctgcacactatatgaaaatgatacaTGAGCTGCGAGAA |
| ACTB N-term PBS 13 RT 29 AttB 46 N189929_49_54 integrase | GCTATTCTCGCAGCTCACCAgttttagagctagaaatagcaagttaaaataaggctagtccgttatcaacttgaaaaagtggcaccgagtcggtgcGACGAGCGCGGCGATATCATCATCCATGGtgtctactatctgtatatgcgacacatgtggcataaagacatagtagacaTGAGCTGCGAGAA |
| ACTB N-term PBS 13 RT 29 AttB 46 N203911_45186_6 integrase | GCTATTCTCGCAGCTCACCAgttttagagctagaaatagcaagttaaaataaggctagtccgttatcaacttgaaaaagtggcaccgagtcggtgcGACGAGCGCGGCGATATCATCATCCATGGcatcgaccctgacgcatgcggaggcggcgctccatgcgtctgacctcattTGAGCTGCGAGAA |
| ACTB N-term PBS 13 RT 29 AttB 46 N687663_53_29 integrase | GCTATTCTCGCAGCTCACCAgttttagagctagaaatagcaagttaaaataaggctagtccgttatcaacttgaaaaagtggcaccgagtcggtgcGACGAGCGCGGCGATATCATCATCCATGGgttagtacccaaatgacaaaaggtcatccttttatcatttgggtactaacTGAGCTGCGAGAA |
| ACTB N-term PBS 13 RT 29 AttB 46 N687611_90_68 integrase | GCTATTCTCGCAGCTCACCAgttttagagctagaaatagcaagttaaaataaggctagtccgttatcaacttgaaaaagtggcaccgagtcggtgcGACGAGCGCGGCGATATCATCATCCATGGcttattaaaacccgttccgcttctgtcaaagcggcatcggttttataaacTGAGCTGCGAGAA |
| ACTB N-term PBS 13 RT 29 AttB 46 N190156_234_12 integrase | GCTATTCTCGCAGCTCACCAgttttagagctagaaatagcaagttaaaataaggctagtccgttatcaacttgaaaaagtggcaccgagtcggtgcGACGAGCGCGGCGATATCATCATCCATGGggcgtgatggtcgtgaacctcaacatgacgacgaacacgacctcgcggccTGAGCTGCGAGAA |
| ACTB N-term PBS 13 RT 29 AttB 46 N191533_224_76 integrase | GCTATTCTCGCAGCTCACCAgttttagagctagaaatagcaagttaaaataaggctagtccgttatcaacttgaaaaagtggcaccgagtcggtgcGACGAGCGCGGCGATATCATCATCCATGGtctacatcttgaatatatcaagttataactttgaattatatcagtttataTGAGCTGCGAGAA |
| ACTB N-term PBS 13 RT 29 AttB 46 N208621_9_15 integrase | GCTATTCTCGCAGCTCACCAgttttagagctagaaatagcaagttaaaataaggctagtccgttatcaacttgaaaaagtggcaccgagtcggtgcGACGAGCGCGGCGATATCATCATCCATGGaattatatctaaaagcactaagctccgccatactgcttttagatataataTGAGCTGCGAGAA |
| ACTB N-term PBS 13 RT 29 AttB 46 Bacillus_cereus_AH187_38 bp_Att | GCTATTCTCGCAGCTCACCAgttttagagctagaaatagcaagttaaaataaggctagtccgttatcaacttgaaaaagtggcaccgagtcggtgcGACGAGCGCGGCGATATCATCATCCATGGgatatggggaagtgaatcagtacaaccgccacagtaccTGAGCTGCGAGAA |
| ACTB N-term PBS 13 RT 29 AttB 46 Bacillus_cereus_AH187_38 bp_Att_rc | GCTATTCTCGCAGCTCACCAgttttagagctagaaatagcaagttaaaataaggctagtccgttatcaacttgaaaaagtggcaccgagtcggtgcGACGAGCGCGGCGATATCATCATCCATGGggtactgtggcggttgtactgattcacttccccatatcTGAGCTGCGAGAA |
| ACTB N-term PBS 13 RT 29 AttB 46 Staphylococcus_lugdunensis _N920143_38bp_Att | GCTATTCTCGCAGCTCACCAgttttagagctagaaatagcaagttaaaataaggctagtccgttatcaacttgaaaaagtggcaccgagtcggtgcGACGAGCGCGGCGATATCATCATCCATGGtgggtggtacaggtgccacattagttgtaccatttatgTGAGCTGCGAGAA |
| ACTB N-term PBS 13 RT 29 AttB 46 Staphylococcus_lugdunensis _N920143_38bp_Att_rc | GCTATTCTCGCAGCTCACCAgttttagagctagaaatagcaagttaaaataaggctagtccgttatcaacttgaaaaagtggcaccgagtcggtgcGACGAGCGCGGCGATATCATCATCCATGGcataaatggtacaactaatgtggcacctgtaccacccaTGAGCTGCGAGAA |
| ACTB N-term PBS 13 RT 29 AttB 46 Bacillus_cytotoxicus_NVH_ 391-98_38bp_Att | GCTATTCTCGCAGCTCACCAgttttagagctagaaatagcaagttaaaataaggctagtccgttatcaacttgaaaaagtggcaccgagtcggtgcGACGAGCGCGGCGATATCATCATCCATGGgttgtttttccagatccagttggtcctgtaaatataagTGAGCTGCGAGAA |

| | |
|---|---|
| ACTB N-term PBS 13 RT 29 AttB 46 Bacillus_cytotoxicus_NVH_391-98_38bp_Att_rc | GCTATTCTCGCAGCTCACCAgttttagagctagaaatagcaagttaaaataaggctagtccgttatcaacttgaaaaagtggcaccgagtcggtgcGACGAGCGCGGCGATATCATCATCCATGGcttatatttacaggaccaactggatctggaaaaacaacTGAGCTGCGAGAA |
| ACTB N-term PBS 13 RT 29 AttB 46 Bacillus_cereus_AH187_Att 36bp | GCTATTCTCGCAGCTCACCAgttttagagctagaaatagcaagttaaaataaggctagtccgttatcaacttgaaaaagtggcaccgagtcggtgcGACGAGCGCGGCGATATCATCATCCATGGgtactgtggcggttgtactgattcacttccccatatTGAGCTGCGAGAA |
| ACTB N-term PBS 13 RT 29 AttB 46 Bacillus_cereus_AH187_Att 34bp | GCTATTCTCGCAGCTCACCAgttttagagctagaaatagcaagttaaaataaggctagtccgttatcaacttgaaaaagtggcaccgagtcggtgcGACGAGCGCGGCGATATCATCATCCATGGtactgtggcggttgtactgattcacttccccataTGAGCTGCGAGAA |
| ACTB N-term PBS 13 RT 29 AttB 46 Bacillus_cereus_AH187_Att 32bp | GCTATTCTCGCAGCTCACCAgttttagagctagaaatagcaagttaaaataaggctagtccgttatcaacttgaaaaagtggcaccgagtcggtgcGACGAGCGCGGCGATATCATCATCCATGGactgtggcggttgtactgattcacttccccatTGAGCTGCGAGAA |
| ACTB N-term PBS 13 RT 29 AttB 46 Bacillus_cereus_AH187_Att_rc 36bp | GCTATTCTCGCAGCTCACCAgttttagagctagaaatagcaagttaaaataaggctagtccgttatcaacttgaaaaagtggcaccgagtcggtgcGACGAGCGCGGCGATATCATCATCCATGGatatggggaagtgaatcagtacaaccgccacagtacTGAGCTGCGAGAA |
| ACTB N-term PBS 13 RT 29 AttB 46 Bacillus_cereus_AH187_Att_rc 34bp | GCTATTCTCGCAGCTCACCAgttttagagctagaaatagcaagttaaaataaggctagtccgttatcaacttgaaaaagtggcaccgagtcggtgcGACGAGCGCGGCGATATCATCATCCATGGtatggggaagtgaatcagtacaaccgccacagtaTGAGCTGCGAGAA |
| ACTB N-term PBS 13 RT 29 AttB 46 Bacillus_cereus_AH187_Att_rc 32bp | GCTATTCTCGCAGCTCACCAgttttagagctagaaatagcaagttaaaataaggctagtccgttatcaacttgaaaaagtggcaccgagtcggtgcGACGAGCGCGGCGATATCATCATCCATGGatggggaagtgaatcagtacaaccgccacagtTGAGCTGCGAGAA |
| ACTB N-term PBS 13 RT 29 AttB 46 Staphylococcus_lugdunensis_N920143_Att 36bp | GCTATTCTCGCAGCTCACCAgttttagagctagaaatagcaagttaaaataaggctagtccgttatcaacttgaaaaagtggcaccgagtcggtgcGACGAGCGCGGCGATATCATCATCCATGGataaatggtacaactaatgtggcacctgtaccacccTGAGCTGCGAGAA |
| ACTB N-term PBS 13 RT 29 AttB 46 Staphylococcus_lugdunensis_N920143_Att 34bp | GCTATTCTCGCAGCTCACCAgttttagagctagaaatagcaagttaaaataaggctagtccgttatcaacttgaaaaagtggcaccgagtcggtgcGACGAGCGCGGCGATATCATCATCCATGGtaaatggtacaactaatgtggcacctgtaccaccTGAGCTGCGAGAA |
| ACTB N-term PBS 13 RT 29 AttB 46 Staphylococcus_lugdunensis_N920143_Att 32bp | GCTATTCTCGCAGCTCACCAgttttagagctagaaatagcaagttaaaataaggctagtccgttatcaacttgaaaaagtggcaccgagtcggtgcGACGAGCGCGGCGATATCATCATCCATGGaaatggtacaactaatgtggcacctgtaccacTGAGCTGCGAGAA |
| ACTB N-term PBS 13 RT 29 AttB 46 Staphylococcus_lugdunensis_N920143_Att_rc 36bp | GCTATTCTCGCAGCTCACCAgttttagagctagaaatagcaagttaaaataaggctagtccgttatcaacttgaaaaagtggcaccgagtcggtgcGACGAGCGCGGCGATATCATCATCCATGGgggtggtacaggtgccacattagttgtaccatttatTGAGCTGCGAGAA |
| ACTB N-term PBS 13 RT 29 AttB 46 Staphylococcus_lugdunensis_N920143_Att_rc 34bp | GCTATTCTCGCAGCTCACCAgttttagagctagaaatagcaagttaaaataaggctagtccgttatcaacttgaaaaagtggcaccgagtcggtgcGACGAGCGCGGCGATATCATCATCCATGGggtggtacaggtgccacattagttgtaccatttaTGAGCTGCGAGAA |

| | |
|---|---|
| ACTB N-term PBS 13 RT 29 AttB 46 Staphylococcus_lugdunensis_N920143_Att_rc 32bp | GCTATTCTCGCAGCTCACCAgttttagagctagaaatagcaagttaaaataaggctagtccgttatcaacttgaaaaagtggcaccgagtcggtgcGACGAGCGCGGCGATATCATCATCCATGGgtggtacaggtgccacattagttgtaccatttTGAGCTGCGAGAA |
| ACTB N-term PBS 13 RT 29 AttB 46 Bacillus_cytotoxicus_NVH_391-98_Att 36bp | GCTATTCTCGCAGCTCACCAgttttagagctagaaatagcaagttaaaataaggctagtccgttatcaacttgaaaaagtggcaccgagtcggtgcGACGAGCGCGGCGATATCATCATCCATGGttatatttacaggaccaactggatctggaaaaacaaTGAGCTGCGAGAA |
| ACTB N-term PBS 13 RT 29 AttB 46 Bacillus_cytotoxicus_NVH_391-98_Att 34bp | GCTATTCTCGCAGCTCACCAgttttagagctagaaatagcaagttaaaataaggctagtccgttatcaacttgaaaaagtggcaccgagtcggtgcGACGAGCGCGGCGATATCATCATCCATGGtatatttacaggaccaactggatctggaaaaacaTGAGCTGCGAGAA |
| ACTB N-term PBS 13 RT 29 AttB 46 Bacillus_cytotoxicus_NVH_391-98_Att 32bp | GCTATTCTCGCAGCTCACCAgttttagagctagaaatagcaagttaaaataaggctagtccgttatcaacttgaaaaagtggcaccgagtcggtgcGACGAGCGCGGCGATATCATCATCCATGGatatttacaggaccaactggatctggaaaaacTGAGCTGCGAGAA |
| ACTB N-term PBS 13 RT 29 AttB 46 Bacillus_cytotoxicus_NVH_391-98_Att_rc 36bp | GCTATTCTCGCAGCTCACCAgttttagagctagaaatagcaagttaaaataaggctagtccgttatcaacttgaaaaagtggcaccgagtcggtgcGACGAGCGCGGCGATATCATCATCCATGGttgttttccagatccagttggtcctgtaaatataaTGAGCTGCGAGAA |
| ACTB N-term PBS 13 RT 29 AttB 46 Bacillus_cytotoxicus_NVH_391-98_Att_rc 34bp | GCTATTCTCGCAGCTCACCAgttttagagctagaaatagcaagttaaaataaggctagtccgttatcaacttgaaaaagtggcaccgagtcggtgcGACGAGCGCGGCGATATCATCATCCATGGtgtttttccagatccagttggtcctgtaaatataTGAGCTGCGAGAA |
| ACTB N-term PBS 13 RT 29 AttB 46 Bacillus_cytotoxicus_NVH_391-98_Att_rc 32bp | GCTATTCTCGCAGCTCACCAgttttagagctagaaatagcaagttaaaataaggctagtccgttatcaacttgaaaaagtggcaccgagtcggtgcGACGAGCGCGGCGATATCATCATCCATGGgttttccagatccagttggtcctgtaaatatTGAGCTGCGAGAA |
| Bacillus_cereus_AH187_Att_rc_36 LMNB1 PBS 9 RT 10 AttB 36 atgRNA | GCTGTCTCCGCCGCCCGCCAgttttagagctagaaatagcaagttaaaataaggctagtccgttatcaacttgaaaaagtggcaccgagtcggtgcAGTCGCCATGatatggggaagtgaatcagtacaaccgccacagtacCGGGCGGCG |
| Bacillus_cereus_AH187_Att_rc_36 NOLC1 PBS 18 RT 29 AttB 36 atgRNA | GCGTATTGCCTGGAGGATGGgttttagagctagaaatagcaagttaaaataaggctagtccgttatcaacttgaaaaagtggcaccgagtcggtgcGAACCACGCGGCGAATGCCGGCGTCCGCCatatggggaagtgaatcagtacaaccgccacagtacTCCTCCAGGCAATACGCG |
| Bacillus_cereus_AH187_Att_rc_36 SUPT16H PBS 13 RT 24 AttB 36 atgRNA | GAGAAGCGGCGTCCGGGGCTAgttttagagctagaaatagcaagttaaaataaggctagtccgttatcaacttgaaaaagtggcaccgagtcggtgcTCTTTGTCCAGAGTCACAGCCATAatatggggaagtgaatcagtacaaccgccacagtacCCCCGGACGCCGC |
| Bacillus_cereus_AH187_Att_rc_36 SRRM2 PBS 13 RT 24 AttB 36 atgRNA | GGGCACGGGGCCATGTACAAgttttagagctagaaatagcaagttaaaataaggctagtccgttatcaacttgaaaaagtggcaccgagtcggtgcGGCGTCGGCAGCCCGATCCCGTTGatatggggaagtgaatcagtacaaccgccacagtacTACATGGCCCCGT |
| Bacillus_cereus_AH187_Att_rc_36 DEPDC4 PBS 18 RT 24 AttB 36 atgRNA | GTGTCAGGTGGGGCGGGGCTAgttttagagctagaaatagcaagttaaaataaggctagtccgttatcaacttgaaaaagtggcaccgagtcggtgcGCTGGCTCCTCCCCTGGCACCATAatatggggaagtgaatcagtacaaccgccacagtacCCCCGCCCCACCTGACAC |
| Bacillus_cereus_AH187_Att_rc_36 NES PBS 13 RT 28 AttB 36 atgRNA | GAGTGGGTCAGACGAGCAGGAgttttagagctagaaatagcaagttaaaataaggctagtccgttatcaacttgaaaaagtggcaccgagtcggtgcCGACTCCTCCCCCATGCAGCCCTCCATCatatggggaagtgaatcagtacaaccgccacagtacTGCTCGTCTGACC |
| B. cereus LMNB1_PBS 9 | GCTGTCTCCGCCGCCCGCCAgttttagagctagaaatagcaagttaaaataaggctagtccgttatca |

| | |
|---|---|
| RT 20 AttB 36 atgRNA | acttgaaaaagtggcaccgagtcggtgcCGGGGGTCGCAGTCGCCATGatatggggaagtgaatcagtacaaccgccacagtacCGGGCGGCG |
| B. cereus LMNB1_PBS 13 RT 20 AttB 36 atgRNA | GGGCACGGGGCCATGTACAAgttttagagctagaaatagcaagttaaaataaggctagtccgttatcaacttgaaaaagtggcaccgagtcggtgcCGGGGGTCGCAGTCGCCATGatatggggaagtgaatcagtacaaccgccacagtacCGGGCGGCGGAGA |
| B. cereus LMNB1_PBS 13 RT 29 AttB 36 atgRNA | GGGCACGGGGCCATGTACAAgttttagagctagaaatagcaagttaaaataaggctagtccgttatcaacttgaaaaagtggcaccgagtcggtgcGCGGCGGCACGGGGGTCGCAGTCGCCATGatatggggaagtgaatcagtacaaccgccacagtacCGGGCGGCGGAGA |
| B. cereus NOLC1_PBS 13 RT 29 AttB 36 atgRNA | GCGTATTGCCTGGAGGATGGgttttagagctagaaatagcaagttaaaataaggctagtccgttatcaacttgaaaaagtggcaccgagtcggtgcGAACCACGCGGCGAATGCCGGCGTCCGCCatatggggaagtgaatcagtacaaccgccacagtacTCCTCCAGGCAAT |
| B. cereus NOLC1_PBS 13 RT 20 AttB 36 atgRNA | GGGCACGGGGCCATGTACAAgttttagagctagaaatagcaagttaaaataaggctagtccgttatcaacttgaaaaagtggcaccgagtcggtgcGGCGAATGCCGGCGTCCGCCatatggggaagtgaatcagtacaaccgccacagtacTCCTCCAGGCAAT |
| B. cereus NOLC1_PBS 18 RT 20 AttB 36 atgRNA | GGGCACGGGGCCATGTACAAgttttagagctagaaatagcaagttaaaataaggctagtccgttatcaacttgaaaaagtggcaccgagtcggtgcGGCGAATGCCGGCGTCCGCCatatggggaagtgaatcagtacaaccgccacagtacTCCTCCAGGCAATACGCG |
| B. cereus SRRM2_PBS 9 RT 24 AttB 36 atgRNA | GGGCACGGGGCCATGTACAAgttttagagctagaaatagcaagttaaaataaggctagtccgttatcaacttgaaaaagtggcaccgagtcggtgcGGCGTCGGCAGCCCGATCCCGTTGatatggggaagtgaatcagtacaaccgccacagtacTACATGGCC |
| B. cereus SRRM2_PBS 9 RT 10 AttB 36 atgRNA | GGGCACGGGGCCATGTACAAgttttagagctagaaatagcaagttaaaataaggctagtccgttatcaacttgaaaaagtggcaccgagtcggtgcGATCCCGTTGatatggggaagtgaatcagtacaaccgccacagtacTACATGGCC |
| B. cereus SRRM2_PBS 13 RT 10 AttB 36 atgRNA | GGGCACGGGGCCATGTACAAgttttagagctagaaatagcaagttaaaataaggctagtccgttatcaacttgaaaaagtggcaccgagtcggtgcGATCCCGTTGatatggggaagtgaatcagtacaaccgccacagtacTACATGGCCCCGT |
| Screen validation guides ACTB_1_11_24_38 | GCTATTCTCGCAGCTCACCAgttttagagctagaaatagcaagttaaaataaggctagtccgttatcaacttgaaaaagtggcaccgagtcggtgcgcgcggcgatatcatcatccatggatgatcctgacgacggagaccgccgtcgtcgacaagcctgagctgcgag |
| Screen validation guides ACTB_1_16_18_43 | GCTATTCTCGCAGCTCACCAgttttagagctagaaatagcaagttaaaataaggctagtccgttatcaacttgaaaaagtggcaccgagtcggtgccgatatcatcatccatggcggatgatcctgacgacggagaccgccgtcgtcgacaagccggctgagctgcgagaatag |
| Screen validation guides LMNB1_1_8_26_38 | GCTGTCTCCGCCGCCCGCCAgttttagagctagaaatagcaagttaaaataaggctagtccgttatcaacttgaaaaagtggcaccgagtcggtgcgcggcacgggggtcgcagtcgccatgatgatcctgacgacggagaccgccgtcgtcgacaagcccgggcggc |
| Screen validation guides NOLC1_1_15_16_43 | GCGTATTGCCTGGAGGATGGgttttagagctagaaatagcaagttaaaataaggctagtccgttatcaacttgaaaaagtggcaccgagtcggtgcaatgccggcgtccgcccggatgatcctgacgacggagaccgccgtcgtcgacaagccggctcctccaggcaatac |
| Screen validation guides NOLC1_1_14_10_38 | GCGTATTGCCTGGAGGATGGgttttagagctagaaatagcaagttaaaataaggctagtccgttatcaacttgaaaaagtggcaccgagtcggtgcggcgtccgccatgatcctgacgacggagaccgccgtcgtcgacaagcctcctccaggcaata |
| Screen validation guides SERPIN_13_32_38 | GGGAAATGCATCTTGCACAAgttttagagctagaaatagcaagttaaaataaggctagtccgttatcaacttgaaaaagtggcaccgagtcggtgcagcccctccatgctctctagctgttgccattgggcttgtcgacgacggcggtctccgtcgtcaggatcattgcaagatgcatt |
| Screen validation guides DEPDC4_8_10_38 | GTGTCAGGTGGGGCGGGGCTAgttttagagctagaaatagcaagttaaaataaggctagtccgttatcaacttgaaaaagtggcaccgagtcggtgctggcaccataatgatcctgacgacggagaccgccgtcgtcgacaag |

| | |
|---|---|
| | ccccccgccc |
| SERPIN Nicking guide -107 guide | GTGGGGACAGCCCCGTCTCTgttttagagctagaaatagcaagttaaaataaggctagtccgttatcaacttgaaaaagtggcaccgagtcggtgc |
| SERPIN Nicking guide -91 guide | GCTCTTGGGAAAAAAACCCTAgttttagagctagaaatagcaagttaaaataaggctagtccgttatcaacttgaaaaagtggcaccgagtcggtgc |
| SERPIN Nicking guide -90 guide | GTCTTGGGAAAAAAACCCTAAgttttagagctagaaatagcaagttaaaataaggctagtccgttatcaacttgaaaaagtggcaccgagtcggtgc |
| SERPIN Nicking guide -84 guide | GAAAAAAACCCTAAGGGCTGgttttagagctagaaatagcaagttaaaataaggctagtccgttatcaacttgaaaaagtggcaccgagtcggtgc |
| SERPIN Nicking guide -67 guide | GCTGAGGATCCTTGTGAGTGTgttttagagctagaaatagcaagttaaaataaggctagtccgttatcaacttgaaaaagtggcaccgagtcggtgc |
| SERPIN Nicking guide -66 guide | GTGAGGATCCTTGTGAGTGTTgttttagagctagaaatagcaagttaaaataaggctagtccgttatcaacttgaaaaagtggcaccgagtcggtgc |
| SERPIN Nicking guide -63 guide | GGATCCTTGTGAGTGTTGGGgttttagagctagaaatagcaagttaaaataaggctagtccgttatcaacttgaaaaagtggcaccgagtcggtgc |
| SERPIN Nicking guide -62 guide | GATCCTTGTGAGTGTTGGGTgttttagagctagaaatagcaagttaaaataaggctagtccgttatcaacttgaaaaagtggcaccgagtcggtgc |
| SERPIN Nicking guide -49 guide | GTTGGGTGGGAACAGCTCCCgttttagagctagaaatagcaagttaaaataaggctagtccgttatcaacttgaaaaagtggcaccgagtcggtgc |
| SERPIN Nicking guide -46 guide | GGGTGGGAACAGCTCCCAGGgttttagagctagaaatagcaagttaaaataaggctagtccgttatcaacttgaaaaagtggcaccgagtcggtgc |
| SERPIN Nicking guide +34 guide | GCTTCTGTGCAGCAGTTTCCCgttttagagctagaaatagcaagttaaaataaggctagtccgttatcaacttgaaaaagtggcaccgagtcggtgc |
| SERPIN Nicking guide +48 guide | GTTTCCCTGGCCACTAAATAGgttttagagctagaaatagcaagttaaaataaggctagtccgttatcaacttgaaaaagtggcaccgagtcggtgc |
| SERPIN Nicking guide +49 guide | GTTCCCTGGCCACTAAATAGTgttttagagctagaaatagcaagttaaaataaggctagtccgttatcaacttgaaaaagtggcaccgagtcggtgc |
| SERPIN Nicking guide +71 guide | GATTAGATAGAAGCCCTCCAgttttagagctagaaatagcaagttaaaataaggctagtccgttatcaacttgaaaaagtggcaccgagtcggtgc |
| SERPIN Nicking guide +72 guide | GATTAGATAGAAGCCCTCCAAgttttagagctagaaatagcaagttaaaataaggctagtccgttatcaacttgaaaaagtggcaccgagtcggtgc |

**Supplementary Table 5.** Sequences of insertion sites

| **Description** | **Forward Sequence (5'-3')** | **Reverse Sequence (5'-3')** |
|---|---|---|
| Bxb1_AttP_GT_original _site | GTGGTTTGTCTGGTCAACCACCGCGGTCTCA GTGGTGTACGGTACAAACCCA | TGGGTTTGTACCGTACACCACTGAGACCGCGGT GGTTGACCAGACAAACCAC |
| Bxb1_AttP_CG_site | GTGGTTTGTCTGGTCAACCACCGCGcgCTCAG TGGTGTACGGTACAAACCCA | TGGGTTTGTACCGTACACCACTGAGCGCGCGGT GGTTGACCAGACAAACCAC |
| Bxb1_AttP_GC_site | GTGGTTTGTCTGGTCAACCACCGCGgcCTCAG TGGTGTACGGTACAAACCCA | TGGGTTTGTACCGTACACCACTGAGGCCGCGGT GGTTGACCAGACAAACCAC |
| Bxb1_AttP_AT_site | GTGGTTTGTCTGGTCAACCACCGCGatCTCAG TGGTGTACGGTACAAACCCA | TGGGTTTGTACCGTACACCACTGAGATCGCGGTG GTTGACCAGACAAACCAC |
| Bxb1_AttP_TA_site | GTGGTTTGTCTGGTCAACCACCGCGtaCTCAG TGGTGTACGGTACAAACCCA | TGGGTTTGTACCGTACACCACTGAGTACGCGGTG GTTGACCAGACAAACCAC |
| Bxb1_AttP_GG_site | GTGGTTTGTCTGGTCAACCACCGCGggCTCAG TGGTGTACGGTACAAACCCA | TGGGTTTGTACCGTACACCACTGAGCCCGCGGTG GTTGACCAGACAAACCAC |
| Bxb1_AttP_TT_site | GTGGTTTGTCTGGTCAACCACCGCGttCTCAGT GGTGTACGGTACAAACCCA | TGGGTTTGTACCGTACACCACTGAGAACGCGGT GGTTGACCAGACAAACCAC |
| Bxb1_AttP_GA_site | GTGGTTTGTCTGGTCAACCACCGCGgaCTCAG TGGTGTACGGTACAAACCCA | TGGGTTTGTACCGTACACCACTGAGTCCGCGGTG GTTGACCAGACAAACCAC |
| Bxb1_AttP_AG_site | GTGGTTTGTCTGGTCAACCACCGCGagCTCAG TGGTGTACGGTACAAACCCA | TGGGTTTGTACCGTACACCACTGAGCTCGCGGTG GTTGACCAGACAAACCAC |
| Bxb1_AttP_CC_site | GTGGTTTGTCTGGTCAACCACCGCGccCTCAG TGGTGTACGGTACAAACCCA | TGGGTTTGTACCGTACACCACTGAGGGCGCGGT GGTTGACCAGACAAACCAC |
| Bxb1_AttP_TC_site | GTGGTTTGTCTGGTCAACCACCGCGtcCTCAG TGGTGTACGGTACAAACCCA | TGGGTTTGTACCGTACACCACTGAGGACGCGGT GGTTGACCAGACAAACCAC |
| Bxb1_AttP_CT_site | GTGGTTTGTCTGGTCAACCACCGCGctCTCAG TGGTGTACGGTACAAACCCA | TGGGTTTGTACCGTACACCACTGAGAGCGCGGT GGTTGACCAGACAAACCAC |
| Bxb1_AttP_AA_site | GTGGTTTGTCTGGTCAACCACCGCGaaCTCAG TGGTGTACGGTACAAACCCA | TGGGTTTGTACCGTACACCACTGAGTTCGCGGTG GTTGACCAGACAAACCAC |
| Bxb1_AttP_CA_site | GTGGTTTGTCTGGTCAACCACCGCGcaCTCAG TGGTGTACGGTACAAACCCA | TGGGTTTGTACCGTACACCACTGAGTGCGCGGTG GTTGACCAGACAAACCAC |
| Bxb1_AttP_AC_site | GTGGTTTGTCTGGTCAACCACCGCGacCTCAG TGGTGTACGGTACAAACCCA | TGGGTTTGTACCGTACACCACTGAGGTCGCGGTG GTTGACCAGACAAACCAC |
| Bxb1_AttP_TG_site | GTGGTTTGTCTGGTCAACCACCGCGtgCTCAG TGGTGTACGGTACAAACCCA | TGGGTTTGTACCGTACACCACTGAGCACGCGGT GGTTGACCAGACAAACCAC |
| Bxb1_AttB_46_GT_ original_site | GGCCGGCTTGTCGACGACGGCGGTCTCCGTC GTCAGGATCATCCGG | CCGGATGATCCTGACGACGGAGACCGCCGTCGT CGACAAGCCGGCC |
| Bxb1_AttB_46_AA_site | GGCCGGCTTGTCGACGACGGCGaaCTCCGTCG TCAGGATCATCCGG | CCGGATGATCCTGACGACGGAGTTCGCCGTCGT CGACAAGCCGGCC |
| Bxb1_AttB_46_GA_site | GGCCGGCTTGTCGACGACGGCGgaCTCCGTCG TCAGGATCATCCGG | CCGGATGATCCTGACGACGGAGTCCGCCGTCGT CGACAAGCCGGCC |
| Bxb1_AttB_46_CA_site | GGCCGGCTTGTCGACGACGGCGcaCTCCGTCG | CCGGATGATCCTGACGACGGAGTGCGCCGTCGT |

| | | |
|---|---|---|
| | TCAGGATCATCCGG | CGACAAGCCGGCC |
| Bxb1_AttB_46_TA_site | GGCCGGCTTGTCGACGACGGCGtaCTCCGTCG TCAGGATCATCCGG | CCGGATGATCCTGACGACGGAGTACGCCGTCGT CGACAAGCCGGCC |
| Bxb1_AttB_46_AG_site | GGCCGGCTTGTCGACGACGGCGagCTCCGTCG TCAGGATCATCCGG | CCGGATGATCCTGACGACGGAGCTCGCCGTCGT CGACAAGCCGGCC |
| Bxb1_AttB_46_GG_site | GGCCGGCTTGTCGACGACGGCGggCTCCGTCG TCAGGATCATCCGG | CCGGATGATCCTGACGACGGAGCCCGCCGTCGT CGACAAGCCGGCC |
| Bxb1_AttB_46_CG_site | GGCCGGCTTGTCGACGACGGCGcgCTCCGTCG TCAGGATCATCCGG | CCGGATGATCCTGACGACGGAGCGCGCCGTCGT CGACAAGCCGGCC |
| Bxb1_AttB_46_TG_site | GGCCGGCTTGTCGACGACGGCGtgCTCCGTCG TCAGGATCATCCGG | CCGGATGATCCTGACGACGGAGCACGCCGTCGT CGACAAGCCGGCC |
| Bxb1_AttB_46_AC_site | GGCCGGCTTGTCGACGACGGCGacCTCCGTCG TCAGGATCATCCGG | CCGGATGATCCTGACGACGGAGGTCGCCGTCGT CGACAAGCCGGCC |
| Bxb1_AttB_46_GC_site | GGCCGGCTTGTCGACGACGGCGgcCTCCGTCG TCAGGATCATCCGG | CCGGATGATCCTGACGACGGAGGCCGCCGTCGT CGACAAGCCGGCC |
| Bxb1_AttB_46_CC_site | GGCCGGCTTGTCGACGACGGCGccCTCCGTCG TCAGGATCATCCGG | CCGGATGATCCTGACGACGGAGGGCGCCGTCGT CGACAAGCCGGCC |
| Bxb1_AttB_46_TC_site | GGCCGGCTTGTCGACGACGGCGtcCTCCGTCG TCAGGATCATCCGG | CCGGATGATCCTGACGACGGAGGACGCCGTCGT CGACAAGCCGGCC |
| Bxb1_AttB_46_AT_site | GGCCGGCTTGTCGACGACGGCGatCTCCGTCG TCAGGATCATCCGG | CCGGATGATCCTGACGACGGAGATCGCCGTCGT CGACAAGCCGGCC |
| Bxb1_AttB_46_CT_site | GGCCGGCTTGTCGACGACGGCGctCTCCGTCG TCAGGATCATCCGG | CCGGATGATCCTGACGACGGAGAGCGCCGTCGT CGACAAGCCGGCC |
| Bxb1_AttB_46_TT_site | GGCCGGCTTGTCGACGACGGCGttCTCCGTCG TCAGGATCATCCGG | CCGGATGATCCTGACGACGGAGAACGCCGTCGT CGACAAGCCGGCC |
| Bxb1_AttB_38_GT_site | GGCTTGTCGACGACGGCGGTCTCCGTCGTCA GGATCAT | ATGATCCTGACGACGGAGACCGCCGTCGTCGAC AAGCC |
| Bxb1_AttB_38_AA_site | GGCTTGTCGACGACGGCGaaCTCCGTCGTCAG GATCAT | ATGATCCTGACGACGGAGTTCGCCGTCGTCGAC AAGCC |
| Bxb1_AttB_38_GA_site | GGCTTGTCGACGACGGCGgaCTCCGTCGTCAG GATCAT | ATGATCCTGACGACGGAGTCCGCCGTCGTCGAC AAGCC |
| Bxb1_AttB_38_CA_site | GGCTTGTCGACGACGGCGcaCTCCGTCGTCAG GATCAT | ATGATCCTGACGACGGAGTGCGCCGTCGTCGAC AAGCC |
| Bxb1_AttB_38_TA_site | GGCTTGTCGACGACGGCGtaCTCCGTCGTCAG GATCAT | ATGATCCTGACGACGGAGTACGCCGTCGTCGAC AAGCC |
| Bxb1_AttB_38_AG_site | GGCTTGTCGACGACGGCGagCTCCGTCGTCAG GATCAT | ATGATCCTGACGACGGAGCTCGCCGTCGTCGAC AAGCC |
| Bxb1_AttB_38_GG_site | GGCTTGTCGACGACGGCGggCTCCGTCGTCAG GATCAT | ATGATCCTGACGACGGAGCCCGCCGTCGTCGAC AAGCC |
| Bxb1_AttB_38_CG_site | GGCTTGTCGACGACGGCGcgCTCCGTCGTCAG GATCAT | ATGATCCTGACGACGGAGCGCGCCGTCGTCGAC AAGCC |
| Bxb1_AttB_38_TG_site | GGCTTGTCGACGACGGCGtgCTCCGTCGTCAG GATCAT | ATGATCCTGACGACGGAGCACGCCGTCGTCGAC AAGCC |

| | | |
|---|---|---|
| Bxb1_AttB_38_AC_site | GGCTTGTCGACGACGGCGacCTCCGTCGTCAG GATCAT | ATGATCCTGACGACGGAGGTCGCCGTCGTCGAC AAGCC |
| Bxb1_AttB_38_GC_site | GGCTTGTCGACGACGGCGgcCTCCGTCGTCAG GATCAT | ATGATCCTGACGACGGAGGCCGCCGTCGTCGAC AAGCC |
| Bxb1_AttB_38_CC_site | GGCTTGTCGACGACGGCGccCTCCGTCGTCAG GATCAT | ATGATCCTGACGACGGAGGGCGCCGTCGTCGAC AAGCC |
| Bxb1_AttB_38_TC_site | GGCTTGTCGACGACGGCGtcCTCCGTCGTCAG GATCAT | ATGATCCTGACGACGGAGGACGCCGTCGTCGAC AAGCC |
| Bxb1_AttB_38_AT_site | GGCTTGTCGACGACGGCGatCTCCGTCGTCAG GATCAT | ATGATCCTGACGACGGAGATCGCCGTCGTCGAC AAGCC |
| Bxb1_AttB_38_CT_site | GGCTTGTCGACGACGGCGctCTCCGTCGTCAG GATCAT | ATGATCCTGACGACGGAGAGCGCCGTCGTCGAC AAGCC |
| Bxb1_AttB_38_TT_site | GGCTTGTCGACGACGGCGttCTCCGTCGTCAG GATCAT | ATGATCCTGACGACGGAGAACGCCGTCGTCGAC AAGCC |
| Cre Lox 66 site | TACCGTTCGTATAATGTATGCTATACGA AGTTAT | ATAACTTCGTATAGCATACATTATACGAAC GGTA |
| Cre Lox 71 site | ATAACTTCGTATAATGTATGCTATACGA ACGGTA | TACCGTTCGTATAGCATACATTATACGAAG TTAT |
| TP901-1 minimal AttB site | TTTACCTTGATTGAGATGTTAATTGTG | CACAATTAACATCTCAATCAAGGTAAA |
| TP901-1 minimal AttP site | GCGAGTTTTTATTTCGTTTATTTCAATTAAGG TAACTAAAAAACTCCTTT | AAAGGAGTTTTTTAGTTACCTTAATTGAAATAAA CGAAATAAAAACTCGC |
| PhiBT1 minimal AttB site | CTGGATCATCTGGATCACTTTCGTCAAAAAC CTG | CAGGTTTTTGACGAAAGTGATCCAGATGATCCA G |
| PhiBT1 minimal AttP site | TTCGGGTGCTGGGTTGTTGTCTCTGGACAGTG ATCCATGGGAAACTACTCAGCACCA | TGGTGCTGAGTAGTTTCCCATGGATCACTGTCCA GAGACAACAACCCAGCACCCGAA |
| Bacillus_cereus_AH187_Int30_38bp_Att | gatatggggaagtgaatcagtacaaccgccacagtacc | ggtactgtggcggttgtactgattcacttccccatatc |
| Staphylococcus_lugdunensis_N920143_Int12_38bp_Att | tgggtggtacaggtgccacattagttgtaccatttatg | cataaatggtacaactaatgtggcacctgtaccaccca |
| Bacillus_cytotoxicus_NVH_391-98_Int13_38bp_Att | gttgtttttccagatccagttggtcctgtaaatataag | cttatatttacaggaccaactggatctggaaaaacaac |
| Bacillus_cereus_AH187_Int30_Att_30 | tggggaagtgaatcagtacaaccgccacag | ctgtggcggttgtactgattcacttcccca |
| Bacillus_cereus_AH187_Int30_Att_28 | ggggaagtgaatcagtacaaccgccaca | tgtggcggttgtactgattcacttcccc |
| Bacillus_cereus_AH187_Int30_Att_26 | gggaagtgaatcagtacaaccgccac | gtggcggttgtactgattcacttccc |
| Bacillus_cereus_AH187_Int30_Att_rc_30 | ctgtggcggttgtactgattcacttcccca | tggggaagtgaatcagtacaaccgccacag |
| Bacillus_cereus_AH1 | tgtggcggttgtactgattcacttcccc | ggggaagtgaatcagtacaaccgccaca |

| | | |
|---|---|---|
| 87_Int30_Att_rc_28 | | |
| Bacillus_cereus_AH187_Int30_Att_rc_26 | gtggcggttgtactgattcacttccc | gggaagtgaatcagtacaaccgccac |
| Bacillus_cytotoxicus_NVH_391-98_Int13_Att_30 | tttttccagatccagttggtcctgtaaata | tatttacaggaccaactggatctggaaaaa |
| Bacillus_cytotoxicus_NVH_391-98_Int13_Att_28 | ttttccagatccagttggtcctgtaaat | atttacaggaccaactggatctggaaaa |
| Bacillus_cytotoxicus_NVH_391-98_Int13_Att_26 | tttccagatccagttggtcctgtaaa | tttacaggaccaactggatctggaaa |
| Bacillus_cytotoxicus_NVH_391-98_Int13_Att_rc_30 | tatttacaggaccaactggatctggaaaaa | tttttccagatccagttggtcctgtaaata |
| Bacillus_cytotoxicus_NVH_391-98_Int13_Att_rc_28 | atttacaggaccaactggatctggaaaa | ttttccagatccagttggtcctgtaaat |
| Bacillus_cytotoxicus_NVH_391-98_Int13_Att_rc_26 | tttacaggaccaactggatctggaaa | tttccagatccagttggtcctgtaaa |
| N680429_560_31_50bp | CATTATATGTTTTTACAATCCGGGCCGCCATACTGTAAGAACATATAATG | cattatatgttcttacagtatggcggcccggattgtaaaaacatataatg |
| N191607_8_101_50bp | CGTTATAGGGTATTGCAGTACCGACCGCCATACTGTAATACCCTATAACG | cgttatagggtattacagtatggcggtcggtactgcaataccctataacg |
| N674992_1_1308_50bp | TGTATCATTTTCATATAGTGTGCAGGTGCTAACTATATGAAAATGATACA | tgtatcattttcatatagttagcacctgcacactatatgaaaatgataca |
| N684613_54_96_50bp | TGTCTACTATGTCTTTATGCCACATGTGTCGCATATACAGATAGTAGACA | tgtctactatctgtatatgcgacacatgtggcataaagacatagtagaca |
| N252616_121_74_50bp | AATGAGGTCAGACGCATGGAGCGCCGCCTCCGCATGCGTCAGGGTCGATG | catcgaccctgacgcatgcggaggcggcgctccatgcgtctgacctcatt |
| N683040_222_19_50bp | GTTAGTACCCAAATGATAAAAGGATGACCTTTTGTCATTTGGGTACTAAC | gttagtacccaaatgacaaaaggtcatccttttatcatttgggtactaac |
| N687537_173_59_50bp | GTTTATAAAACCGATGCCGCTTTGACAGAAGCGGAACGGGTTTTAATAAG | cttattaaaacccgttccgcttctgtcaaagcggcatcggttttataaac |
| N183629_47_40_50bp | GGCCGCGAGGTCGTGTTCGTCGTCATGTTGAGGTTCACGACCATCACGCC | ggcgtgatggtcgtgaacctcaacatgacgacgaacacgacctcgcggcc |
| N191533_224_76_50bp | TATAAACTGATATAATTCAAAGTTATAACTTGATATATTCAAGATGTAGA | tctacatcttgaatatatcaagttataactttgaattatatcagtttata |
| N682356_188_20_50bp | TATTATATCTAAAAGCAGTATGGCGGAGCTTAGTGCTTTTAGATATAATT | aattatatctaaaagcactaagctccgccatactgcttttagatataata |

**Supplementary Table 6.** Mammalian expression plasmids

| Name | Full Description | Benchling link |
|---|---|---|
| PE2-Bxb1 Single Vector | pCMV-PE2-P2A-Bxb1 | https://benchling.com/s/seq-NIW2rI9OUqYpAmedqicN |
| PE2 prime editor | pCMV-PE2 / Addgene #132775 | https://benchling.com/s/seq-C6YkN30h9vvBPv8XKj47 |
| PASTEv3 | pCMV-SpCas9-XTEN3-RT(L139P)-(GGS)6-BxbINT | https://benchling.com/s/seq-6PrptF4jyyE6fkDLKc88 |
| PASTEv4 | pCMV-SpCas9-XTEN-RT(L139P)-(GGGS)3-BceINT | https://benchling.com/s/seq-mGirm9x1J7IS0fKRHhXE |
| ACTB atgRNA with v1 scaffold | ACTB N-term PBS 13 RT 29 AttB 46 atgRNA | https://benchling.com/s/seq-h2yh6lfsvGpgG7XRRLbT |
| ACTB atgRNA with v2 scaffold | ACTB N-term PBS 13 RT 29 AttB 46 atgRNA with v2 scaffold | https://benchling.com/s/seq-qsyXbHzeyxBMzPJEciMg |
| ACTB Nicking +48 | ACTB N-term Nicking guide 1 +48 guide | https://benchling.com/s/seq-VWT8CHMkrIRRnsqoIRUS |
| Bxb1 integrase | pCAG-NLS-HA-Bxb1integrase / Addgene #51271 | https://benchling.com/s/seq-bLuI9aGZCKRbquNPXpjy |
| TP901-1 Integrase | TP901-1 Integrase | https://benchling.com/s/seq-TV8nlYedd6KaXcyMYQ3O |
| PhiBT Integrase | PhiBT Integrase | https://benchling.com/s/seq-aUFmVRg2Cg7kteXnIai2 |
| Cre recombinase | pCAG-Cre recombinase | https://benchling.com/s/seq-DOwArJA1JjD51YCsVUyd |
| BceINT expression vector | pCAG-NLS-BceINT | https://benchling.com/s/seq-hoKLjTbug98wVgg9cpCM |
| SscINT expression vector | pCAG-NLS-SscINT | https://benchling.com/s/seq-D37aMwZ0Gg5PlwLDtdCK |

| | | |
|---|---|---|
| SacINT expression vector | pCAG-NLS-SacINT | https://benchling.com/s/seq-dVDhPf8bez3pHz1WxLkI |
| HDR sgRNA guide backbone | Minicircle U6-sgRNA EFS-SpCas9 | https://benchling.com/s/seq-2k8wqDuZZC8JY5wD1qGo |
| HDR EGFP cargo | Cas9 HDR template site with EGFP | https://benchling.com/s/seq-K79T50GVDRzDWmZLzxry |
| AAV helper plasmid | PDF6 AAV helper plasmid | https://benchling.com/s/seq-wGjfkT7PBA2YmmgrWuyn |
| AAV EGFP donor | GFP AAV donor plasmid | https://benchling.com/s/seq-BNsdctCG40dNVgOEs4pA |
| AAV2/8 | AAV2/8 capsid protein | https://benchling.com/s/seq-Pk1fGlxnNCAsriHCCBfo |
| Adenovirus EGFP donor | Adenovirus AdEasy-1 backbone with EGFP gene | https://benchling.com/s/seq-HOCw7Wq0ifHTMak47oIA |
| Adenovirus for SpCas9-RT expression | Adenovirus AdEasy-1 backbone pCMV-SpCas9-RT-P2A-Blast | https://benchling.com/s/seq-G9fBVk4IB310dSQ3yF2P |
| Adenovirus for Bxb1 and guide expression | Adenovirus AdEasy-1 backbone pCMV-NLS-Bxb1Blast; pCAG-EGFP; ACTB atgRNA and nicking guide | https://benchling.com/s/seq-1rvrcTnyksAjHncDKCp2 |

**Supplementary Table 7.** Minicircle cargo gene maps

| **Name** | **Full Description** | **Benchling link** |
|---|---|---|
| Cargo EGFP | Parent minicircle plasmid - Cargo EGFP with AttP Bxb1 site | https://benchling.com/s/seq-RksgOMHffn5auc9ZbcHA |
| Cargo EGFP post cleavage | Cargo EGFP with AttP Bxb1 site - post minicircle cleavage | https://benchling.com/s/seq-Na8dODu7S991LQkifkPB |
| Cargo EGFP for fusion | Parent minicircle plasmid - Cargo EGFP with AttP Bxb1 site for fusion | https://benchling.com/s/seq-xdAIg3Ows9xH4RmIv6wP |
| mCherry Cargo post cleavage | Cargo mCherry with AttP Bxb1 site - post minicircle cleavage | https://benchling.com/s/seq-j7JDfHwufSdGxkskf9q8 |
| YFP Cargo post cleavage | Cargo YFP with AttP Bxb1 site - post minicircle cleavage | https://benchling.com/s/seq-161oVqnrJvKvT5Gckzc1 |
| SERPINA1 Cargo post cleavage | Cargo SERPINA1 with AttP Bxb1 site - post minicircle cleavage | https://benchling.com/s/seq-mJmyrX4bGCemUysu4obP |
| CPS1 Cargo post cleavage | Cargo CPS1 with AttP Bxb1 site - post minicircle cleavage | https://benchling.com/s/seq-5VlYu3w2FQUWIfAETQvJ |
| NYESO TCR Cargo post cleavage | Cargo NYESO TCR with AttP Bxb1 site - post minicircle cleavage | https://benchling.com/s/seq-ZgCsKas7TIkxis23XEDq |
| HBB Cargo | Parent minicircle plasmid - Cargo HBB with AttP Bxb1 site | https://benchling.com/s/seq-SjQrU2dc7rCjZ5ktbTVE |

| | | |
|---|---|---|
| Hexa Cargo | Parent minicircle plasmid - Cargo Hexa with AttP Bxb1 site | https://benchling.com/s/seq-JlrqcAgvuRBTvoBrocEM |
| PAH Cargo | Parent minicircle plasmid - Cargo PAH with AttP Bxb1 site | https://benchling.com/s/seq-tmrugQOvFp1TyMDy95tE |
| GBA Cargo | Parent minicircle plasmid - Cargo GBA with AttP Bxb1 site | https://benchling.com/s/seq-sRE7PLmHEA269xToMHyI |
| ADA Cargo | Parent minicircle plasmid - Cargo ADA with AttP Bxb1 site | https://benchling.com/s/seq-1cxKDHbDJisWtqLzWOJW |
| CEP290 Cargo | Parent minicircle plasmid - Cargo CEP290 with AttP Bxb1 site | https://benchling.com/s/seq-nkJagJzDXsN4kNJGYXYq |
| SERPINA1-HIBIT Cargo | Parent minicircle plasmid - Cargo SERPINA1-HIBIT with AttP Bxb1 site | https://benchling.com/s/seq-Gfx0YkqhDaJCr8RjYL1G |
| CPS1-HIBIT Cargo | Parent minicircle plasmid - Cargo CPS1- HIBIT with AttP Bxb1 site | https://benchling.com/s/seq-CnPeB8MAX1V943N47B3q |
| ACTB HITI post cleavage | HITI template for ACTB locus - post minicircle cleavage | https://benchling.com/s/seq-q3ExdFaCoiExkr9cZAvx |
| SUPTH16 HITI post cleavage | HITI template for SUPTH16 locus - post minicircle cleavage | https://benchling.com/s/seq-N3JNZ3DSi7h2YlOo2ELQ |
| SRRM2 HITI post cleavage | HITI template for SRRM2 locus - post minicircle cleavage | https://benchling.com/s/seq-xORs0oLiIQI56GrZc0aJ |
| NOLC1 HITI post cleavage | HITI template for NOLC1 locus - post minicircle | https://benchling.com/s/seq-uSGfnt2OkPS5UCtPJFMj |

| | | |
|---|---|---|
| | cleavage | |
| LMNB1 HITI post cleavage | HITI template for LMNB1 locus - post minicircle cleavage | https://benchling.com/s/seq-jYHbVXXsBKxxcEytXC01 |
| DEPDC4 HITI post cleavage | HITI template for DEPDC4 locus - post minicircle cleavage | https://benchling.com/s/seq-fS7vKseJCDTjJJITfzzd |
| NES HITI post cleavage | HITI template for NES locus - post minicircle cleavage | https://benchling.com/s/seq-Tx6nu4NiQkN7cYzjRujo |

**Supplementary Table 8.** Primers, probes and restriction enzymes used in ddPCR readout

| Locus | Cargo | Forward Primer | Reverse Primer | Probe | Restriction Enzymes |
|---|---|---|---|---|---|
| ACTB | GFP (pDY0186) | CCCGGCTTCCTTTG TCC | GAACTCCACGCCG TTCA | /56-FAM/CC GGC TTG T/ZEN/C GAC GAC GGC G/3IABkFQ/ | Eco91I, HindIII |
| ACTB | TP90-1 GFP (pDY0333) | CCCGGCTTCCTTTG TCC | AACCACAACTAGA ATGCAGTGA | /56-FAM/TG CTA TTG C/ZEN/T TTA TTT GTG GGC CCG /3IABkFQ/ | None |
| ACTB | TP90-1 rc GFP (pDY0334) | CCCGGCTTCCTTTG TCC | GAACTCCACGCCG TTCA | /56-FAM/CC ATG AAG A/ZEN/T CGA GTG CCG CAT CA/3IABkFQ/ | None |
| ACTB | Bt1INT GFP (pDY0367) | CCCGGCTTCCTTTG TCC | AACCACAACTAGA ATGCAGTGA | /56-FAM/TG CTA TTG C/ZEN/T TTA TTT GTG GGC CCG /3IABkFQ/ | None |
| ACTB | Bt1INT rc GFP (pDY0368) | CCCGGCTTCCTTTG TCC | GAACTCCACGCCG TTCA | /56-FAM/CC ATG AAG A/ZEN/T CGA GTG CCG CAT CA/3IABkFQ/ | None |
| LMNB1 | GFP (pDY0186) | TCCTTATCACGGT CCCGCTCG | GAACTCCACGCCG TTCA | /56-FAM/CC ATG AAG A/ZEN/T CGA GTG CCG CAT CA/3IABkFQ/ | Eco91I, HindIII |
| NOLC1 | GFP (pDY0186) | CGTCGACAACGGT AGTG | GAACTCCACGCCG TTCA | /56-FAM/CC ATG AAG A/ZEN/T CGA GTG CCG CAT CA/3IABkFQ/ | Eco91I, HindIII |
| SERPIN1 | GFP (pDY0186) | GGATCCTTGTGAG TGTTGGG | GAACTCCACGCCG TTCA | /56-FAM/CC ATG AAG A/ZEN/T CGA GTG CCG CAT CA/3IABkFQ/ | Eco91I, HindIII |
| SUPT16H | GFP (pDY0186) | TCGCGTGATTCTC GGAAC | GAACTCCACGCCG TTCA | /56-FAM/CC ATG AAG A/ZEN/T CGA GTG CCG CAT CA/3IABkFQ/ | Eco91I, HindIII |
| SRRM2 | GFP (pDY0186) | GGGCGGTAAGTGG TTAGTTT | GAACTCCACGCCG TTCA | /56-FAM/CC ATG AAG A/ZEN/T CGA GTG CCG CAT CA/3IABkFQ/ | Eco91I, HindIII |
| DEPDC4 | GFP (pDY0186) | AAGAGGCGGAGCC AGTA | GAACTCCACGCCG TTCA | /56-FAM/CC ATG AAG A/ZEN/T CGA GTG CCG CAT CA/3IABkFQ/ | Eco91I, HindIII |
| NES | GFP (pDY0186) | CTCCCTTCTCCCGG TGCCC | GAACTCCACGCCG TTCA | /56-FAM/CC GGC TTG T/ZEN/C GAC GAC GGC G/3IABkFQ/ | Eco91I, HindIII |
| ACTB | ACTB HITI template GFP (pDY0219) | CCCGGCTTCCTTTG TCC | GAACTCCACGCCG TTCA | /56-FAM/CC ATG AAG A/ZEN/T CGA GTG CCG CAT CA/3IABkFQ/ | Eco91I |
| SRRM2 | SRRM2 HITI template GFP (aRY0182_A2) | GGGCGGTAAGTGG TTAGTTT | GAACTCCACGCCG TTCA | /56-FAM/CC ATG AAG A/ZEN/T CGA GTG CCG CAT CA/3IABkFQ/ | Eco91I |

| | | | | | |
|---|---|---|---|---|---|
| NOLC1 | NOLC1 HITI template GFP (aRY0182_A3) | CGTCGACAACGGT AGTG | GAACTCCACGCCG TTCA | /56-FAM/CC ATG AAG A/ZEN/T CGA GTG CCG CAT CA/3IABkFQ/ | Eco91I |
| DEPDC4 | DEPDC4 HITI template GFP (aRY0182_A5) | AAGAGGCGGAGCC AGTA | GAACTCCACGCCG TTCA | /56-FAM/CC ATG AAG A/ZEN/T CGA GTG CCG CAT CA/3IABkFQ/ | Eco91I |
| NES | NES HITI template GFP (aRY0182_A7) | CTCCCTTCTCCCGG TGCCC | GAACTCCACGCCG TTCA | /56-FAM/CC ATG AAG A/ZEN/T CGA GTG CCG CAT CA/3IABkFQ/ | Eco91I |
| LMNB1 | LMNB1 HITI template GFP (aRY0182_A4) | TCCTTATCACGGT CCCGCTCG | GAACTCCACGCCG TTCA | /56-FAM/CC ATG AAG A/ZEN/T CGA GTG CCG CAT CA/3IABkFQ/ | Eco91I |
| SUPT16H | SUPT16H HITI template GFP (aRY0182_A1) | TCGCGTGATTCTC GGAAC | GAACTCCACGCCG TTCA | /56-FAM/CC ATG AAG A/ZEN/T CGA GTG CCG CAT CA/3IABkFQ/ | Eco91I |
| ACTB | SERPINA (pDY0298) | CCCGGCTTCCTTTG TCC | GGCCTGCCAGCAG GAGGA | /56-FAM/CC GGC TTG T/ZEN/C GAC GAC GGC G/3IABkFQ/ | EcoRI, XhoI, HindIII |
| ACTB | CPS1 (pDY299) | CCCGGCTTCCTTTG TCC | GGTGTGCAGTCAC ATTGGTAAAGCC | /56-FAM/AC AGC TTT C/ZEN/A AAG TGG TGA GGA CAC T/3IABkFQ/ | XhoI, HindIII |
| ACTB | CFTR (pDY0373) | CCCGGCTTCCTTTG TCC | GATGGGTCTAGTC CAGCTAAAG | /56- FAM/TACGGTACA/ZEN/AACCC ACCCGAGAGA/3IABkFQ/ | Eco91I, HindIII |
| ACTB | NYESO TRAC (pDY0318) | CCCGGCTTCCTTTG TCC | GAGAGACAAGGC TGCACA | /56- FAM/TACGGTACA/ZEN/AACCC ACCCGAGAGA/3IABkFQ/ | Eco47III, HindIII |
| LMNB1 | NYESO TRAC (pDY0318) | TCCTTATCACGGT CCCGCTCG | GAGAGACAAGGC TGCACA | /56- FAM/TACGGTACA/ZEN/AACCC ACCCGAGAGA/3IABkFQ/ | Eco47III, HindIII |
| LMNB1 | SERPINA (pDY0298) | TCCTTATCACGGT CCCGCTCG | GGCCTGCCAGCAG GAGGA | /56-FAM/CC GGC TTG T/ZEN/C GAC GAC GGC G/3IABkFQ/ | EcoRI, XhoI, HindIII |
| NC_000003 | GFP (pDY0186) | CCAGGTGAGAGTC AGGGTAGTGTTCA | GAACTCCACGCCG TTCA | /56-FAM/CC GGC TTG T/ZEN/C GAC GAC GGC G/3IABkFQ/ | Eco91I, HindIII |
| NC_000002 | GFP (pDY0186) | AGGGACCTTTGCC TGTGTGAGTC | GAACTCCACGCCG TTCA | /56-FAM/CC GGC TTG T/ZEN/C GAC GAC GGC G/3IABkFQ/ | Eco91I, HindIII |
| NC_000009 | GFP (pDY0186) | TCAGCTCTGTGCT GAGGCGAA | GAACTCCACGCCG TTCA | /56-FAM/CC GGC TTG T/ZEN/C GAC GAC GGC G/3IABkFQ/ | Eco91I, HindIII |
| chr6:1490459 59 | GFP (pDY0186) | AAGCCATCTCCCA GAATATCTGCTTA GAAATG | GAACTCCACGCCG TTCA | /56-FAM/CC GGC TTG T/ZEN/C GAC GAC GGC G/3IABkFQ/ | Eco91I, HindIII |
| chr16:186077 30 | GFP (pDY0186) | GAGAGGAGCAAC AGTGAGCATGATG | GAACTCCACGCCG TTCA | /56-FAM/CC GGC TTG T/ZEN/C GAC GAC GGC G/3IABkFQ/ | Eco91I, HindIII |
| chr6:1490459 | ACTB HITI template | AAGCCATCTCCCA | GAACTCCACGCCG | /56-FAM/CC GGC TTG T/ZEN/C | Eco91I |

| | | | | | |
|---|---|---|---|---|---|
| 59 | GFP (pDY0219) | GAATATCTGCTTA GAAATG | TTCA | GAC GAC GGC G/3IABkFQ/ | |
| chr16:186077 30 | ACTB HITI template GFP (pDY0219) | GAGAGGAGCAAC AGTGAGCATGATG | GAACTCCACGCCG TTCA | /56-FAM/CC GGC TTG T/ZEN/C GAC GAC GGC G/3IABkFQ/ | Eco91I |
| ACTB | CAG_Kozak_bGH_ther apeutic_genes generic minicircle | CCCGGCTTCCTTTG TCC | GGCTATGAACTAA TGACCCCGT | /56-FAM/CC GGC TTG T/ZEN/C GAC GAC GGC G/3IABkFQ/ | Eco91I, HindIII |
| ACTB | Hibit-SERPINA (pDY0405) | CCCGGCTTCCTTTG TCC | GGCCTGCCAGCAG GAGGA | /56-FAM/CC GGC TTG T/ZEN/C GAC GAC GGC G/3IABkFQ/ | EcoRI, XhoI, HindIII |
| ACTB | Hibit-CPS1 (pDY406) | CCCGGCTTCCTTTGT CC | GGTGTGCAGTCAC ATTGGTAAAGCC | /56-FAM/AC AGC TTT C/ZEN/A AAG TGG TGA GGA CAC T/3IABkFQ/ | XhoI, HindIII |
| LMNB1 | mCherry (pDY0414) | TCCTTATCACGGT CCCGCTCG | cgcatgaactccttgatgatg gcc | /56-FAM/CC GGC TTG T/ZEN/C GAC GAC GGC G/3IABkFQ/ | EheI, HindIII |
| NOLC1 | (YBP pDY0415) | CGTCGACAACGGT AGTG | GGCACCACTCCGG TAAA | /56-FAM/CC GGC TTG T/ZEN/C GAC GAC GGC G/3IABkFQ/ | Eco47I, HindIII |
| ACTB | AAV and Adenovirus Cargo (pDY0611) | CCCGGCTTCCTTTG TCC | GAACTCCACGCCG TTCA | /56-FAM/CC GGC TTG T/ZEN/C GAC GAC GGC G/3IABkFQ/ | PacI, HindIII |
| ACTB | GFP universal, (novel integrases) | CCCGGCTTCCTTTG TCC | GAACTCCACGCCG TTCA | /56-FAM/CC ATG AAG A/ZEN/T CGA GTG CCG CAT CA/3IABkFQ/ | Eco91I, HindIII |

**Supplementary Table 9.** Primers used for NGS readout

| Description | ID | Sequence (5'-3') |
|---|---|---|
| N-term ACTB Tn5 readout F 1 | PD0966 | ACACTCTTTCCCTACACGACGCTCTTCCGATCTCCGACCTCGGCTCACAGCG |
| N-term ACTB Tn5 readout F 2 | PD0967 | ACACTCTTTCCCTACACGACGCTCTTCCGATCTACCGACCTCGGCTCACAGCG |
| N-term ACTB Tn5 readout F 3 | PD0968 | ACACTCTTTCCCTACACGACGCTCTTCCGATCTGACCGACCTCGGCTCACAGCG |
| N-term ACTB Tn5 readout F 4 | PD0969 | ACACTCTTTCCCTACACGACGCTCTTCCGATCTTGACCGACCTCGGCTCACAGCG |
| N-term ACTB Tn5 readout F 5 | PD0970 | ACACTCTTTCCCTACACGACGCTCTTCCGATCTCTGACCGACCTCGGCTCACAGCG |
| N-term ACTB Tn5 readout F 6 | PD0971 | ACACTCTTTCCCTACACGACGCTCTTCCGATCTACTGACCGACCTCGGCTCACAGCG |
| N-term ACTB Tn5 readout F 7 | PD0972 | ACACTCTTTCCCTACACGACGCTCTTCCGATCTTACTGACCGACCTCGGCTCACAGCG |
| N-term ACTB Tn5 readout F 8 | PD0973 | ACACTCTTTCCCTACACGACGCTCTTCCGATCTGTACTGACCGACCTCGGCTCACAGCG |
| ACTB N-term NGS R for Cas14 indels | FP0952 | GTGACTGGAGTTCAGACGTGTGCTCTTCCGATCTCCACCCAGCCAGCTCCC |
| LMNB1 locus NGS F 1 | PD3337 | ACACTCTTTCCCTACACGACGCTCTTCCGATCTCCGCTTCGCCCCCTGCC |
| LMNB1 locus NGS F 2 | PD3338 | ACACTCTTTCCCTACACGACGCTCTTCCGATCTACCGCTTCGCCCCCTGCC |
| LMNB1 locus NGS F 3 | PD3339 | ACACTCTTTCCCTACACGACGCTCTTCCGATCTGACCGCTTCGCCCCCTGCC |
| LMNB1 locus NGS F 4 | PD3340 | ACACTCTTTCCCTACACGACGCTCTTCCGATCTTGACCGCTTCGCCCCCTGCC |
| LMNB1 locus NGS F 5 | PD3341 | ACACTCTTTCCCTACACGACGCTCTTCCGATCTCTGACCGCTTCGCCCCCTGCC |
| LMNB1 locus NGS F 6 | PD3342 | ACACTCTTTCCCTACACGACGCTCTTCCGATCTACTGACCGCTTCGCCCCCTGCC |
| LMNB1 locus NGS F 7 | PD3343 | ACACTCTTTCCCTACACGACGCTCTTCCGATCTTACTGACCGCTTCGCCCCCTGCC |
| LMNB1 locus NGS F 8 | PD3344 | ACACTCTTTCCCTACACGACGCTCTTCCGATCTGTACTGACCGCTTCGCCCCCTGCC |
| LMNB1 locus NGS R | PD3336 | GTGACTGGAGTTCAGACGTGTGCTCTTCCGATCTGGTCATTGAGCTCGCGCAGC |
| NOLC1 NGS F 1 | A9 aD0046 | ACACTCTTTCCCTACACGACGCTCTTCCGATCTCGAGTCGTGCTGCGTCGACAA |
| NOLC1 NGS F 2 | A10 aD0046 | ACACTCTTTCCCTACACGACGCTCTTCCGATCTACGAGTCGTGCTGCGTCGACAA |
| NOLC1 NGS F 3 | A11 aD0046 | ACACTCTTTCCCTACACGACGCTCTTCCGATCTGACGAGTCGTGCTGCGTCGACAA |

| | | |
|---|---|---|
| NOLC1 NGS F 4 | A12 aD0046 | ACACTCTTTCCCTACACGACGCTCTTCCGATCTTGACGAGTCGTGCTGCGTCGACAA |
| NOLC1 NGS R | C3 aD0046 | GTGACTGGAGTTCAGACGTGTGCTCTTCCGATCTTGGCGAACTTATTGGCCACCTCT |
| SUPT16H NGS F 1 | A1 aD0046 | ACACTCTTTCCCTACACGACGCTCTTCCGATCTCCGGGACCTCGCGTGATTCTC |
| SUPT16H NGS F 2 | A2 aD0046 | ACACTCTTTCCCTACACGACGCTCTTCCGATCTACCGGGACCTCGCGTGATTCTC |
| SUPT16H NGS F 3 | A3 aD0046 | ACACTCTTTCCCTACACGACGCTCTTCCGATCTGACCGGGACCTCGCGTGATTCTC |
| SUPT16H NGS F 4 | A4 aD0046 | ACACTCTTTCCCTACACGACGCTCTTCCGATCTTGACCGGGACCTCGCGTGATTCTC |
| SUPT16H NGS R | C1 aD0046 | GTGACTGGAGTTCAGACGTGTGCTCTTCCGATCTCTCACCCGCCAATTGCTGTACA |
| DEPDC4 NGS F 1 | B1 aD0046 | ACACTCTTTCCCTACACGACGCTCTTCCGATCTCAAGAGGCGGAGCCAGTACTTCTC |
| DEPDC4 NGS F 2 | B2 aD0046 | ACACTCTTTCCCTACACGACGCTCTTCCGATCTACAAGAGGCGGAGCCAGTACTTCTC |
| DEPDC4 NGS F 3 | B3 aD0046 | ACACTCTTTCCCTACACGACGCTCTTCCGATCTGACAAGAGGCGGAGCCAGTACTTCTC |
| DEPDC4 NGS F 4 | B4 aD0046 | ACACTCTTTCCCTACACGACGCTCTTCCGATCTTGACAAGAGGCGGAGCCAGTACTTCTC |
| DEPDC4 NGS R | C4 aD0046 | GTGACTGGAGTTCAGACGTGTGCTCTTCCGATCTCCCGGAAGCTCGTTCTGACTGA |
| NES NGS F1 | PD5198 | ACACTCTTTCCCTACACGACGCTCTTCCGATCTCAAGCTCTGCGAGCCGCTC |
| NES NGS F2 | PD5199 | ACACTCTTTCCCTACACGACGCTCTTCCGATCTACAAGCTCTGCGAGCCGCTC |
| NES NGS F3 | PD5200 | ACACTCTTTCCCTACACGACGCTCTTCCGATCTGACAAGCTCTGCGAGCCGCTC |
| NES NGS F4 | PD5201 | ACACTCTTTCCCTACACGACGCTCTTCCGATCTTGACAAGCTCTGCGAGCCGCTC |
| NES NGS R | PD5202 | GTGACTGGAGTTCAGACGTGTGCTCTTCCGATCTCCGCGCTGAGCAGCTCAT |
| CFTR NGS F 1 | PD3500 | ACACTCTTTCCCTACACGACGCTCTTCCGATCTCAGCGGCAGGCACCCAGA |
| CFTR NGS F 2 | PD3501 | ACACTCTTTCCCTACACGACGCTCTTCCGATCTACAGCGGCAGGCACCCAGA |
| CFTR NGS F 3 | PD3502 | ACACTCTTTCCCTACACGACGCTCTTCCGATCTGACAGCGGCAGGCACCCAGA |
| CFTR NGS F 4 | PD3503 | ACACTCTTTCCCTACACGACGCTCTTCCGATCTTGACAGCGGCAGG |

| | | |
|---|---|---|
| | | CACCCAGA |
| CFTR NGS R | PD3504 | GTGACTGGAGTTCAGACGTGTGCTCTTCCGATCTCTTTTCCAGAGGCGACCTCTGC |
| SERPINA1 locus new NGS 1 F 1 | PD3487 | ACACTCTTTCCCTACACGACGCTCTTCCGATCTCTGTGTCTGGGACCACAGAGCATTG |
| SERPINA1 locus new NGS 1 F 2 | PD3488 | ACACTCTTTCCCTACACGACGCTCTTCCGATCTACTGTGTCTGGGACCACAGAGCATTG |
| SERPINA1 locus new NGS 1 F 3 | PD3489 | ACACTCTTTCCCTACACGACGCTCTTCCGATCTGACTGTGTCTGGGACCACAGAGCATTG |
| SERPINA1 locus new NGS 1 F 4 | PD3490 | ACACTCTTTCCCTACACGACGCTCTTCCGATCTTGACTGTGTCTGGGACCACAGAGCATTG |
| SERPINA1 locus new NGS 1 F 5 | PD3491 | ACACTCTTTCCCTACACGACGCTCTTCCGATCTCTGACTGTGTCTGGGACCACAGAGCATTG |
| SERPINA1 locus new NGS 1 F 6 | PD3492 | ACACTCTTTCCCTACACGACGCTCTTCCGATCTACTGACTGTGTCTGGGACCACAGAGCATTG |
| SERPINA1 locus new NGS 1 F 7 | PD3493 | ACACTCTTTCCCTACACGACGCTCTTCCGATCTTACTGACTGTGTCTGGGACCACAGAGCATTG |
| SERPINA1 locus new NGS 1 F 8 | PD3494 | ACACTCTTTCCCTACACGACGCTCTTCCGATCTGTACTGACTGTGTCTGGGACCACAGAGCATTG |
| SERPINA1 locus new NGS 1 R | PD3581 | GTGACTGGAGTTCAGACGTGTGCTCTTCCGATCTccaggaatcgggccagatcagaatc |
| NGS EMX1 Forward 1 | PD0313 | ACACTCTTTCCCTACACGACGCTCTTCCGATCTCCGGTGGCGCATTGCCAC |
| NGS EMX1 Forward 2 | PD0314 | ACACTCTTTCCCTACACGACGCTCTTCCGATCTACCGGTGGCGCATTGCCAC |
| NGS EMX1 Forward 3 | PD0315 | ACACTCTTTCCCTACACGACGCTCTTCCGATCTGACCGGTGGCGCATTGCCAC |
| NGS EMX1 Forward 4 | PD0316 | ACACTCTTTCCCTACACGACGCTCTTCCGATCTTGACCGGTGGCGCATTGCCAC |
| NGS EMX1 Forward 5 | PD0317 | ACACTCTTTCCCTACACGACGCTCTTCCGATCTCTGACCGGTGGCGCATTGCCAC |
| NGS EMX1 Forward 6 | PD0318 | ACACTCTTTCCCTACACGACGCTCTTCCGATCTACTGACCGGTGGCGCATTGCCAC |
| NGS EMX1 Forward 7 | PD0319 | ACACTCTTTCCCTACACGACGCTCTTCCGATCTTACTGACCGGTGGCGCATTGCCAC |
| NGS EMX1 Forward 8 | PD0320 | ACACTCTTTCCCTACACGACGCTCTTCCGATCTGTACTGACCGGTGGCGCATTGCCAC |
| NGS EMX1 Reverse | PD0321 | GTGACTGGAGTTCAGACGTGTGCTCTTCCGATCTCAGAGTCCAGCTTGGGCCCA |

**Supplementary Table 10.** UMI TN5 primers for genome-wide off target integration detection

| | | |
|---|---|---|
| Tn5 off-target EGFP pDY232 minicircle Upstream Round 1 Forward primer | PD3327 | AGGAGCTGCACAGCAACACC |
| Tn5 off-target EGFP minicircle pDY232 NGS F 1 | PD3328 | ACACTCTTTCCCTACACGACGCTCTTCCGATCTCCCTTCGCCAGATCTCGAGCTC |
| Tn5 off-target EGFP minicircle pDY232 NGS F 2 | PD3329 | ACACTCTTTCCCTACACGACGCTCTTCCGATCTACCCTTCGCCAGATCTCGAGCTC |
| Tn5 off-target EGFP minicircle pDY232 NGS F 3 | PD3330 | ACACTCTTTCCCTACACGACGCTCTTCCGATCTGACCCTTCGCCAGATCTCGAGCTC |
| Tn5 off-target EGFP minicircle pDY232 NGS F 4 | PD3331 | ACACTCTTTCCCTACACGACGCTCTTCCGATCTTGACCCTTCGCCAGATCTCGAGCTC |
| Tn5 off-target EGFP minicircle pDY232 NGS F 5 | PD3332 | ACACTCTTTCCCTACACGACGCTCTTCCGATCTCTGACCCTTCGCCAGATCTCGAGCTC |
| Tn5 off-target EGFP minicircle pDY232 NGS F 6 | PD3333 | ACACTCTTTCCCTACACGACGCTCTTCCGATCTACTGACCCTTCGCCAGATCTCGAGCTC |
| Tn5 off-target EGFP minicircle pDY232 NGS F 7 | PD3334 | ACACTCTTTCCCTACACGACGCTCTTCCGATCTTACTGACCCTTCGCCAGATCTCGAGCTC |
| Tn5 off-target EGFP minicircle pDY232 NGS F 8 | PD3335 | ACACTCTTTCCCTACACGACGCTCTTCCGATCTGTACTGACCCTTCGCCAGATCTCGAGCTC |
| nested_outside_U6_F | PD4229 | TGGACTATCATATGCTTACCGTAACTTGAAAGTAT |
| nested_inside_Sp_scaffold_NGS_F1 | PD4225 | ACACTCTTTCCCTACACGACGCTCTTCCGATCTCGTCCGTTATCAACTTGAAAAAGTGGCACC |
| nested_inside_Sp_scaffold_NGS_F2 | PD4226 | ACACTCTTTCCCTACACGACGCTCTTCCGATCTACGTCCGTTATCAACTTGAAAAAGTGGCACC |
| nested_inside_Sp_scaffold_NGS_F3 | PD4227 | ACACTCTTTCCCTACACGACGCTCTTCCGATCTGACGTCCGTTATCAACTTGAAAAAGTGGCACC |
| nested_inside_Sp_scaffold_NGS_F4 | PD4228 | ACACTCTTTCCCTACACGACGCTCTTCCGATCTTGACGTCCGTTATCAACTTGAAAAAGTGGCACC |
| Index primer rev | FP0808 | GTGACTGGAGTTCAGACGTGTGCTC |

**Supplementary Table 11.** Off-target sites

| **Description** | **Sequence (5’-3’)** |
|---|---|
| Cas9_chr6:149045959 | GATATTTTCCCAGCTCACCA |

| | |
|---|---|
| Cas9_chr16:18607730 | TCTATTCTCCCAGCTCCCCA |
| Bxb1_NC_000002 (chr 2) | AGCGGCTTCTGTCTCTGTGAGTGAGCTGGCGGTCTCCGTC |
| Bxb1_NC_000003 (chr 3) | GACTAGCCCACGCTCCGGTTCTGAGCCGCGACGGCGGTCTCCG |
| Bxb1_NC_000009 (chr 9) | CCCAGGGTCCCATGCGCTCCCCGGCCCTGACGGCGGTCTCC |
